# Modeling the long-term dynamics of tropical forests: from leaf traits to whole-tree growth patterns

**DOI:** 10.1101/2020.06.01.128256

**Authors:** Gunnar Petter, Holger Kreft, Yongzhi Ong, Gerhard Zotz, Juliano Sarmento Cabral

**Affiliations:** Biodiversity, Macroecology & Biogeography, University of Göttingen, Germany; Ecoinformatics, Biometrics and Forest Growth, University of Göttingen, Germany; Functional Ecology Group, Institute of Biology and Environmental Sciences, University of Oldenburg, Germany; Smithsonian Tropical Research Institute, Panama City, Republic of Panama

**Keywords:** 3D forest stand model, functional-structural tree model, agent-based model, individual-based model, leaf economic spectrum, leaf traits, canopy structure, canopy dynamics, tropical forest, physiological processes, pipe model theory, tree architecture

## Abstract

Tropical forests are the most diverse terrestrial ecosystems and home to numerous tree species with diverse ecological strategies competing for resources in space and time. Functional traits influence the ecophysiological performance of tree species, yet the relationship between traits and emergent long-term growth pattern is poorly understood. Here, we present a novel 3D forest stand model in which growth patterns of individual trees and forest stands are emergent properties of leaf traits. Individual trees are simulated as 3D functional-structural tree models (FSTMs), considering branches up to the second order and leaf dynamics at a resolution of one m^3^. Each species is characterized by a set of leaf traits that corresponds to a specific position on the leaf economic spectrum and determines light-driven carbon assimilation, respiration and mortality rates. Applying principles of the pipe model theory, these leaf scale-processes are coupled with within-tree carbon allocation, i.e., 3D tree growth emerges from low-level processes. By integrating these FSTMs into a dynamic forest stand model, we go beyond modern stand models to integrate structurally-detailed internal physiological processes with interspecific competition, and interactions with the environment in diverse tree communities. For model calibration and validation, we simultaneously compared a large number of emergent patterns at both the tree and forest levels in a pattern-oriented modeling framework. At the tree level, varying specific leaf area and correlated leaf traits determined the maximum height and age of a tree, as well as its size-dependent growth rate and shade tolerance. Trait variations along the leaf economic spectrum led to a continuous transition from fast-growing, short-lived and shade-intolerant to slow-growing, long-lived and shade-tolerant trees. These emerging patterns resembled well-known functional tree types, indicating a fundamental impact of leaf traits on long-term growth patterns. At the forest level, a large number of patterns taken from lowland Neotropical forests were reproduced, indicating that our forest model simulates structurally realistic forests over long time spans. Our ecophysiological approach improves the understanding of how leaf level processes scale up to the tree and the stand level, and facilitates the development of next-generation forest models for species-rich forests in which tree performance emerges directly from functional traits.

## Introduction

Tropical forests provide multiple social, ecological and economical services, represent the most diverse terrestrial ecosystem, and play an important role in the global carbon cycle (Heywood and Watson 1995, Malhi and Grace 2000, Hassan et al. 2005). In addition to high tree diversity, there are even more plant and animal species that directly or indirectly depend on the resources and habitat provided by trees (Erwin 1988, Nakamura et al. 2017). Almost 9% of all vascular plant species, for instance, live as epiphytes on trees, predominantly in subtropical or tropical regions (Zotz 2013). Ongoing deforestation and potential adverse effects of climate change thus pose a threat to all species associated with tropical forests (Wright 2005). To assess the impact of a changing environment on tropical biodiversity, we thus need a mechanistic understanding of the functioning of these forests, e.g., how they will respond to those changes and how associated species respond to changing forest dynamics.

To predict future changes of tropical forests and to analyze the functioning of these ecosystems, various dynamic ecophysiological models are available that differ substantially in the level of detail and temporal and spatial resolution. Among these models, dynamic global vegetation models focus on large-scale predictions of vegetation dynamics and carbon cycles, but commonly use simplistic forest structures (e.g., Cramer et al. 2001, Purves and Pacala 2008). At small to medium scales (<1 ha to >100 km^2^), forest gap models and forest landscape models simulate forest dynamics and tree species composition (reviewed in Bugmann 2001). Such models represent forest structure in more detail by including stems and crowns of individual trees or cohorts, allowing simulations of within-canopy light attenuation and competition among different species or functional types of trees (Köhler and Huth 1998, Tietjen and Huth 2006). Even finer structural details can be simulated by functional-structural tree models (FSTMs), in which trees are represented in 3D space by interconnected structural and functional units, such as branches, leaves, or reproductive organs (Godin and Sinoquet 2005; Sievänen et al. 2014). These ‘virtual trees’ explicitly integrate complex, mechanistic interactions between tree architecture and physiological processes, for instance, light-dependent within-tree carbon acquisition and allocation at the meristem level in growing trees (Sterck et al. 2005, Fourcaud et al. 2008). FSTMs are therefore suitable tools to study structural tree growth as an emergent property of lower level ecophysiological processes. Furthermore, integrating FSTMs into forest stand models can improve our understanding of the fundamental principles that interlink tree and forest dynamics. In addition, such detailed 3D forest models could be useful for model-based studies of the spatio-temporal dynamics of canopy-dwelling plants and animals (Cabral et al. 2015). However, only few attempts have been made to couple FSTMs with forest stand models, and these studies focused on growth of even-age monocultures over short time periods (Feng et al. 2011, Guillemot et al. 2014). So far, species-rich forest models that are based on FSTMs and that simulates all crucial demographic processes over long time spans, i.e., regeneration, growth and mortality, are not available.

Extending FSTMs to the tropical forest stands is both computationally and conceptually challenging. On the one hand, stand-scale FSTMs are computer-intensive due to their complexity and thus require efficient modeling techniques to keep simulations feasible. On the other hand, tropical forests pose particular challenges due to their high species diversity (ter Steege et al. 2013, Slik et al. 2015). In contrast to temperate forests, for which the low number of well-studied tree species allows calibrating ecophysiological parameters at the species level (e.g., Seidl et al. 2012), alternative approaches are required for species-rich forests where species-specific physiological information is scarce. Individual-based models for tropical forests typically use distinct functional groups aggregating tree species with similar characteristics (e.g., Köhler and Huth 1998; Tietjen and Huth 2006). In the simplest case, only light-demanding pioneers and shade-tolerant climax species are distinguished (Swaine and Whitmore 1988), but a classification into more groups has also been proposed (Gourlet-Fleury et al. 2005; Chazdon et al. 2010). While functional group approaches are useful, they retain a simplification of the continuum from fast growing, short-lived pioneer to slow growing, long-lived shade-tolerant species (Denslow 1987, Wright et al. 2003). Similar trade-offs between growth and mortality occur at the leaf scale (Wright et al. 2004): many leaf traits co-vary strongly and this variation is largely explained by a single principle axis - the leaf economics spectrum (LES). The LES runs from leaves with high photosynthetic capacities and low life spans to leaves with low photosynthetic capacities and long life spans. Hence, a relationship between leaf traits and whole-tree performance can be assumed, and significant relationships were observed for many tropical tree species (Sterck et al. 2006, Poorter and Bongers 2006). A leaf trait-based approach should thus be a promising way to integrate the different life history strategies of trees into forest models. However, we are not aware of any study in which 3D growth over a tree’s entire life span has been modelled as an emergent property of the tree’s set of traits.

Here, we present a dynamic forest stand model in which trees are represented as 3D functionally- and structurally-explicit individuals. This model simulates the long-term forest dynamics (500-1000 years) at the plot scale (∼1 ha) with a high level of structural detail and functional diversity. Branches are considered up to the second order and leaf biomass development is modelled at a resolution of 1 m^3^, which allows detailed consideration of competition for light and space. Tree species are characterized by a set of leaf traits under consideration of the trade-offs and correlations between traits (LES; Wright et al. 2004). Using the principles of the pipe model theory (Shinozaki et al. 1964), the light-driven carbon assimilation at the leaf level and the within-tree carbon allocation are coupled. We hypothesize that this continuous, trait-based approach captures essential life history variations across species with regard to their growth, survival, and light demand. In addition, we hypothesize that the long-term dynamics of diverse, tropical forest communities can be reproduced by coupling the FSTM with a forest stand model, in which key demographic processes and between-tree competition are simulated (e.g., competition for light and space, neutral recruitment). To test these hypotheses, we contrast emerging growth patterns at the tree level (e.g., diameter growth rates, maximum height and life span) against observations. In addition, we simulate forest stands from bare soil over 500 years and test if the model can successfully reproduce multiple attributes of Neotropical forests. Specifically, we test if a dynamic equilibrium is reached and if 12 forest attributes (e.g., basal area, net primary production, mortality rate) and several ecological patterns (e.g., diameter and height distribution, vertical leaf area distribution) resemble field observations. Findings from this exercise indicate how trait trade-offs at the leaf level are functionally linked with patterns of tree growth and forest stand dynamics. Furthermore, by providing 3D forest structure and dynamics, our ecophysiological approach can form the basis for future studies of forest and canopy ecology.

## Methods

The model developed in this study generates 3D structural growth of individual trees and (long-term) dynamics of forest stands at the plot level. We term this model a functional-structural forest model (FSFM). This ecophysiological model was implemented using the open-source 3D modeling platform GroIMP (Growth Grammar Interactive Modeling Platform; available at www.grogra.de). In GroIMP, relational growth grammars are implemented in the programming language XL, which is a graph-based extension of the Lindenmayer-Systems (L-Systems), a formal language for the description of plant structure (Lindenmayer 1968a, 1968b). The FSFM code is available at https://github.com/julianoscabral/MoF3D

### Model description

The model description follows the ODD protocol for agent-based models (**O**verview, **D**esign concepts, **D**etails; see Grimm et al. 2006, 2010). The first block of the protocol (**O**verview – subdivided into the elements ’Purpose’, ’State variables and scales’ and ’Process overview and scheduling’) is presented here. The full description including all ecophysiological equations and parameters is available in Appendix S1.

#### Purpose

The FSFM serves two main purposes. First, to study the relationship between leaf trait trade-offs and life-history variation in trees. Ontogenetic growth patterns, maximum height, and life span, as well as the light-dependent growth behavior in our model emerge from continuous plant traits. Second, to generate long-term dynamics of forest stands at a high level of structural detail. By combining the leaf trait-based tree growth with local population and community dynamics, emergent patterns can be evaluated with field observations at the forest level. Our ecophysiological approach to forest dynamics was designed to increase the understanding of bottom-up mechanisms controlling forest dynamics, and, in addition, to be useful for follow-up studies requiring a detailed 3D representation of forest structure and dynamics.

#### Entities, state variables and scales

The FSFM simulates establishment, growth, and mortality of virtual 3D trees at the plot level. The spatial and temporal scale of the forest plot can be defined during experimental design. Here, we simulate forest stands between 0.25 and 1 hectare over 500 to 1000 years in annual time steps. The vertical spatial dimension is associated with typical maximum tree heights (ca. 50-60 m). The 3D space is divided into a regular grid of cubic voxels with a side length of 1 m (Fig. 1a), which defines the spatial resolution of both light and leaf area/biomass distribution. Light is the main driver of vegetative growth and is calculated for all voxels based on the 3D distribution of leaf area.

**Fig. 1.**
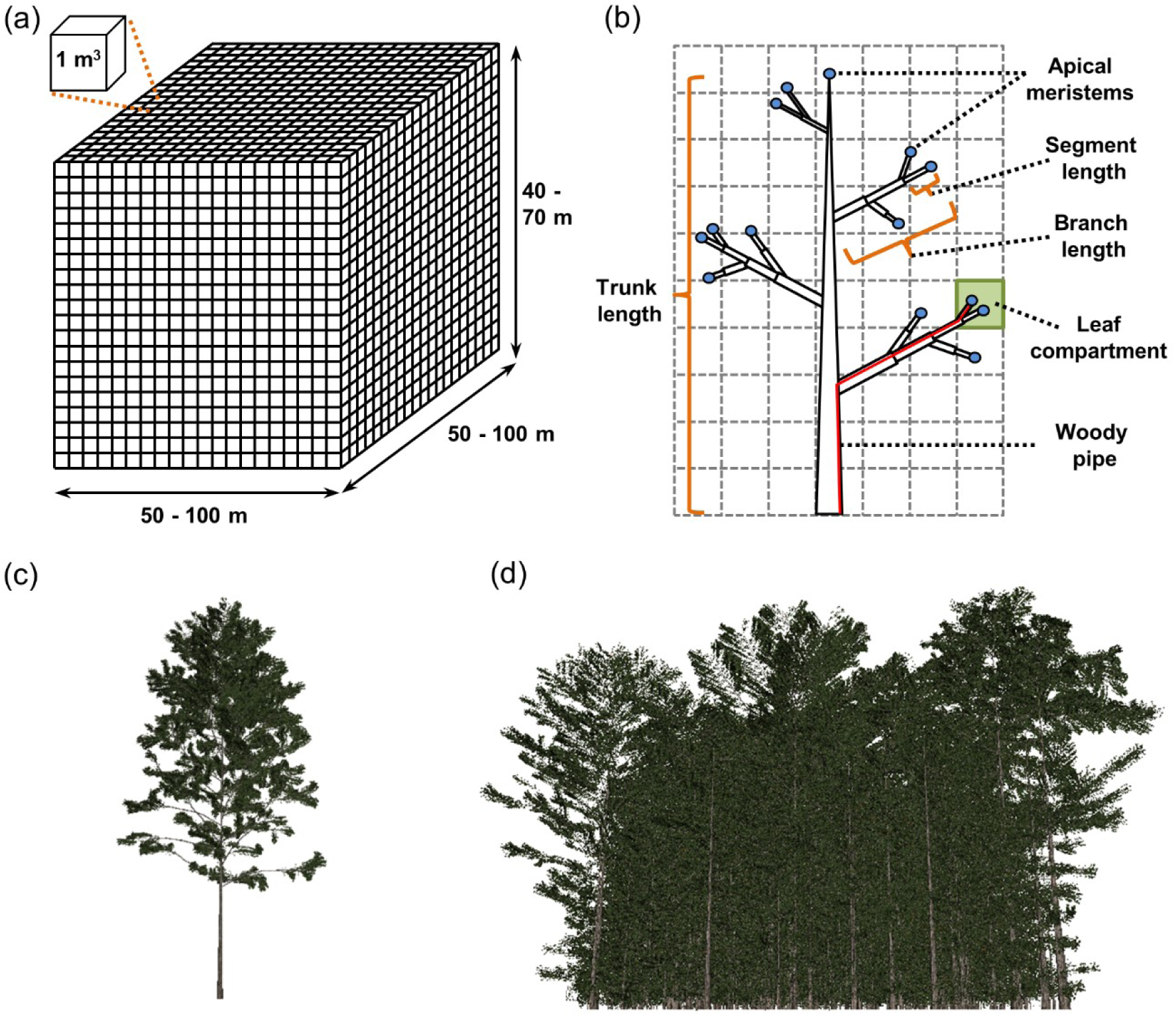
State variables, scales and visualization. (a) 3D model space. The spatiotemporal extent can be defined during experimental design. The model space is a 3D grid that is subdivided into cubic voxels with a volume of 1 m^3^ containing the information about local leaf biomass and area, as well as, light intensity. (b) Overview of tree components: trunks, branches, apical meristems, and leaf compartments. Trees consist of a trunk and branches up to the second order, which are terminated by an apical meristem. Leaf compartments describe the leaf biomass and area within a voxel attached to a specific section of a second order branch, as well as the woody pipes connected to these leaves. The length of the pipe system depends on the within-tree position. One leaf compartment (green square) and its woody pipe (red line) are exemplified. (c) 3D tree visualization (visualization options in Appendix S1). Here, the leaf biomass in the leaf compartments is displayed by spatial objects imitating ‘real’ leaves. (d) 3D forest visualization. The forest structure can be displayed, which allows visual inspections and comparisons with real forests.

This model comprises three hierarchical levels: tree components, individual trees, and the forest stand. Tree components are trunks, branches, apical meristems, and leaf compartments (Fig. 1b). Each tree consists of one erect trunk described by length and diameter. Attached to the trunk are branches up to the second order. Branches are defined at two different scales. At the coarse scale, branches are described by their total length and diameter, while at the fine scale branches are described as topologically connected smaller segments. This multi-scale approach optimizes both speed of the ecophysiological simulations and visual aspects (see Appendix S1 for more details). Located at the end of each trunk or branch, apical meristems sense the local environment and control primary growth. Leaf compartments are connected with second order branches and are conceptualized as aggregations of leaves within the cubic voxels. Leaf compartments comprise leaves and active pipes, which represent the sapwood connecting leaves and roots. This means that leaf compartments form leaf-pipe elements following the pipe model theory (Shinozaki et al. 1964; Fig 1b). Each tree component is characterized by state variables (Table 1), including its absolute 3D position and its topological position within its tree. Based on this information, the 3D structure of each tree can be deduced (Fig. 1c). Structural tree growth results from creation, loss, and dynamic changes of tree components. Structural tree growth is driven by the 3D distribution of light within the forest stand and the functional and the structural traits of trees (Table 1). While the functional traits regulate light-mediated carbon balance, the structural traits reflect inherent architectural parameters defining the tree’s structural organization. This includes, for instance, branching angles or average internode lengths (see Appendix S1 submodel *Structural growth* for more details). Some functional trait combinations promote effective carbon assimilation under low light conditions, allowing growth and survival in the dark understory. Other trait combinations may be more favorable under high light conditions. Consequently, forest dynamics results from structural growth of individual trees with different traits competing for space and light, whose distribution, in turn, is influenced by the forest structure (Fig. 1d).

**Table 1.** State variables, functional and structural traits of the forest model. Each tree component (trunk, branch, leaf compartment, apical meristem) is characterized by a set of state variables. The functional and structural traits describe the intrinsic properties of each tree species. Empirical correlations between leaf functional traits (Wright et al. 2004) are considered (see Appendix S1: Table S2 for more details).

| Symbol | Description | Unit | Type |
| --- | --- | --- | --- |
| $A_B$ | Cross-sectional area of branch | $\text{cm}^2$ | State variable |
| $A_L$ | Leaf area in leaf compartment | $\text{cm}^2$ | State variable |
| $A_{L\text{Prod}}$ | Total leaf area produced in leaf compartment | $\text{cm}^2$ | State variable |
| $A_S$ | Cross-sectional area of branch segment | $\text{cm}^2$ | State variable |
| $A_T$ | Cross-sectional area of trunk | $\text{cm}^2$ | State variable |
| $B_L$ | Leaf biomass in leaf compartment | g | State variable |
| $D_B$ | Diameter of branch | cm | State variable |
| $D_S$ | Diameter of branch segment | cm | State variable |
| $D_T$ | Diameter of trunk | cm | State variable |
| $I_M$ | Light conditions at apical meristem | $\mu\text{mol m}^{-2} \text{s}^{-1}$ | State variable |
| $L_B$ | Length of branch | cm | State variable |
| $L_P$ | Pipe length of leaf compartment (corrected after apical control) | cm | State variable |
| $L_{PS}$ | Pipe length of leaf compartment | cm | State variable |
| $L_S$ | Length of branch segment | cm | State variable |
| $L_T$ | Length of trunk | cm | State variable |
| $O_B$ | Branch order | - | State variable |
| $P_{B\text{End}}^{\text{XYZ}}$ | End position of branch (in X, Y and Z direction) | cm | State variable |
| $P_{B\text{Start}}^{\text{XYZ}}$ | Start position of branch (in X, Y and Z direction) | cm | State variable |
| $P_{LC}^{\text{XYZ}}$ | Position of leaf compartment (in X, Y and Z direction) | cm | State variable |
| $P_M^{\text{XYZ}}$ | Position of apical meristem (in X, Y and Z direction) | cm | State variable |
| $P_{S\text{End}}^{\text{XYZ}}$ | End position of branch segment (in X, Y and Z direction) | cm | State variable |
| $P_{S\text{Start}}^{\text{XYZ}}$ | Start position of branch segment (in X, Y and Z direction) | cm | State variable |
| $P_T^{\text{XY}}$ | Position of trunk (in X and Y direction) | cm | State variable |
| $G_{\text{max}}$ | Maximum gross photosynthetic rate | $\text{g g}^{-1} \text{d}^{-1}$ | Functional trait |
| $k$ | Light intensity at which the gross photosynthetic rate is half maximal | $\mu\text{mol m}^{-2} \text{s}^{-1}$ | Functional trait |
| $LL$ | Leaf lifespan | d | Functional trait |
| $N_{\text{mass}}$ | Nitrogen concentration | % | Functional trait |
| $R_L$ | Leaf respiration rate | $\text{g g}^{-1} \text{d}^{-1}$ | Functional trait |
| $SLA$ | Specific leaf area | $\text{cm}^2 \text{g}^{-1}$ | Functional trait |
| $\rho_w$ | Wood density | $\text{g cm}^{-3}$ | Functional trait |
| $A_{L\text{Prod}}$ | Maximal leaf area production per leaf compartment | $\text{cm}^2$ | Structural trait |
| $I_T$ | Light intensity threshold regulating apical dominance of SAM | $\mu\text{mol m}^{-2} \text{s}^{-1}$ | Structural trait |
| $k_{\text{int}}$ | Factor controlling the increase in internode length | - | Structural trait |
| $LD_B$ | Length-diameter ratio of branches | $\text{m cm}^{-1}$ | Structural trait |
| $LD_T$ | Length-diameter ratio of trunks | $\text{m cm}^{-1}$ | Structural trait |
| $L_{IBMax}$ | Maximal internode length of branches | cm | Structural trait |
| $L_{IBMin}$ | Minimal internode length of branches | cm | Structural trait |
| $L_{ITMax}$ | Maximal internode length of trunk | cm | Structural trait |
| $L_{ITMin}$ | Minimal internode length of trunk | cm | Structural trait |
| $PH_{FO}$ | Number of first order branches arranged in a 360° circle | - | Structural trait |
| $P_{RU}$ | Pipe-reuse factor | - | Structural trait |
| $S_F$ | Shortening factor | - | Structural trait |
| $S_{Trop}$ | Strength of tropism | - | Structural trait |
| $ST_{Trop}$ | Stochasticity in tropism strength ($S_{Trop}$); only used if *Stochasticity*=1 | % | Structural trait |
| $ST_{Tw}$ | Stochasticity in branch growth; only used if *Stochasticity*=1 | % | Structural trait |
| $ST_{aSFO}$ | Stochasticity in second order angle ($\alpha_{SFO}$); only used if *Stochasticity*=1 | % | Structural trait |
| $ST_{aTFO}$ | Stochasticity in first order angle ($\alpha_{TFO}$); only used if *Stochasticity*=1 | % | Structural trait |
| $ST_{aTSO}$ | Stochasticity in first order angle ($\alpha_{TSO}$); only used if *Stochasticity*=1 | % | Structural trait |
| $\alpha_{SFO}$ | Angle between first order branches and trunk from side view | ° | Structural trait |
| $\alpha_{TFO}$ | Angle between first order branches from top view | ° | Structural trait |
| $\alpha_{TSO}$ | Angle between second order branches and first order branches from top view | ° | Structural trait |
| $\beta_D$ | Maximum relative increase in height growth when the $I_M < I_T$ | - | Structural trait |
| $\beta_S$ | Shape parameter regulating apical dominance of trunk apical meristem | - | Structural trait |

#### Initialization

At the beginning of each simulation, a pool of virtual species is generated with the number of species defined during experimental design. Each virtual species is characterized by a set of functional and structural traits (all species-specific traits are listed in Table 1). While the structural traits are uncorrelated and randomly selected from ranges defined during experimental design, functional leaf traits integrate between-trait correlations following the LES (Wright et al. 2004; Marino, Aqil and Shipley 2010). The LES quantifies relationships between leaf economic traits, such as the SLA, leaf lifespan or mass-based photosynthetic capacity. These leaf traits co-vary strongly and, in multidimensional trait space, the vast majority of variation is explained by a single principle axis (Wright et al. 2004). This axis can be considered as a spectrum, ranging from leaves with low SLA values, low photosynthetic capacities, and respiration rates, but long lifespans, to leaves with high SLA values, high photosynthetic capacities, and respiration rates, but short lifespans. To obtain a set of leaf traits, the values for SLA are randomly selected from ranges defined during experimentla design, and the values of the other traits are determined based on between-trait correlations, meaning that each set of leaf traits thus represents a position on the LES. Furthermore, functional traits characterizing specific photosynthetic light-response curves are also predicted from the leaf traits of the LES (following Marino, Aqil and Shipley 2010; Appendix S1: Table S2).

#### Process overview and scheduling

After initialization, light distribution, tree establishment, tree growth, and tree mortality are simulated successively in annual time steps (Fig. 2). The 3D distribution of light intensity is calculated via the Lambert-Beer light extinction law considering the distribution of leaf area. Subsequently, establishment of tree seedlings is simulated as a neutral process. Depending on an average user-defined neutral germination rate (in number of seedlings per ha), new seedlings are initialized at random positions. Each seedling is randomly assigned to a species from the species pool. Seedlings with unsuitable traits for local voxel conditions may die due to carbon starvation. Tree growth is simulated in three subsequent sub-processes: (I) apical control/dominance, (II) carbon balance, (III) structural growth (Fig. 2).

**Fig. 2.**
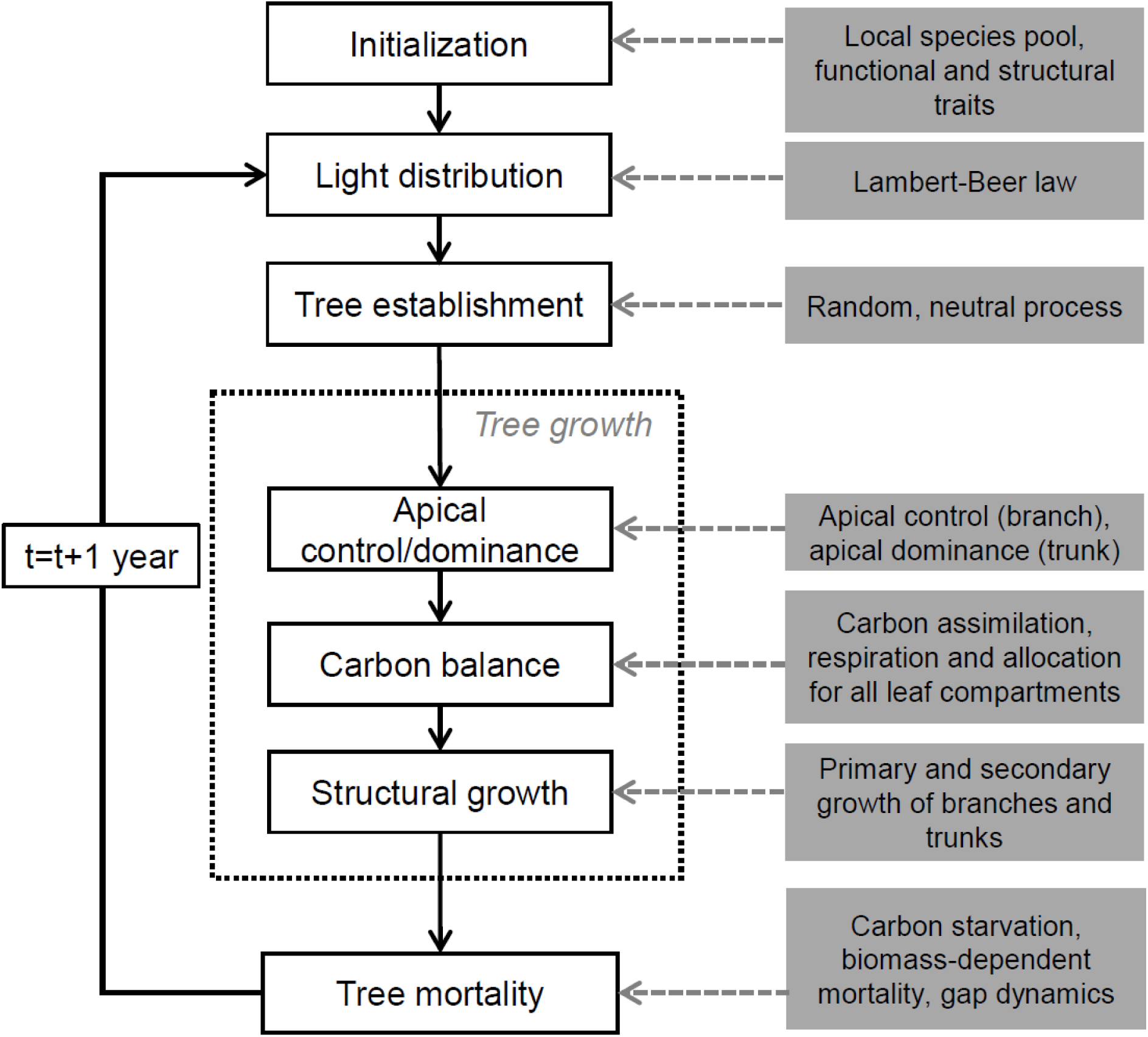
Flowchart of the ecophysiological framework of the model with integrated mechanisms. After initialization, light distribution, tree establishment, growth and mortality take place consecutively in annual time steps. Tree growth is the most complex process and thus split into three subprocesses: apical control, carbon balance and structural growth (details in Appendix S1).

(I) Controlled by plant hormones, carbon allocation to apical meristems can either be inhibited (apical control) or intensified (apical dominance; Wilson 2000). These processes control how much of the carbon assimilated by photosynthesis is invested into primary growth of branches and the trunk. In this model, apical control is considered for branches. Branches inhibit carbon allocation to primary growth when branch apical meristems are 1) deeply shaded, i.e., if the carbon balance under local light conditions at the meristem is negative, or 2) when branches from neighboring trees grow into the same voxel. Hence, light and space competition takes place at the branch level. Apical dominance, in contrast, is considered at the trunk level, i.e., carbon allocation to trunk apical meristems is intensified under shade as a mechanism to quickly reach higher, potentially less shaded zones (Poorter 1999; Poorter et al. 2011). By influencing the within-tree carbon allocation, the processes of apical dominance/control slightly affect local carbon balance, the next tree growth subprocess.

(II) Local carbon balance is simulated at the level of leaf compartments. Apart from the usually only small percentage of carbon assimilated by leaf compartment and allocated to primary growth (influenced by apical control/dominance), leaf compartments are assumed to be independent from each other. Here, we follow the physiological principles of module autonomy stating that different tree parts may be regarded as autonomous modules (Sprugel et al. 1991). This means that no carbon flow occurs between leaf compartments and thus assimilated carbon is locally reinvested. Local re-investment means investments in leaf biomass within the leaf compartment, which, however, are coupled with investments in connected woody pipes following the pipe model theory (Shinozaki et al. 1964). For each new unit of leaf biomass an equivalent unit of pipes connecting leaves and roots has to be grown. Both construction and respiration costs for leaves and pipes are considered, which for the latter depend on the within-tree position of the leaf compartment. The carbon balance is central for the next and final tree growth sub-process – structural growth.

(III) In the structural growth sub-process, secondary and primary growth of branches and trunks are calculated. The sum of new pipes over all leaf compartments connected to a branch yields the branch diameter increment, and the sum over all leaf compartments of a tree yields the trunk diameter increment. Primary growth, in turn, is calculated based on the secondary growth using species-specific allometric relationships between diameter and length/height. Primary growth causes the establishment of new apical branch segments and sometimes new lateral branch segments, which may be associated with new leaf compartments and apical meristems. In addition, trees may also shed branches if all photosynthetically active leaf compartments are lost. After tree growth, tree mortality is simulated via different ecophysiological and environmental processes. Trees may die due to carbon starvation when they have lost all leaves. In addition, we integrated a biomass-dependent mortality rate according to the metabolic theory of ecology (Brown et al. 2004). This rate accounts for processes not explicitly simulated (e.g., herbivory, pathogens) and assumes that the chance of survival increases non-linearly with tree biomass. Furthermore, gap dynamics are also important in tropical forests (Brokaw 1985). Therefore, falling dead trees may kill neighboring trees and create gaps. After each of the processes illustrated in Fig. 2, the state variables of all tree components are updated synchronously. For more details see Appendix S1.

### Outputs

The updated state variables can be saved as model output in annual time steps at all hierarchical levels (tree components, individual trees, forest stand). Output data at the lowest hierarchical level contain the state variables including spatial coordinates. From these variables, outputs at higher hierarchical levels are deduced. For instance, the crown dimensions of a tree result from size and position of its trunk and branch segments. Likewise, forest attributes (e.g., aboveground biomass or the leaf area index) are calculated by aggregating all relevant components of all trees. A list of important outputs is given in Fig. 3; an exhaustive list is available in Appendix S1: Table S5. In addition, data on the 3D light distribution, tree mortality, and species traits may be monitored. Visualization options are described in Appendix S1.

**Fig. 3.**
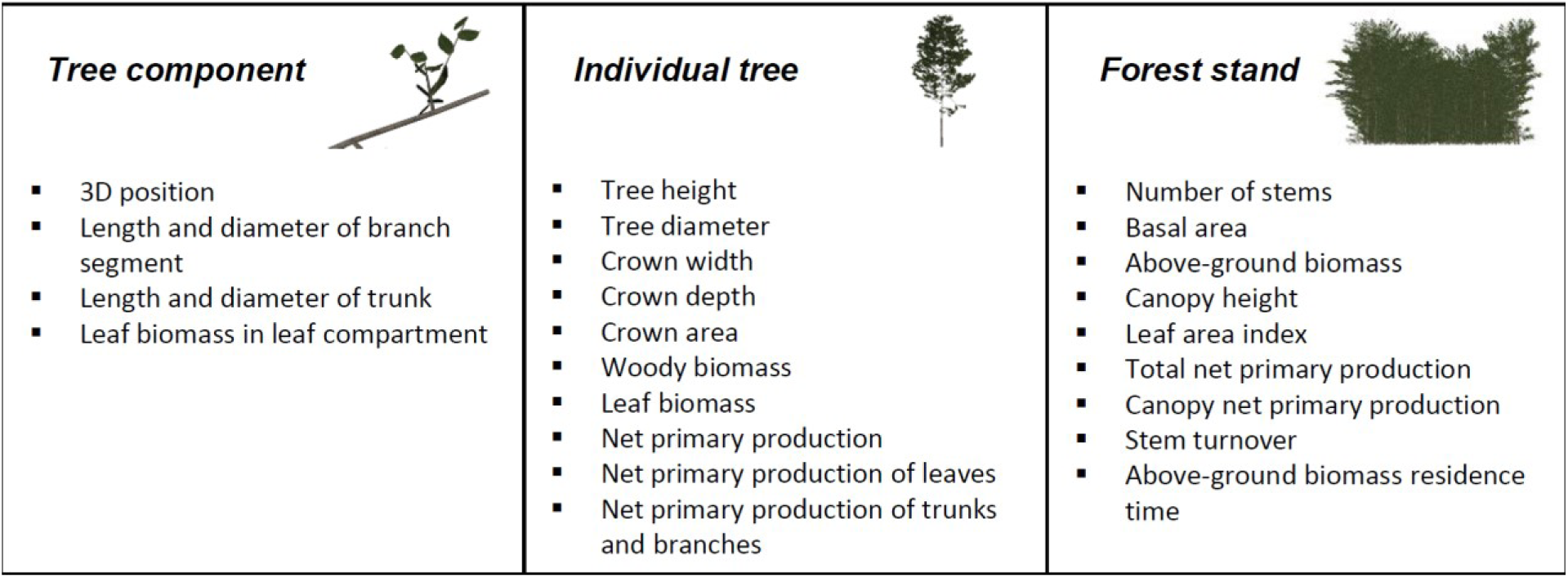
Example of emergent results from our ecophysiological approach at the three hierarchical scales: tree component, individual tree and forest stand. A complete list of all emergent variables is provided in Appendix S1: Table S5.

### Experimental design: model parameterization and validation

Our experimental design aimed at finding values of model parameters that yield realistic reproductions of tree and forest dynamics. Model parameters can be grouped into three categories: (I) global parameters, (II) functional and structural traits, and (III) external environmental drivers (see Appendix S1: Tables S6 and S8 for full list of parameters). We first defined values for those global parameters whose uncertainty, according to literature data, was low. For the functional and structural traits, value ranges instead of single values were identified, as simulated forest communities with a reasonable trait space should reflect the trait space of a species-rich forest. As the natural distribution of functional traits used in this study (i.e., SLA and wood density) is well-studied (e.g., Baker et al. 2004; Patiño et al. 2012), and structural traits are easy-to-interpret characteristics of tree structure, such as branching angles or internode lengths, reasonable trait spaces could be retrieved from the literature. Nevertheless, 12 parameters with uncertain values remained, which were thus treated as free parameters to be calibrated (Table 2). These free parameters were either global parameters or external environmental drivers (i.e., light conditions, site index), and for most of these parameters, ecologically meaningful value ranges could be identified (Table 2).

**Table 2.** Free parameters varied during model parameterization. Value ranges were determined based on literature values or own estimates, and calibrated values are given in brackets. A complete list of all parameter values used in the model is given in Appendix S1: Table S6.

| Parameter<br>(Symbol) | Unit | Explanation | Type | Value range<br>(Fitted value) | References |
| --- | --- | --- | --- | --- | --- |
| $A_{LMax}$ | $cm^2$ | Maximum leaf area per $m^3$ leaf compartment | Global parameter | 10,000-20,000 (15,000) | Own estimate |
| $A_{LProdMax}$ | $cm^2$ | Absolute maximum leaf area production per $m^3$ leaf compartment | Global parameter | 40,000-80,000 (65,000) | Own estimate |
| $I_{max}$ | $\mu mol\ m^{-2}\ s^{-1}$ | Average annual light intensity above canopy during the day | Environmental driver | 700-1200 (900) | Chazdon and Fetcher 1984; Berry, Varney and Flanagan 1997; Valladares, Allen and Pearcy 1997; Sterck et al. 2011; Seyoum et al. 2014 |
| $k_L$ | - | Light extinction coefficient of Beer-Lambert equation | Global parameter | 0.5-0.8 (0.6) | Huth and Ditzer 2000; Kitajima, Mulkey and Wright 2005b; Malhi et al. 2013 |
| $LP_{ratio}$ | $cm^2\ cm^{-2}$ | Ratio between leaf area and pipe cross-sectional area | Global parameter | 5,000-50,000 (40,000) | Wright et al. 2006; Calvo-Alvarado, McDowell and Waring 2008; Patiño et al. 2012 |
| $m_{Neigh}$ | $a^{-1}$ | Probability of a tree dying within reach of a large falling tree | Global parameter | 0.05-0.3 (0.15) | Own estimate |
| $m_{TB}$ | $g^{-1}\ a^{-1}$ | Parameter of biomass-based tree mortality rate | Global parameter | 0.01-0.05 (0.032) | Own estimate |
| $M_{TS}$ | - | Scaling exponent of biomass-based tree mortality rate | Global parameter | 0.1-0.4 (0.13) | Brown et al. 2004; Muller-Landau et al. 2006a,b |
| $n_{Seed}$ | $ha^{-1}\ a^{-1}$ | Number of new seedlings per ha and year | Global parameter | 100-1,000 (500) | Own estimate |
| $P_{RU}$ | - | Pipe-reuse factor, specifies the ratio of new pipes to reused old pipes when new leaves are generated | Global parameter | 0.5-0.9 (0.6) | Baker et al. 2004; Patiño et al. 2009; Quesada et al. 2012; Iida et al. 2014 |
| $R_w$ | $gC\ g^{-1}\ d^{-1}$ | Respiration rate of sapwood | Global parameter | $10^{-4}$-$10^{-6}$ ($5*10^{-4}$) | Penning De Vries 1975; Ryan et al. 1994, 1995; Vose and Ryan 2002; Sterck et al. 2005 |
| $SI$ | - | Site index describing the relative quality of the forest patch | Environmental driver | 0.5-1 (0.9) | Own estimate |

We used key ideas of the pattern-oriented modeling (POM) framework to parameterize and validate our model (Grimm et al. 2005; Grimm and Railsback 2012). The rationale behind POM is to reduce the uncertainty in model structure and parameters by comparing model results with multiple ecological patterns at different hierarchical levels and scales (Grimm et al. 2005). The more patterns reproduced by a model, the higher the likelihood that integrated bottom-up mechanisms are robust and ecologically meaningful. The hierarchical structure of our approach, linking tree components, individual trees and forest stands, suits the POM framework perfectly. Accordingly, we included a large number of patterns at the tree and forest stand level, and tested if these patterns can be reproduced simultaneously. We opted to calibrate the model manually for the following two reasons. On the one hand, we included some qualitative patterns that require visual inspection. On the other hand, we had no access to cluster computers and a full-factorial calibration or advanced parameter estimation techniques (e.g., Bayesian calibration) were not feasible. The manual calibration started by varying each free parameter over its estimated range, while keeping the other parameters constant at mean values, to assess the influence of the parameter on the ecological patterns as described below. Based on these results, the parameters were fine-tuned by recursively adjusting the parameter values and comparing the resulting patterns, using the information obtained in each step in the next step. This recursive process was repeated until we found an appropriate combination of parameter values that reproduced the ecological pattern at the tree and forest level simultaneously. In the following, these patterns are described.

#### Tree level

At the tree level, we evaluated simulated growth of individual trees. For each parameter combination, a visual analysis of ontogenetic tree growth trajectories in combination with an analysis of the changes in height and diameter, biomass, and productivity during tree ontogeny served as first indicator for the structural realism of 3D tree growth. Furthermore, for each unique combination of global parameters, we systematically analyzed how changes in species traits and environmental conditions influence tree growth. To cover the trait and environmental space in tropical forests, we varied each of these factors within their natural ranges (Table 2) while keeping the other factors constant at intermediate levels. Due to the size and longevity of trees, controlled experiments on changes in growth rates and morphology over a tree’s lifespan are practically impossible, and consequently data equivalent to our simulation experiment is missing. Nevertheless, based on numerous field and theoretical studies, there is an understanding of several qualitative and quantitative patterns during tree ontogeny. For instance, while the height growth of undisturbed trees is expected to continue at decreasing rates until reaching maximum height, diameter growth rates tend to peak at a certain height or age (Clark and Clark 1999). In addition, diameter growth rates, maximum tree heights and partly also maximum tree ages are well-studied (Martínez-Ramos and Alvarez-Buylla 1998, Clark and Clark 2001, Chao et al. 2008).

#### Stand level

At the stand level, we evaluated model performance in reproducing the general structure and dynamics of tropical forests. Comparative studies have shown substantial differences in the characteristics of tropical forests between continents (e.g., Feldpausch et al. 2011), and in this study, we focused on the well-studied Neotropical lowland forests. While typical ranges of attributes of mature lowland forests, such as the basal area or net primary production per hectare, are relatively well-known, long-term data to which the model could be fitted remain scarce. Considering this limitation, we estimated ranges of 12 important attributes characterizing both forest structure and dynamics based on an extensive literature review (see Table 3, and Appendix S1: Table S7 for more details). We excluded data from climatically extreme or recently disturbed sites; the estimated ranges should be representative for average lowland forests in dynamic equilibrium state. We assessed model performance by simulating 1 ha forest plots starting from bare soil for 500 years, and calculated the model performance criterion α_M_ that tests if the attributes of the simulated forest in dynamic equilibrium state are within the empirical ranges:

**Table 3.** Typical ranges of forest attributes in Neotropical lowland forests derived from a literature review. We concentrated on reviews covering multiple forest plots or larger forest areas. More details are given in Appendix S1: Table S7.

| Forest attribute | Unit | Typical range | References |
| --- | --- | --- | --- |
| Stem density (> 10cm dbh) | ha<sup>-1</sup> | 480-620 | Feldpausch et al. 2011, Banin et al. 2012 |
| Stem density | ha<sup>-1</sup> | 2500-5000 | Girardin et al. 2013, Chave et al. 2003, DeWalt and Chave 2004 |
| Basal area | m<sup>2</sup> ha<sup>-1</sup> | 25-35 | Feldpausch et al. 2011, Banin et al. 2012, Girardin et al. 2013, Malhi et al. 2006, Baker et al. 2004 |
| Above-ground biomass | Mg ha<sup>-1</sup> | 250-350 | Malhi et al. 2006, Feldpausch et al. 2012, Baker et al. 2004 |
| Canopy height | m | 27-38 | Feldpausch et al. 2011, Asner et al. 2013, Girardin et al. 2013 |
| Mean dbh (> 10cm dbh) | m | 0.18-0.26 | Lieberman et al. 1996, Banin et al. 2014, Sawada et al. 2015 |
| Leaf area index | m m<sup>-2</sup> | 3.5-5.5 | Myneni et al. 2007, Doughty and Goulden 2008, Caldararu, Palmer and Purves 2012 |
| Total net primary production | Mg ha<sup>-1</sup> a<sup>-1</sup> | 10-20 | Malhi, Doughty and Galbraith 2011, Aragão et al. 2009, Malhi et al. 2013 |
| Canopy net primary production | Mg ha<sup>-1</sup> a<sup>-1</sup> | 5-10 | Chave et al. 2010, Malhi et al. 2013, Aragão et al. 2009 |
| Basal area growth | m<sup>2</sup> ha<sup>-1</sup> a<sup>-1</sup> | 0.5-1.0 | Malhi et al. 2004, Lewis et al. 2004b, Banin et al. 2014 |
| Above-ground biomass residence time | a | 25-60 | Malhi et al. 2011, 2015, Galbraith et al. 2014 |
| Stem turnover (> 10cm dbh) | a<sup>-1</sup> | 1-4 | Quesada et al. 2012, Lewis et al. 2004, Malhi et al. 2009 |

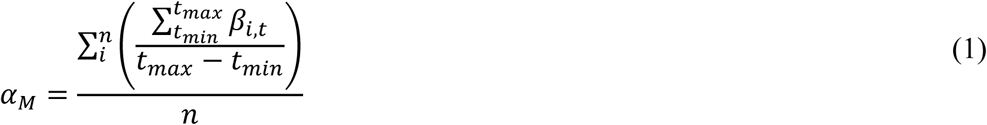

where *t*_min_ is the time after which an equilibrium state is expected (here, *t*_min_=200 years), *t*_max_ is the total number of years simulated (here, *t*_max_=500), *n* is the total number of forest attributes (here, *n*=12), and β_i,t_ is a Boolean variable describing for each attribute at each time step if the attribute value is within the estimated range (β_i,t_=1) or not (β_i,t_=0). The optimal values of α_M_=1 can be reached if all attributes of the simulated forest are within the estimated ranges continuously from *t*_min_ to *t*_max_. This approach assumes that a sufficiently stable equilibrium state is modelled, and the time to reach this state when starting from bare soil is thus an additional aspect we considered as validation criterion.

In addition, we evaluated whether our model adequately reproduced more complex patterns at the forest level. This mainly involved qualitative comparisons between simulations and observations based on visualized patterns. The following patterns were considered: (I) Crown architectures of trees in stands usually change markedly with tree height, and while the crown area usually shows a non-linear relationship with height, crown width and branching height are commonly rather linearly correlated with height (Alves and Santos 2002, Iida et al. 2011). (II) The vertical leaf area density in undisturbed forests within stands often peaks in the upper canopy, sometimes with an additional peak in the understory (Stark et al. 2012, Taubert et al. 2015). (III) The height-diameter relationship is a typical characteristic of a forest and for the Neotropics, this relationship is best described using a three-parameter exponential equation with an asymptotic maximum height of 38.8 m (Banin et al. 2012). (IV) The frequency distributions of tree diameter, height and age are typically right-skewed when considering all trees in a stand (Campbell et al. 1986, Worbes et al. 2003). When considering only trees above 10 cm in diameter at breast height (dbh), a normal or slightly right-skewed distribution is commonly observed (Oliveira-Filho et al. 1994, Worbes et al. 2003). To visualize the patterns, we used simulated data after reaching dynamic equilibrium state in intervals of 50 years to avoid temporal autocorrelation.

#### Best parameter combination

With the best parameter combination, we performed 10 replicate simulations for 1 ha forest plots to demonstrate our hypotheses that our FSFM successfully integrates tree level dynamics and species-rich forest stands while examining the effects of model-inherent stochasticity. However, as the patterns were consistent across all replicates, the results of a single model run are shown here to highlight temporal fluctuations.

## Results

We found an appropriate combination of parameter values that reproduced the ecological pattern at the tree and forest level simultaneously (Table 2; Appendix S1: Table S6). The emergent patterns using this best combination of parameter values are described below. Additonnally, the effects of variations of the free model parameters and the two main traits (SLA, wood density) on forest model results are shown in Table 4.

**Table 4.**
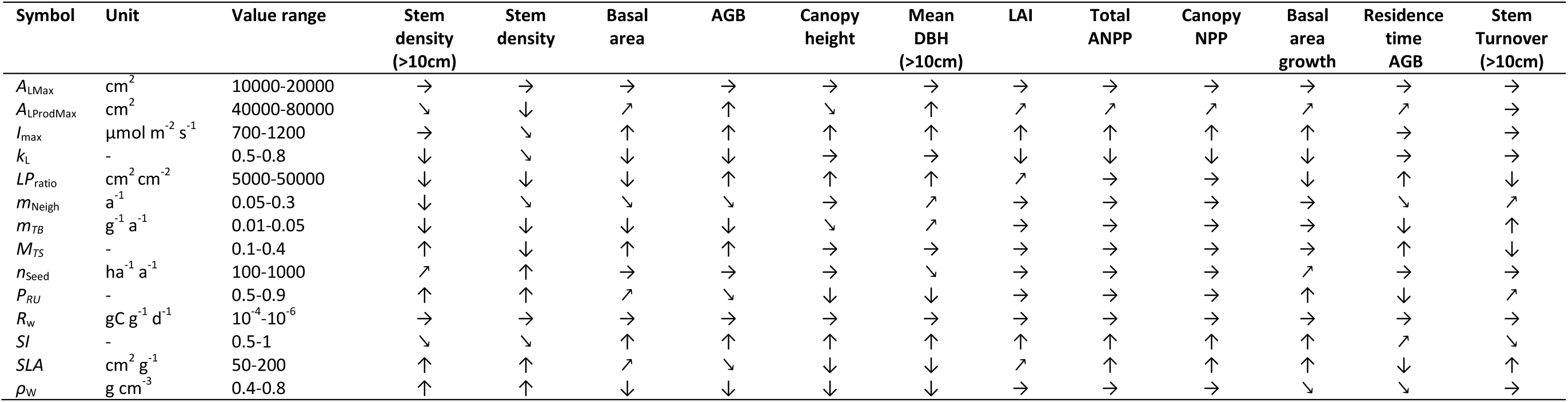
Results of sensitivity analysis, in which the effects of changes in free model parameters and two traits (SLA, wood density *ρ*_W_) on the main forest attributes were recorded. The parameters of the best model (Table 2) form the basis of this analysis, and single parameters were varied over the range shown in the column ‘Value range’. The arrows indicate whether increasing parameters values had a strong positive effect (↑), a slight positive effect (↗), no or an indifferent effect (→), strong negative effect (↓), or a slight negative effect (↘) on the forest attributes.

| Symbol | Unit | Value range | Stem density<br>(>10cm) | Stem density | Basal area | AGB | Canopy height | Mean DBH<br>(>10cm) | LAI | Total ANPP | Canopy NPP | Basal area growth | Residence time AGB | Stem Turnover<br>(>10cm) |
| --- | --- | --- | --- | --- | --- | --- | --- | --- | --- | --- | --- | --- | --- | --- |
| $A_{LMax}$ | $\text{cm}^2$ | 10000-20000 | → | → | → | → | → | → | → | → | → | → | → | → |
| $A_{LProdMax}$ | $\text{cm}^2$ | 40000-80000 | ↘ | ↓ | ↗ | ↑ | ↘ | ↑ | ↗ | ↗ | ↗ | ↗ | ↗ | → |
| $I_{max}$ | $\mu\text{mol m}^{-2} \text{s}^{-1}$ | 700-1200 | → | ↘ | ↑ | ↑ | ↑ | ↑ | ↑ | ↑ | ↑ | ↑ | → | → |
| $k_L$ | - | 0.5-0.8 | ↓ | ↘ | ↓ | ↓ | → | → | ↓ | ↓ | ↓ | ↓ | → | → |
| $LP_{ratio}$ | $\text{cm}^2 \text{cm}^{-2}$ | 5000-50000 | ↓ | ↓ | ↓ | ↑ | ↑ | ↑ | ↗ | → | → | ↓ | ↑ | ↓ |
| $m_{Neigh}$ | $\text{a}^{-1}$ | 0.05-0.3 | ↓ | ↘ | ↘ | ↘ | → | ↗ | → | → | → | → | ↘ | ↗ |
| $m_{TB}$ | $\text{g}^{-1} \text{a}^{-1}$ | 0.01-0.05 | ↓ | ↓ | ↓ | ↓ | ↘ | ↗ | → | → | → | → | ↓ | ↑ |
| $M_{TS}$ | - | 0.1-0.4 | ↑ | ↓ | ↑ | ↑ | → | → | → | → | → | → | ↑ | ↓ |
| $n_{Seed}$ | $\text{ha}^{-1} \text{a}^{-1}$ | 100-1000 | ↗ | ↑ | → | → | → | ↘ | → | → | → | ↗ | → | → |
| $p_{RU}$ | - | 0.5-0.9 | ↑ | ↑ | ↗ | ↘ | ↓ | ↓ | → | → | → | ↑ | ↓ | ↗ |
| $R_w$ | $\text{gC g}^{-1} \text{d}^{-1}$ | $10^{-4}$-$10^{-6}$ | → | → | → | → | → | → | → | → | → | → | → | → |
| $SI$ | - | 0.5-1 | ↘ | ↘ | ↑ | ↑ | ↑ | ↑ | ↑ | ↑ | ↑ | ↑ | ↗ | ↘ |
| $SLA$ | $\text{cm}^2 \text{g}^{-1}$ | 50-200 | ↑ | ↑ | ↗ | ↘ | ↓ | ↓ | ↗ | ↑ | ↑ | ↑ | ↓ | ↑ |
| $\rho_w$ | $\text{g cm}^{-3}$ | 0.4-0.8 | ↑ | ↑ | ↓ | ↓ | ↓ | ↓ | → | → | → | ↘ | ↘ | → |

### Tree level

Tree-level simulations revealed that under constant environmental conditions and without competition, tree growth could be divided into three successive life stages (Fig. 4b-d). In early developmental stages, there was a rapid increase in height and diameter as well as continuously increasing net primary production rates. In this stage, all major branches were foliated (Fig. 4a). Subsequently, the lower branches began to shed leaves (Fig. 4a), accompanied with lower increments in height and diameter and reduced net primary production (Fig. 4b, d). This stage ended when the tree reached its maximum height (see below for more details on maximum tree height). In the subsequent senescence stage, height and diameter growth ceased and net primary production decreased via loss of photosynthetically-active leaf biomass. The leaf loss continued until the tree ultimately died from carbon starvation (i.e., emergent senescence).

**Fig. 4.**
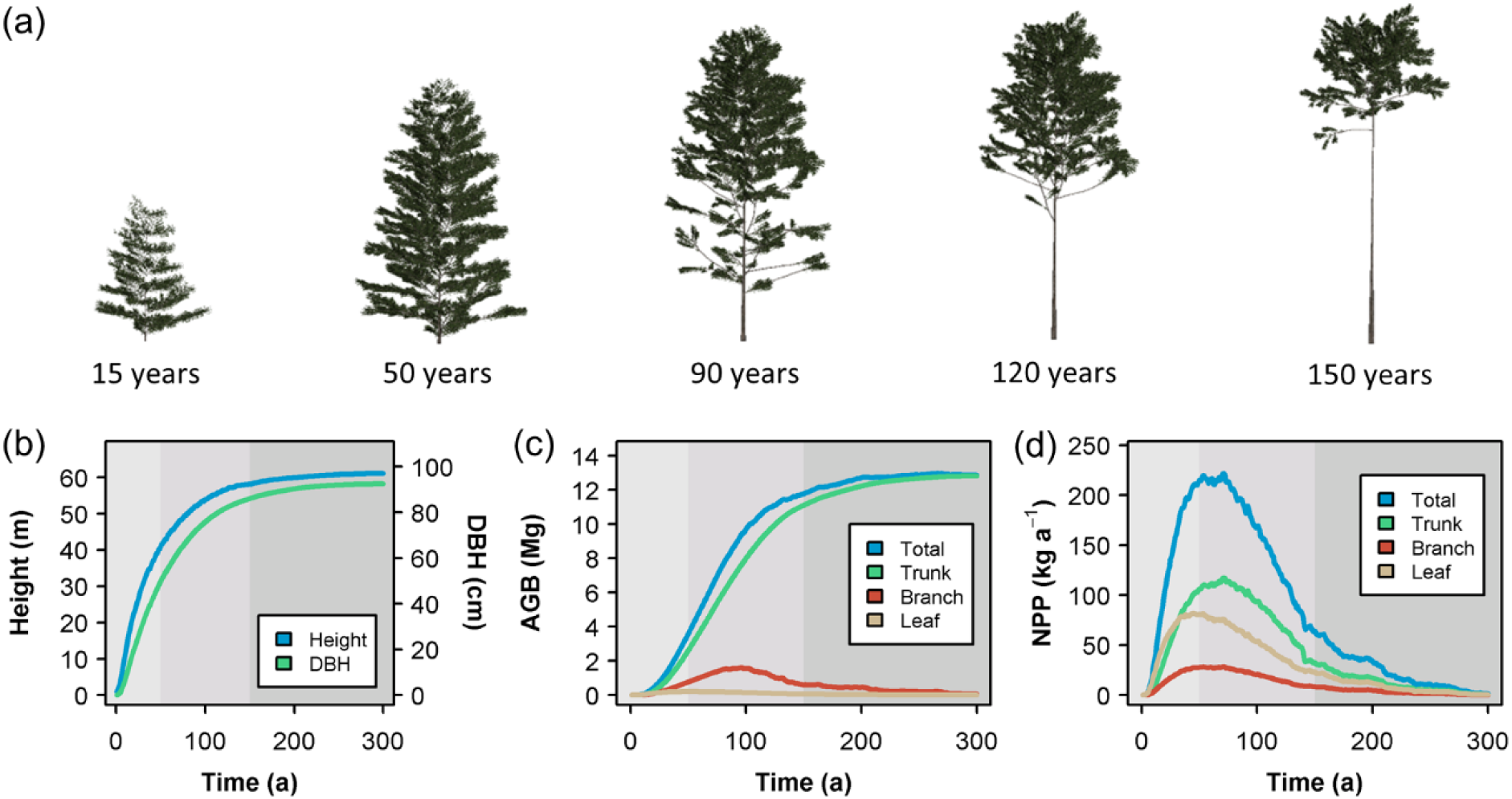
Ontogenetic development of an individual tree. (a) Structure of a freestanding tree at different ages. (b) Changes in tree height and diameter, (c) above-ground biomass (AGB) of the trunk, branches, leaves and the entire tree, and (d) net primary production (NPP) of the trunk, branches, leaves and the entire tree over time (panels b-d correspond to the tree shown in a). This example shows how a long-lived emergent tree species characterized by a low SLA grows without competition with neighbors over 300 years. Growth can be roughly divided into three life stages, which are indicated by different gray tones in panels b-d. The first stage is characterized by a quick increase in height and diameter, and continuously increasing net primary production. In this example, it ends at an age of ∼50 years. In the succeeding stage, NPP decreases and the tree sheds lower branches. The final senescent stage begins at ∼150 years when the tree growth close to its maximum height. In this stage, it gradually sheds all leaves and branches, which results in death.

When grown under constant external conditions, species generally followed the illustrated tree growth pattern over their life spans. However, tree traits and environmental conditions had a large influence on all aspects of growth (Fig. 5). Species with high SLA showed high initial growth rates (Fig. 5c) and rapidly reached their maximum height. Consequently, they entered the senescence stage after a shorter time and died at a comparably young age (Fig. 5a-d). In contrast, low-SLA species had a lower growth rate, but were able to maintain their growth rate for a longer time. They reached larger maximum heights later and had longer life spans. Wood density also affected life history growth, mainly by influencing maximum height (Fig. 5e-h). Both external factors (light and site index) affected tree growth in a similar way (Fig. 5k-o). Due to the trade-off between carbon gain and carbon costs, lower light and site index values also decreased maximum height. In contrast to SLA, variations in external factors influenced maximum height, but longevity was only slightly affected (Fig. 5i-p).

**Fig. 5.**
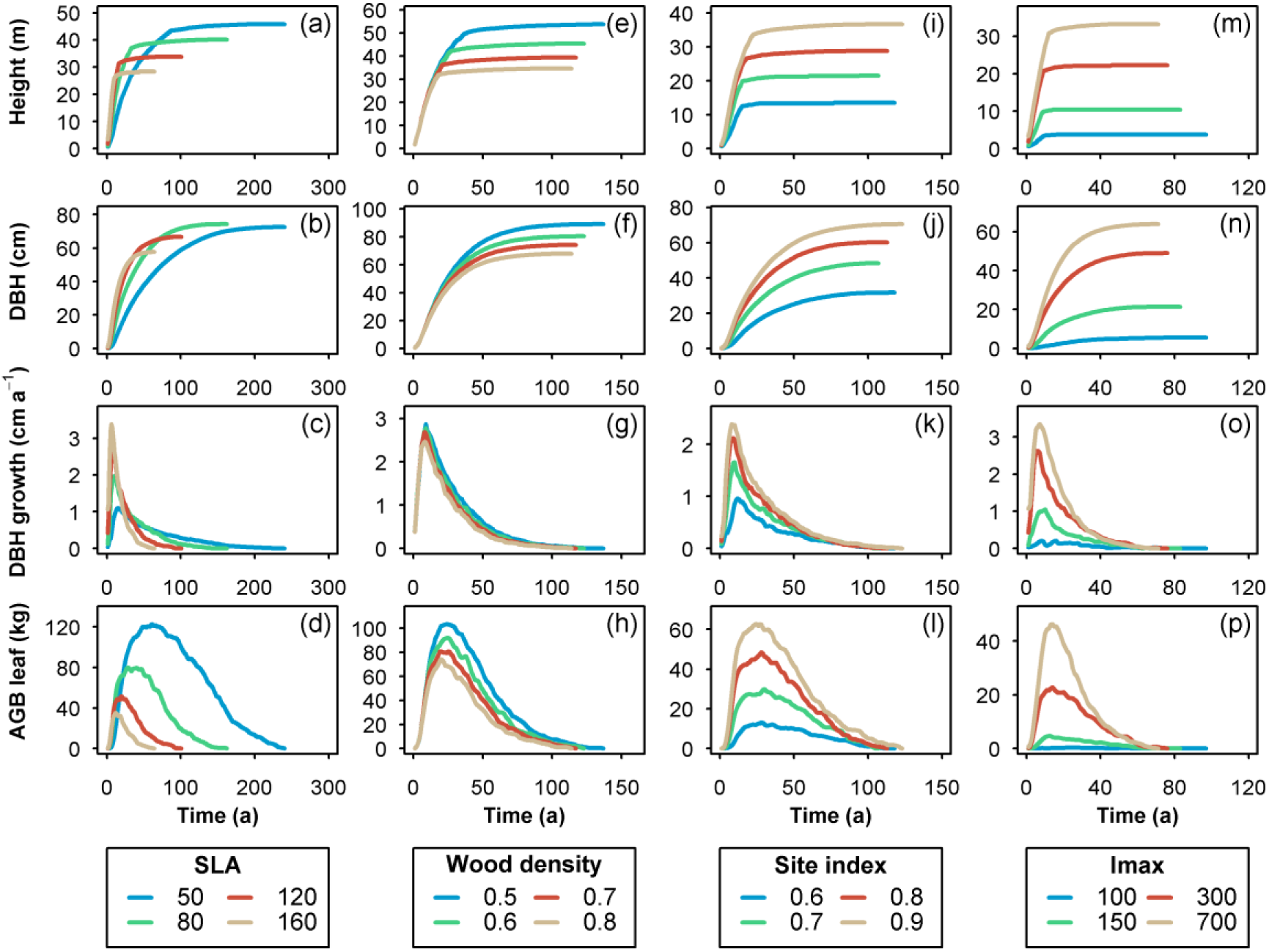
Tree dynamics as a function of their traits and the environmental conditions. Development of height, diameter, diameter growth rates and leaf biomass of trees with otherwise identical functional and structural traits, differing only in their specific leaf area (SLA; a-d) or their wood density (e-h). The right-hand panels illustrate the effects of the site index (i-l) and the light intensity *I*_max_ (m-p) on growth of trees with an identical set of traits.

The potential maximum height of a tree and the light compensation point for each leaf compartment were emergent properties (Fig. 6; see Appendix S1: Eq. 41 for details on maximum height calculation). Both the internal traits of a tree and the external environmental conditions determined the maximum attainable height, which generally decreased with increasing SLA and wood density (Fig. 6a), and increased with light and site conditions (Fig. 6b). The light compensation point of each leaf compartment was a function of its pipe length, as construction and maintenance costs increased (Fig. 6c). With short pipe lengths, species with high SLA values had a lower light compensation point than those with low SLA. However, the compensation point of high-SLA species steeply increased with increasing pipe length, while the increase was shallower for low-SLA species (Fig. 6c). Consequently, the latter ones had lower light compensation points already above ca. 4 m, i.e., in the understory (Fig. 6c).

**Fig. 6.**
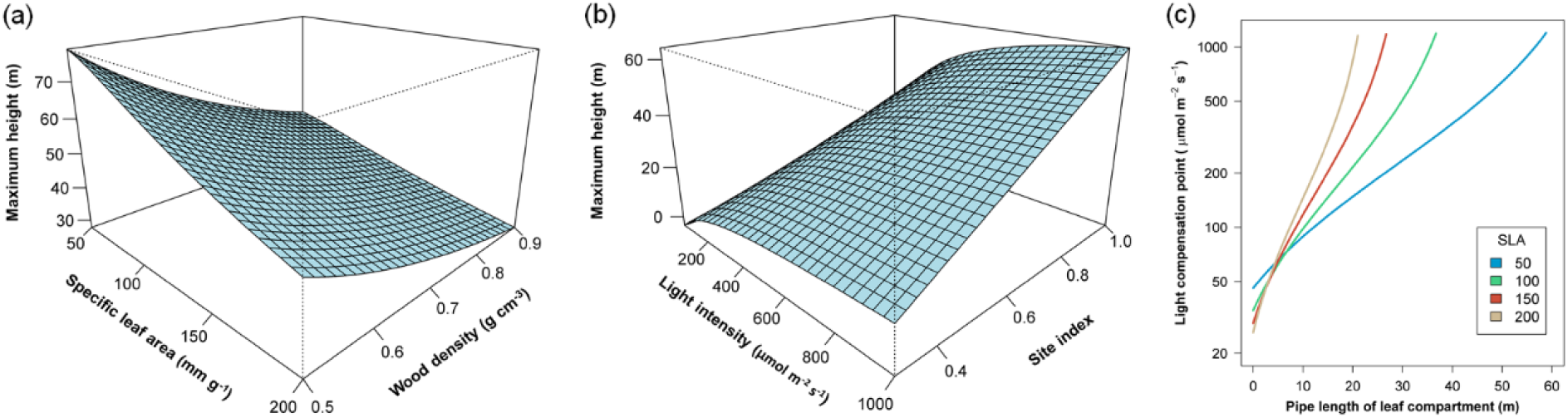
Maximum tree height as a function of (a) tree traits and (b) environmental conditions. The maximum height of a tree is directly related to the maximum pipe length *L*_PMax_, which is an emergent model property in our model (see Appendix S1: Eq. 41). The maximum height decreases with (a) SLA and wood density, and increases with (b) light intensity and site index. (c) Light compensation point of a leaf compartment in dependence on SLA and pipe length. The compensation point represents the light intensity at which assimilation and respiration rates match. Each leaf compartment of a tree forms a leaf-pipe element that acts as autonomous module, and consequently, the light compensation point can be assessed at the leaf compartment level.

### Stand level

Starting from bare soil, the simulated forest increased in stem number, above-ground biomass, and basal area, reaching a dynamic equilibrium after ca. 80-100 years (Fig. 7). In this equilibrium state, all 12 monitored forest attributes were within the ranges of typical Neotropical lowland forest (α_M_=1, see 1 ha stand result in Fig. 7). Fluctuations around the equilibrium increased with decreasing stand area as the relative effect of gap-creating mortality events was weaker at larger plot sizes (Fig. 7).

**Fig. 7.**
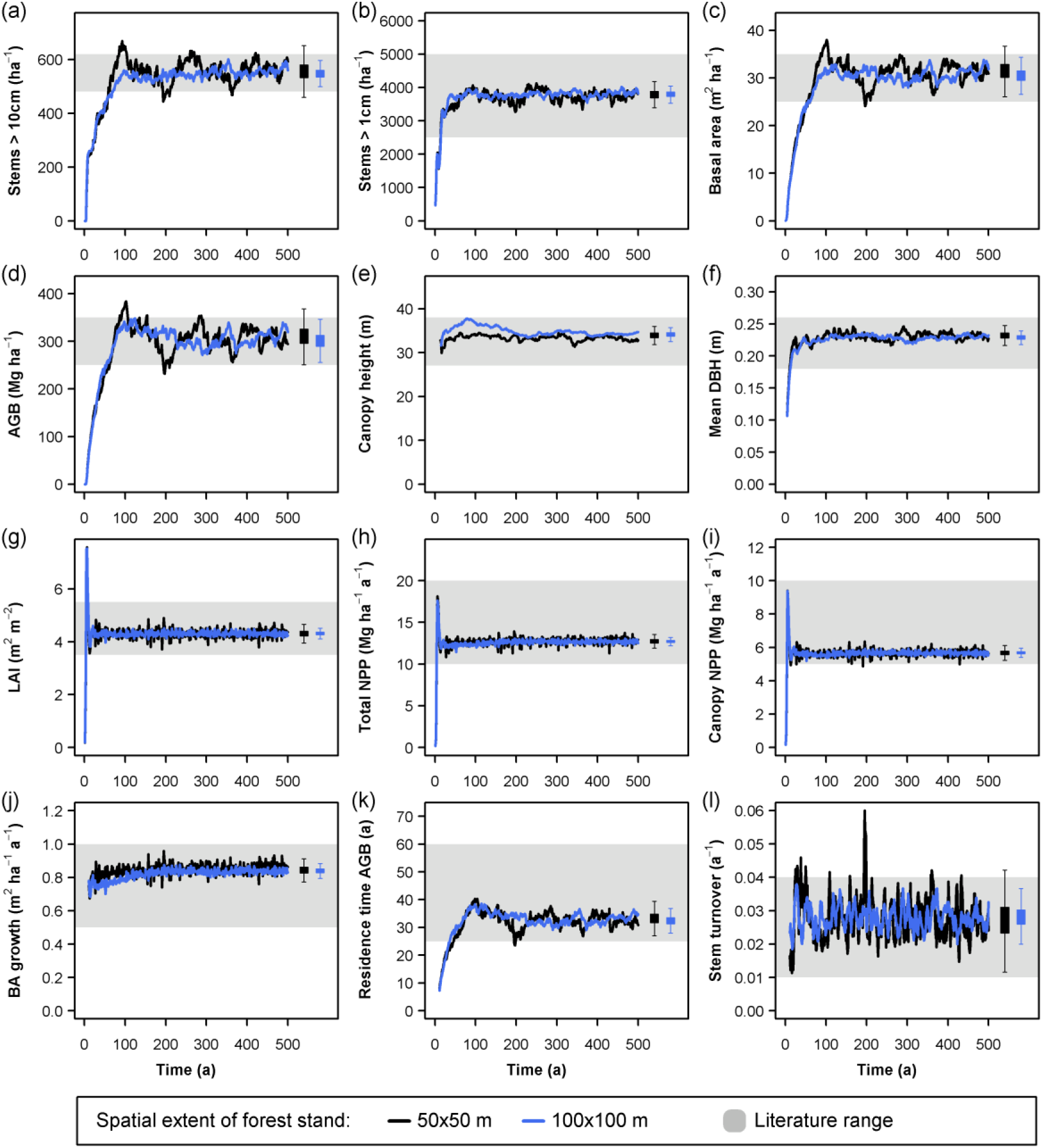
Simulated long-term forest dynamics. (a) Stem density of all stems with diameter at breast height (dbh) > 10 cm, (b) Stem density of all stems with dbh > 1 cm, (c) Basal area, (d) Above-ground biomass (ABG), (e) Canopy height (mean height of all trees > 40 cm in dbh), (f) Mean dbh of all stems > 10 cm in dbh, (g) Leaf area index (LAI), (h) Total above-ground net primary production (NPP), (i) Canopy net primary production (NPP of leaves and second order branches), (j) Basal area (BA) growth, (k) Residence time of above-ground biomass, (l) Turnover of all stems > 10 cm in dbh. Black lines represent simulations at 0.25 ha scale, and blue lines at 1 ha scale. The grey-shaded areas indicate typical ranges of the forest attributes in Neotropical forests (Table 3). Boxplots show interquartile ranges (boxes) and approximate 95% confidence intervals (whiskers) of the forest attributes in dynamic equilibrium state, i.e., from years 200-500, based on 10 replicates.

In dynamic equilibrium state, tree dbh, height, and age showed right-skewed distributions (Fig. 8). Trees reached maximum diameters of ∼100 cm, maximum heights of ∼ 50 m and maximum ages of ∼250 years. When only considering stems >10 cm in dbh, height and age were nearly normally distributed (Fig. 8a, c). On log-scale, tree numbers decreased almost linearly with dbh and age, with deviations from this pattern only at large size/age classes. In contrast, tree height peaked between 25 and 35 m (Fig. 8b). On log-log-scale, tree height distribution was linear for small individuals but became curvilinear at larger diameters (Fig. 8d).

**Fig. 8.**
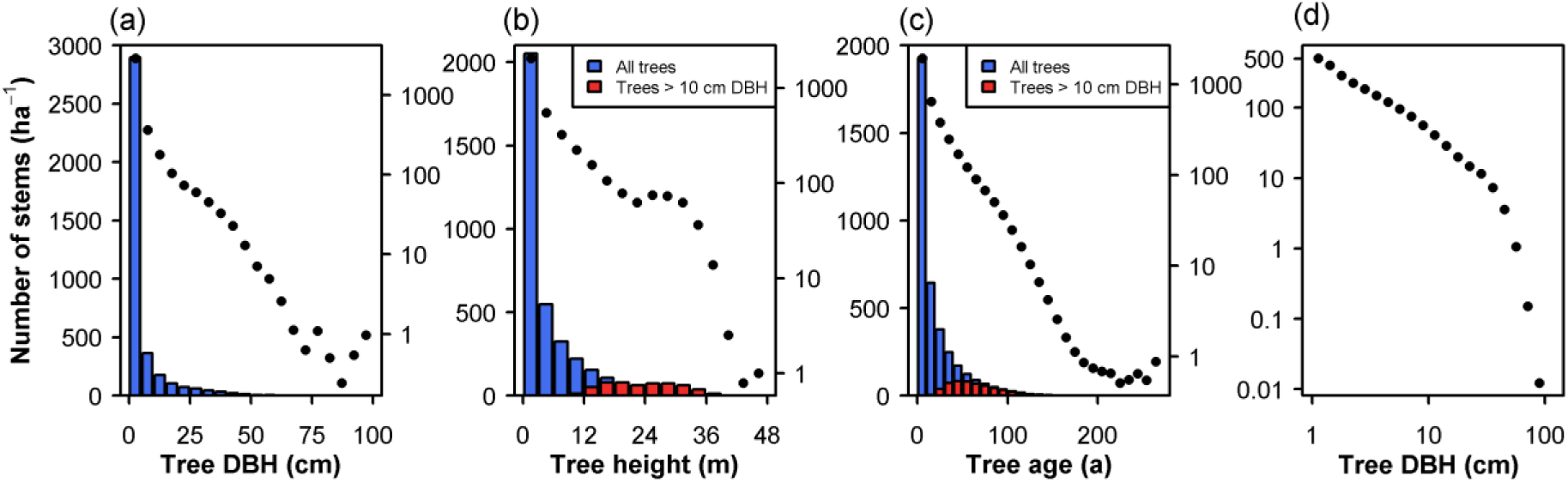
Frequency distributions: (a) tree diameter at breast height (dbh), (b) tree height, and (c) tree age. Distributions are shown on normal (colored bars, left axes) and log scale (black dots, right axes). The average frequency in each size class over the years 200-500 (equilibrium state) is shown here. (d) Tree dbh distribution on log-log scale. Values in each size class were binned to the class width.

Crown architectures changed significantly with tree height (Fig. 9a-c). There was a positive linear relationship between tree height and crown width (Fig. 9b), and an exponential relationship between tree height and branching height (Fig. 9a), as well as between tree height and crown area (Fig. 9c). However, due to differences in functional and structural traits and variations in ontogenetic stages, there was substantial scatter around the average trends. Visualizations of the simulated forest stand illustrate the level of detail and structural realisms of the model (Fig. 9e-f).

**Fig. 9.**
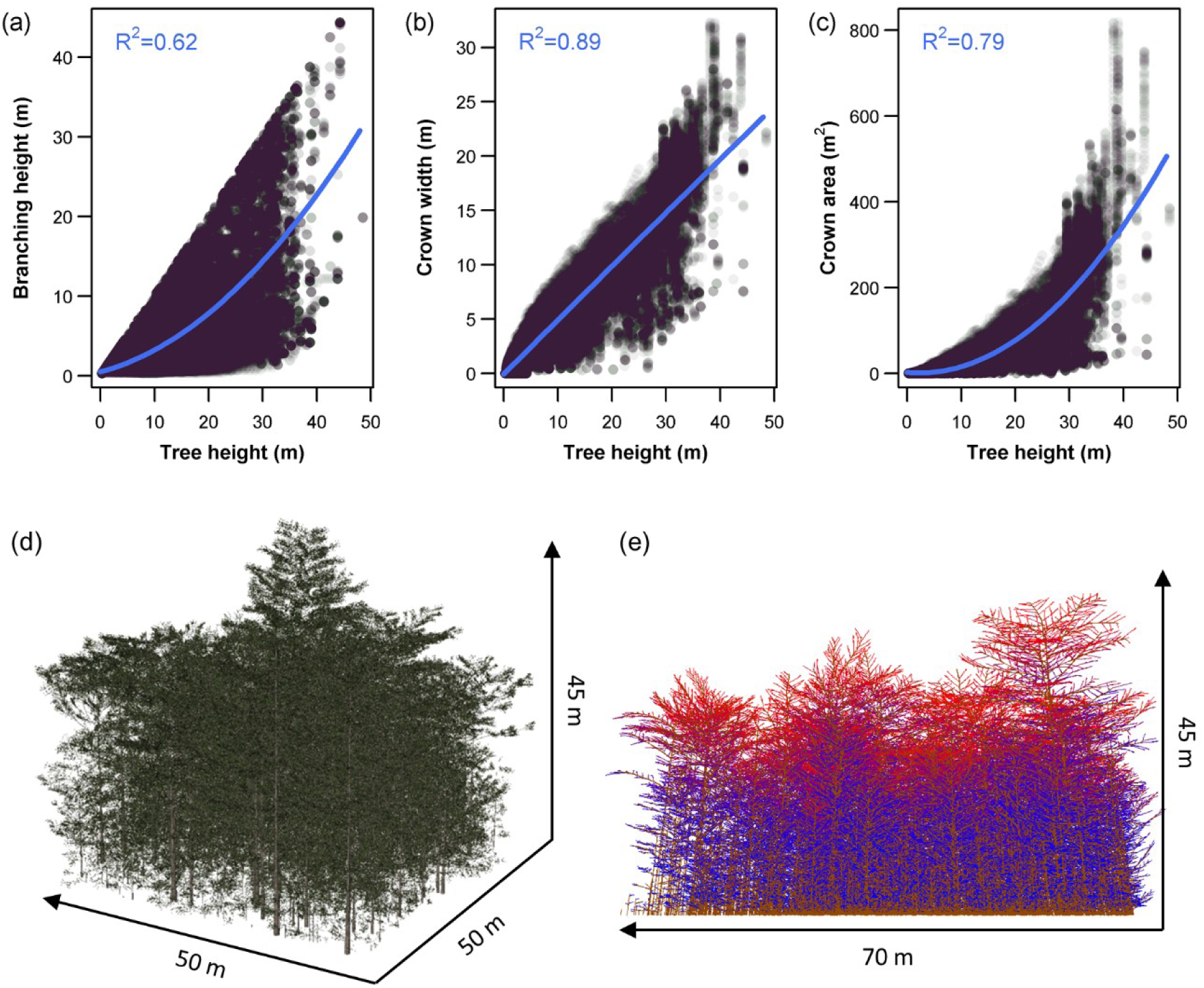
Relationship between tree height and crown parameters (a-c), and visual representation of the simulated forest (d-e): (a) branching height (height of lowest first order branch), (b) crown width and (c) crown area. Each dot represents a single tree in the simulated forest stand. To reduce the degree of temporal pseudoreplication, all trees in the forest stand were sampled in time intervals of 50 years in dynamic equilibrium state (200-500 years). Simple linear models and linear models including a quadratic term were fitted to the data and the minimal adequate model based on AIC values is shown here (ΔAIC > 4). (d) Oblique top view on the simulated forest stand (50×50 m) using the optimal parameter value combination at a representative time step in in dynamic equilibrium state. (e) Side view on the simulated forest stand (70×70 m) using the best parameter value combination at a representative time step in the dynamic equilibrium state. Here, the wireframe representation is shown, where second order branches are colored according to the light conditions (see Appendix S1 for details). Colors represent the shift from high light intensities (red) to low light intensities (blue).

The height-diameter relationship deviated slightly from the average allometry previously reported for the Neotropics (Banin et al. 2012) and overestimated the asymptotic maximum height by ∼3 m (Fig. 10a). The average leaf area density profile of the simulated forest showed a unimodal distribution, in which the leaf area density peaked in the mid-canopy between 15 and 25 m (Fig. 10b).

**Fig. 10.**
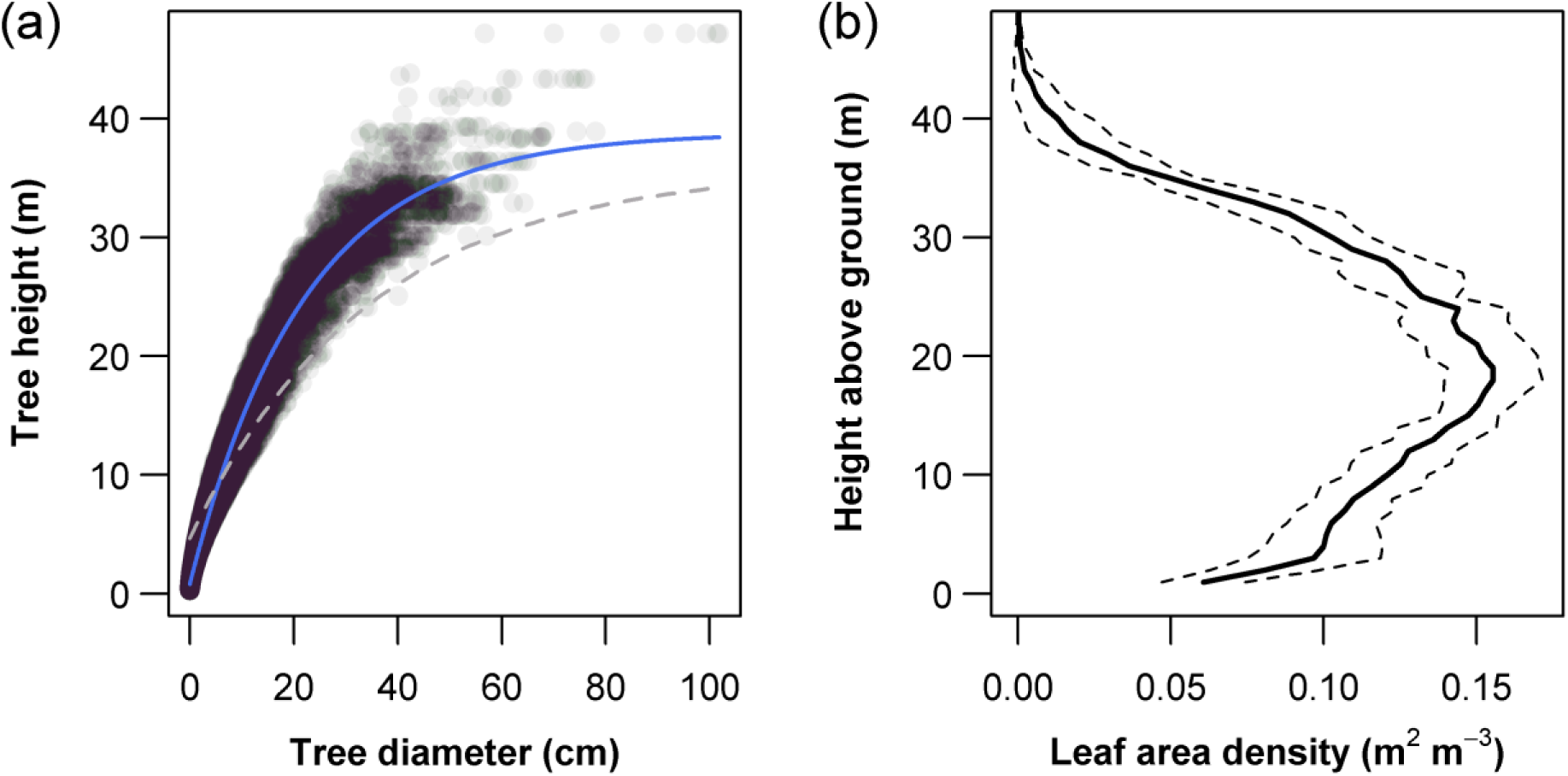
Height-diameter relationship (a) and vertical leaf area density distribution (b). (a) Each dot represents a single tree in the simulated forest stand. To reduce the degree of temporal pseudoreplication, all trees in the forest stand were sampled in time intervals of 50 years in dynamic equilibrium state (200-500 years). The relationship between tree height and diameter was described by the three-parameter exponential equation 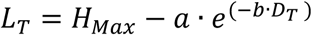, where *L*_T_ and *D*_T_ are the height and diameter of the tree, and *H*_Max_, *a* and *b* are curve parameters (*H*_Max_ represents the asymptotic maximum height, *a* the difference between maximum and minimum height, and *b* the shape of the curve). This equation was used by Banin et al. (2012), who estimated an asymptotic maximum height of *H*_Max_=35.8 (*a*=31.1, *b*=0.029) for Neotropical forests based on 49 forest plots (grey dotted line). Our model (blue line) predicted *H*_Max_=38.8, *a*=37.9 and *b*=0.045. (b) The vertical leaf area density profile was calculated based on the simulated total leaf area in each voxel *A*_LTot_. For each vertical 1 m layer, the mean *A*_LTot_ was estimated. The sold black line shows the means over all time step in dynamic equilibrium state (years 200-500), and the dotted lines indicate the standard deviation.

## Discussion

When comparing our model with other individual- or cohort-based forest models (e.g., Liu 1998; Köhler and Huth 1998; Huth and Ditzer 2000; Phillips et al. 2004a), two main differences emerge. First, our model simulates 3D tree structures in detail, including branch segments up to the second order and within-tree leaf distribution at high resolution (here, at 1 m^3^). This allows three-dimensional light and space competition within and between individuals. Second, tree species are not *a priori* classified into distinct functional groups (e.g. Köhler & Huth 1998; Tietjen & Huth 2006), but drawn from the full continuous trait space of leaf and wood traits. Here, as suggested by Wright et al. (2004), the leaf investment strategy of each species depends on the position on the continuous LES. Growth rate, shade tolerance, and maximum height of each tree species, typically defined *a priori* in other models, emerge in our approach from the leaf investment strategy. By combining functional and structural realism at the level of tree components, our model reproduced a high number of real-world patterns at the tree and the stand level. This demonstrates the suitability of our ecophysiological approach and that life-history variation among tree species emerge from their differences in leaf functional strategy.

### Tree level

Variations in leaf traits had a strong impact on life-history pattern in our model. Species with a high SLA were characterized by faster growth, shorter life spans, lower potential heights (Fig. 5), and lower shade tolerance (Fig. 6c). All of these characteristics are typical for pioneer species, which have significantly higher SLA than shade-tolerant species in nature (Kitajima 1994, Rijkers et al. 2000). With decreasing SLA, life history characteristics changed towards those of climax species. Several studies indicate that the transition from fast growing, short-lived pioneers to slow growing, long-lived shade-tolerant species is indeed rather continuous (Wright et al. 2003, Poorter and Bongers 2006), and our bottom-up ecophysiological approach supports the notion that leaf trait trade-offs influence the ecological strategy of trees.

At first glance, it may be counterintuitive that pioneer species with their generally high SLAs are shade-intolerant. A high SLA means a high photosynthetically active leaf area per dry mass investment (Evans and Poorter 2001), and observed growth rates of pioneer seedlings in shade were indeed higher than those of shade-tolerant species (Kitajima 1994). Furthermore, as shade leaves within individual trees generally have a higher SLA than sun leaves (Rozendaal et al. 2006, Markesteijn et al. 2007), a positive relationship between SLA and shade tolerance could be expected. However, the increased efficiency of light capture with increased SLA comes at a cost, as such leaves are more susceptible to herbivory and physical damage and hence are short-lived (Coley 1983, Wright and Cannon 2001, Díaz et al. 2004). In our approach, both such leaf trait correlations and the woody pipes connected with the leaves are considered, with construction and maintenance costs increasing with increasing pipe length. Hence, pipe length, which is determined by the within-tree position of a leaf compartment, is an additional factor for resource use efficiency. We found that a low SLA was more efficient under low light conditions (i.e., had a lower light compensation point) as long as pipe length exceeded 4 m (Fig. 6c). This means that, in line with observations, species with low SLA emerged as more shade-tolerant in our proposed ecophysiological approach.

Wood density influenced tree growth much less than leaf traits. The most obvious effect was an increase in maximum height with decreasing wood density (Fig. 5). Observed relationships between maximum tree height and wood density are inconsistent across studies, and while the trend observed for Iberian canopy tree species agrees with our simulation results (Poorter et al. 2012), other studies found no significant relationships (Wright et al. 2010) or even positive trends (Osunkoya et al. 2007). Our ecophysiological processes are carbon-based, and by decreasing the construction and maintenances costs per volume of wood, low wood densities are advantageous and allow trees to grow taller. However, low wood densities usually decrease mechanical stability and increase vulnerability to hydraulic failure (Hacke et al. 2001, Anten and Schieving 2010). Such trade-offs among wood traits influence tree architecture and may have adverse effects on the maximum attainable height (van Gelder et al. 2006). Furthermore, correlations among wood traits might also explain why increasing mortality rates with decreasing wood density are consistently reported (Chao et al. 2008, Wright et al. 2010, Visser et al. 2016) - a pattern we did not obtain from the simulations (Fig. 5). Thus, only considering wood density is not sufficient to reproduce all interspecific differences related to wood traits. Consequently, our FSFM may be further improved by integrating additional mechanisms, such as mechanical and hydraulic stability of wood pipes.

Site quality, characterized by the dimensionless site index parameter, positively influenced tree growth rates and maximum tree heights (Fig. 5i-l). We were not explicitly integrating water/nutrient cycles or temperature dependencies in our FSFM. Hence, the site index represents a proxy for the strength of several factors limiting tree growth, such as low water availabilities, low temperatures, or poor soil conditions. Such limiting factors are generally associated with decreasing productivity and lower tree/canopy heights (Girardin et al. 2010, 2013, Pan et al. 2013). Therefore, site index variations in our ecophysiological approach generate tree dynamics that are qualitatively in accordance with these observations. Changing light conditions had a similar effect on tree growth, and the increase in growth rates with light intensity emerging from our FSFM is consistent with observed light-dependent responses of most tropical species (Kobe 1999, Rüger et al. 2011, Philipson et al. 2014).

Irrespective of environmental conditions and functional traits, tree growth was size-dependent, with characteristic ontogenetic trajectories (Figs. 4-5). A significant effect of size on growth rates has also been observed for most tree species in field studies (e.g., Rüger et al. 2011; Iida et al. 2014). However, growth rates may both increase and decrease with diameter (Rüger et al. 2012), or responses may be humped-shaped (Clark and Clark 1999, Davies 2001). These observed differences may be related to species-specific variability, or incomplete or unbalanced data sets; tree growth data from natural forests often does not cover the entire size range of species or is skewed towards the more frequent smaller size classes. This causes ambiguity when estimating ontogenetic growth patterns from field data, and whole-life growth trajectories are thus still debated (Rüger et al. 2011, Bowman et al. 2013). However, Hérault et al. (2011), found that the growth trajectories of 50 rainforest tree species could well be predicted using hump-shaped size-dependent models, and several additional empirical and theoretical studies suggest similar trajectories (e.g., Clark and Clark 1999). Our results generally agree with these studies, although the transition from increasing to decreasing growth rates at larger sizes in real trees seems smoother because trees exhibit a variety of non-implemented mechanisms to optimize their carbon budget. Integrating such mechanisms into our ecophysiological framework could improve the tree growth trajectories, and thus could be considered as scope for future studies.

### Stand level

As first indication for structural realism emerging from the integrated ecophysiological processes, a dynamic equilibrium was reached. It is generally assumed that under constant environmental conditions, carbon gains and losses are relatively balanced in old-growth forests, resulting in gap-driven oscillations around an equilibrium biomass level (Whitmore 1990, Galbraith et al. 2014). In accordance with simulations by Chambers et al. (2013), gap formation produced stronger oscillations in smaller stands (Fig. 7). The time required to reach equilibrium biomass after deforestation differs with environmental conditions, ranging from 40 to 60 years in high latitudes (Puerto Rico; Mitche et al. 2000; Marin-Spiotta, Ostertag and Silver 2007) to ∼190 years in Amazonian forests (Columbia and Venezuela; Saldarriaga et al. 1988). Hence, the time span between 80 and 100 years emerging in our simulations lies within the reported range, but may be too fast for equatorial forests. Nevertheless, our experimental design does not follow typical succession as we seed the plot with all species from the species pool (i.e. plots are seeded at a high functional diversity). In fact, such strategy of initial seeding with high diversity (> 50 species) and mixed functional types has been succesfully used in restoration in Neotropical forests, with near old-growth conditions being reached within 80-85 years (see Rodrigues et al. 2009 for a review). Furthermore, the time needed to reach dynamic equilibrium in a simulation model normally depends on the initial conditions and speed of internal dynamics. This is strongly controlled by species trait composition and productivity, which in our model is mostly given by the site index, and a decreasing site index and decreasing mean SLA would slow the dynamics (Table 4). Moreover, note that typical components of an old-growth tropical forests that might require longer periods, such as lianas, epiphytes and strangler ficus, are not explicitly modelled. All these considerations suggest that 100 years to reach equilibrium biomass in a highly productive forest is indeed realistic.

In dynamic equilibrium, all 12 important forest attributes fell within the observed ranges of Neotropical lowland forests (Fig. 7). We chose using ranges instead of fitting our model to observed forest attributes of individual forest plots because the few plots with repeated inventories over years to decades focus on only a few key forest attributes (e.g., Condit 1995; Bradford et al. 2014). This means, we opted to consider as *many* attributes in reasonable ranges as possible by considering many forest plots, instead of reproducing *some* attributes with a higher accuracy for specific forest plots. The lower number of reported patterns in the alternative parameterization strategy would have increased the risk of equifinality, further limiting ecophysiological insights.

In fact, several complex patterns at the forest level could also be realistically represented (Figs. 9-10). The right-skewed tree diameter distribution (Fig. 8a) is consistent with observations (Oliveira-Filho et al. 1994, Hector et al. 2011). Simulated normal or slightly skewed distributions for height and age of trees > 10 cm in dbh (Fig. 8b, c) also agree with empirical studies (Campbell et al. 1986, Oliveira-Filho et al. 1994, Worbes et al. 2003, Adekunle et al. 2013). As diameter, height, and age are usually correlated, similar frequency distribution of these attributes can be expected. Interestingly, the height distribution deviated from the age distribution and showed a slight hump between 25 and 35 m (Fig. 8b). We speculate that the crowns of these trees in the upper canopy are well illuminated and less exposed to between-crown competition for space. Consequently, mortality by carbon starvation decreases, explaining why the otherwise generally negative trend is temporarily stopped in this height class.

The metabolic theory of ecology predicts a linear decrease in stem diameter frequency on a log-log scale. Muller-Landau et al. (2006) and Enquist, West and Brown (2009), however, found deviations from this theoretical prediction particularly among the larger diameter classes, whose frequency was lower than predicted. Interestingly, our model shows similar deviations (Fig. 8d). Enquist, West and Brown (2009) speculated that other sources of mortality than competitive thinning, which is the reason for the predicted linear trend, for instance wind damages, lightning strikes, or diseases, are particularly severe in larger size classes. In our FSFM, trees growing at their maximum height enter senescence, which inevitably ends with death. This emergent behavior increased mortality probability of very large trees and thus is close to observations while still considering metabolic-driven mortality.

The linear increase in crown width with height is in accordance with approximately linear trends observed in many tropical tree species (King 1996, Alves and Santos 2002, Iida et al. 2011). Interestingly, a bulge in the crown width-tree height relationship was observed for tall trees > 35 m (Fig. 9b), indicating a disproportionate increase in crown width when trees rise above the average canopy height. King (1996) made similar observations for larger trees potentially growing above the canopy. Crown development of such emergent trees is less constrained by competition for space, and our ecophysiological approach can reproduce such plastic crown responses. With an average crown width of ∼10 m at a height of 20 m, our model only slightly overestimates crown dimension compared to observations, which are in a reported range of 7-9 m (King 1996, Alves and Santos 2002). Due to the relatively simple integration of space competition in this model, particularly ignoring wind-driven crown movement, the area of overlap between two crowns may be larger than in nature, leading to wider crowns.

Branching height showed a slightly curvilinear trend with tree height (Fig. 9a). Although most studies reported linear species-specific relationships (e.g., Alves and Santos 2002; Iida et al. 2011), the average slopes of simulated and observed relationships agree quite well. Compared with the other crown measures, branching height was less strongly correlated with tree height (Fig. 9). In our model, branch growth followed the architecture defined by structural traits, and branch shedding resulted from physiological process. These processes lead to a distinct species-specific branching height during tree ontogeny (Fig. 4), which could be modified, however, by competition and environmental variables (Appendix S1: Fig. S10). In real trees, branching architecture is a complex trait, and the processes of branch growth and shedding are likely related to within-tree optimization of carbon gain (Farnsworth and Niklas 1995). For example, the complex crowns of emergent trees often develop through light-mediated activation of dormant buds (Hallé et al. 1978, Barthélémy and Caraglio 2007). Due to such mechanisms, which are not yet integrated in our FSFM, the correlation between branching height and tree height is probably stronger in real forests. Unfortunately, we are not aware of any study reporting the strength of correlation between branching height and tree height at stand scale, which would allow direct comparisons.

The simulated leaf area density peaked at ∼20 m (Fig. 10b), which is in accordance with observations of the canopy in Neotropical lowland forests on Barro Colorado Island (Taubert et al. 2015) and near Manaus (Stark et al. 2012). In the latter study, an additional increase in leaf area density near the forest floor was observed, probably due to herbaceous vegetation and shrubs, which are not included in our model. Although vertical leaf area profiles have not been as extensively studied as other forest attributes, a leaf area maximum in the canopy layer is generally expected for undisturbed old-growth forests and our simulations are in line with this expectation. Interestingly, in a lowland forest analyzed by Stark et al. (2012), the leaf area density peaked in the lower canopy around 10 m. These authors considered past disturbances and the resulting non-equilibrium forest state as a possible cause of the deviating pattern. Analyzing the effect of disturbance regimes on the stand-level vertical leaf area distribution may thus be an interesting future application of our model.

The asymptotic height of the diameter-height relationship observed here (38.8 m; Fig. 10a) is close to the observed mean for Neotropical forests (35.8 ± 6.0 m; Banin et al. 2012). However, the shape of the simulated relationship differed from the observed average trend, and the simulated height at smaller diameter classes was slightly overestimated (Fig. 10a). In our model, height scales with diameter to the power 2/3, controlled by species-specific shape parameters. Trees only deviate from their species-specific relationship under low light conditions (increased height growth) or at maximum height (cessation of height growth). In nature, trees show a plastic response to environmental factors such as light, precipitation or stand density, which can alter the specific allometric relationships (Feldpausch et al. 2011, Banin et al. 2012). Trees may, for instance, cease height growth when growing as emergents under full light. Including more such mechanism in FSFMs may thus improve the emergent allometric scaling.

## Limitations

We were able to find parameter values yielding realistic ecological patterns for our FSFM by manually applying pattern-oriented modeling (Grimm et al. 2005, Grimm and Railsback 2012). However, the problem of equifinality persists, i.e., we cannot rule out the possibility that other parameter value combinations would lead to similarly good or better results. Inverse modeling techniques such as Bayesian approaches (e.g., Martínez et al. 2011; Matsushita et al. 2015) could help to explore the entire parameter space more systematically and to quantifying local/global minima, parameter uncertainties, and collinearities among parameters. However, due to the considerable computation time of our model and the lack of plot data spanning the entire set of variables evaluated here, such systematic model calibration is currently unfeasible. Nevertheless, that fact that we found a suitable parameter value combination indicates that our ecophysiological framework can generate realistic tree and forest structure. Further ecophysiological insights might be gained when empirical data becomes available at the same structural level than our FSFM, such as time series data for several tree components, or for detailed tree and forest attributes of the same forest stands (e.g., via repeated LiDAR data).

Some ecological patterns were not accurately reproduced by our model. For instance, trees in natural forests commonly show fast initial height growth rates, but when crowns are well-illuminated, they tend to cease height growth and continue to mostly grow in diameter (e.g., Matsushita et al. 2015). In this model, height and diameter growth are coupled more tightly, and the time frame over which high diameter growth rates at large heights can be maintained appears larger in nature (Clark and Clark, 1999) than in the ecophysiological dependencies depicted in our framework. This deviation may be explained by intra-individual trait plasticity. In our FSFM, each individual is characterized by non-variable traits, whereas in nature SLA is typically adjusted to local light conditions (Rozendaal et al. 2006, Markesteijn et al. 2007). Furthermore, trees can modify their architecture to avoid self-shading of branches and to maximize carbon gain. In addition, activation of dormant buds is regarded as important additional branching mechanism (Hallé et al. 1978, Sterck and Bongers 2001, Osada 2011). If such plastic responses were integrated, larger trees would likely use resources more efficiently, ultimately maintaining higher growth rates over a longer period and reaching larger diameters and ages.

## Conclusion and perspectives

The position of leaf traits along the leaf economics spectrum determined the maximum height and age, size-dependent growth rates and shade tolerance, as well as a smooth continuous transition from fast-growing, short-lived pioneers to slow-growing, long-lived climax species as seen in the tree-level patterns when varying SLA. These emergent results were remarkably consistent with well-known functional tree types. Therefore, our model reveals a fundamental role of leaf traits in determining a) life history growth patterns of trees and b) structural, functional, and temporal dynamics of species-rich tropical forests.

Bottom-up functional-structural tree and forest models thus have the potential to increase our understanding of the mechanisms controlling tree and forest dynamics. Ecologists are able to empirically measure processes at lower organizational levels (e.g., photosynthesis at the leaf scale) or to track some tree variables over a limited time (e.g., diameter growth), but it is yet virtually impossible to record 3D tree growth as a whole. In this situation, functional-structural forest models can be helpful tools test and evaluate the consequences of low-level mechanisms on whole-tree growth patterns. So far, FSTMs have not received much attention, but we indicate that they offer ample opportunities for future studies both at the tree and forest levels. For instance, the importance of within-individual trait plasticity on whole-tree carbon budget could be explored. Also, the importance of frequent disturbances for forest stability and stand level trait distribution could be assessed. Moreover, further integration of currently missing mechanisms into FSTMs will facilitate the development of next-generation predictive forest models in which tree performance emerges directly and exclusively from continuous functional traits.

Finally, we highlight the potential of FSFMs for future model-based studies of canopy-dwelling organisms. Tropical forest canopies harbor numerous arboreal animals and epiphytic plants, but accessing their dynamic habitat is logistically challenging. FSFMs provide information on 3D forest dynamics and microclimatic changes, which can be used as input for studies on other organisms. For instance, our knowledge on long-term dynamics of vascular epiphytes is still very scarce, and by coupling a FSFMs with an epiphyte model, the importance of 3D forest dynamics on epiphyte communities could be investigated. Such integrative modeling studies are particularly timely as tropical forest dynamics are already changing in response to atmospheric changes, and a more detailed understanding of the response of canopy-dwelling organisms is paramount and urgently needed.

## Acknowledgements

GP and HK were funded by the DFG Initiative of Excellence Free Floater Program at the University of Göttingen. JSC acknowledges financial support by the German Research Foundation (DFG SA-21331).

