## Appendix S1 for "Modeling the long-term dynamics of tropical forests: from leaf traits to whole-tree growth patterns"

*Manuscript title: Modeling the long-term dynamics of tropical forests: from leaf traits to whole-tree growth patterns*

*List of authors: Gunnar Petter, Holger Kreft, Yongzhi Ong, Gerhard Zotz, Juliano Sarmento Cabral*

This appendix is divided into 3 sections: model description; external model control, export and visualization; figures and tables. In the first section, the forest model is described in detail. The second section contains information about the control files required to run the model, and about how to export and visualize results. The third section contains additional figures and tables, including all symbols and parameters.

#### 1 Model description

The functional-structural forest model (FSFM) developed in this study can be applied to simulate 3D structural growth of individual trees, or the (long-term) dynamics of forest stands at the plot level. The model description follows the ODD (**O**verview, **D**esign concepts, **D**etails) protocol, which was proposed as a standard protocol to communicate agent-based models or large, complex models (Grimm et al. 2006, 2010). The individual blocks of the protocol are subdivided as follows: **O**verview - Purpose, State variables and scales, Process overview and scheduling; **D**esign concepts - Basic principles, Emergence, Adaptation/Sensing, Interaction, Stochasticity, Observation; **D**etails - Initialization, Input, Submodels. To ensure a cohesive and completed model description in one document, the first block of the protocol (**O**verview), which is contained in the main manuscript, is also included here. The model was implemented using the open-source 3D modeling platform GroIMP (Growth Grammar Interactive Modeling Platform; available at www.grogra.de). In GroIMP, relational growth grammars are implemented in the programming language XL, which is a graph-based extension of the Lindenmayer-Systems (L-Systems), a formal language for the description of plant structure (Lindenmayer 1968a, 1968b). The model code required to run the FSFM is available at https://github.com/julianoscabral/MoF3D.

#### 1.1 Purpose

The FSFM serves two main purposes. First, to study the relationship between leaf trait trade-offs and life-history variation in trees. Ontogenetic growth patterns, maximum height, and life span, as well as the light-dependent growth behavior in our model emerge from continuous plant traits. Second, to generate long-term dynamics of forest stands at a high level of structural detail. By combining the leaf trait-based tree growth with local population and community dynamics, emergent patterns can be evaluated with field observations at the forest level. Our ecophysiological approach to forest dynamics was designed to increase the understanding of bottom-up mechanisms controlling forest dynamics, and, in addition, to be useful for follow-up studies requiring a detailed 3D representation of forest structure and dynamics.

#### 1.2 Entities, state variables and scales

The FSFM simulates establishment, growth, and mortality of virtual 3D trees at the plot level. The spatial and temporal scale of the forest plot can be defined during experimental design. Here, we simulate forest stands between 0.25 and 1 hectare over 500 to 1000 years in annual time steps. The vertical spatial dimension is associated with typical maximum tree heights (ca. 50-60 m). The 3D space is divided into a regular grid of cubic voxels with a side length of 1 m (Fig. S1a), which defines the spatial resolution of both light and leaf area/biomass distribution. Light is the main driver of vegetative growth and is calculated for all voxels based on the 3D distribution of leaf area.

This model comprises three hierarchical levels: tree components, individual trees, and the forest stand. Tree components are trunks, branches, apical meristems, and leaf compartments (Fig. S1b). Each tree consists of one erect trunk described by length and diameter. Attached to the trunk are branches up to the second order. Branches are defined at two different scales. At the coarse scale, branches are described by their total length and diameter, while at the fine scale branches are described as topologically connected smaller segments. This multi-scale approach optimizes both speed of the ecophysiological simulations and visual aspects. Located at the end of each trunk or branch, apical meristems sense the local environment and control primary growth. Leaf compartments are connected with second order branches and are conceptualized as aggregations of leaves within the cubic voxels. Leaf compartments comprise leaves and active pipes, which represent the sapwood connecting leaves and roots. This means that leaf compartments form leaf-pipe elements following the pipe model theory (Shinozaki et al. 1964; Fig S1b). Each tree component is characterized by state variables (Table S1), including its absolute 3D position and its topological position within its tree. Based on this information, the

3D structure of each tree can be deduced (Fig. S1c). Structural tree growth results from creation, loss, and dynamic changes of tree components. Structural tree growth is driven by the 3D distribution of light within the forest stand and the functional and the structural traits of trees (Table S1). While the functional traits regulate light-mediated carbon balance, the structural traits reflect inherent architectural parameters defining the tree´s structural organization. This includes, for instance, branching angles or average internode lengths (see submodel *Structural growth* for more details). Some functional trait combinations promote effective carbon assimilation under low light conditions, allowing growth and survival in the dark understory. Other trait combinations may be more favorable under high light conditions. Consequently, forest dynamics results from structural growth of individual trees with different traits competing for space and light, whose distribution, in turn, is influenced by the forest structure (Fig. S1d).

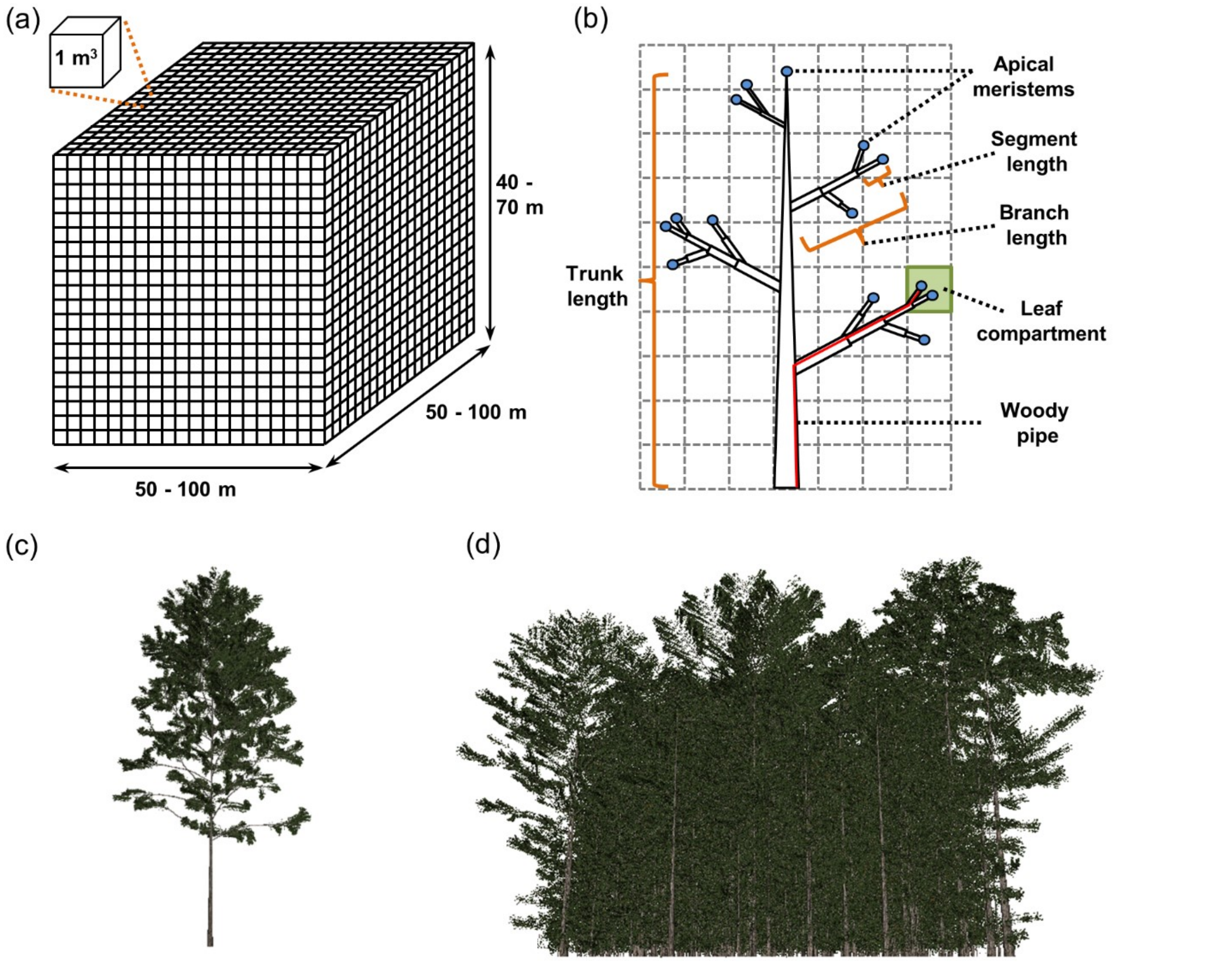


**Figure S1.** State variables, scales and visualization. (a) 3D model space. The extent of the model space can be defined by the user. The model space is a 3D grid that is subdivided into cubic voxels with a volume of 1 $m^3$ containing the information about local leaf biomass and area, as well as, light intensity. (b) Overview of tree components: trunks, branches, apical meristems, and leaf compartments. Trees consist of a trunk and branches up to the second order, which are terminated by an apical meristem. Leaf compartments describe the leaf biomass and area within a voxel attached to a specific section of a second order branch, as well as the woody pipes connected to these leaves. The length of the pipe system depends on the within-tree position. One leaf compartment (green square) and its woody pipe (red line) are exemplified. (c) 3D tree visualization. Tree structures are visualized based on the state variables and topology of each tree. Several visualization options are integrated in this model (see section *External model control, export and visualization* for more detail). Here, the leaf biomass in the leaf compartments is displayed by spatial objects imitating 'real' leaves. (d) 3D forest visualization. The forest structure can be displayed in this model, which allows visual inspections and comparisons with real forests.

**Table S1.** State variables, functional and structural traits of the forest model. Each tree component (trunk, branch, leaf compartment, apical meristem) is characterized by a set of state variables. The functional and structural traits describe the intrinsic properties of each tree species. Empirical correlations between leaf functional traits (Wright et al. 2004) are considered (see Table S2 for more details).

| Symbol | Description | Unit | Type |
|---|---|---|---|
| $A_B$ | Cross-sectional area of branch | $cm^2$ | State variable |
| $A_L$ | Leaf area in leaf compartment | $cm^2$ | State variable |
| $A_{LProd}$ | Total leaf area produced in leaf compartment | $cm^2$ | State variable |
| $A_S$ | Cross-sectional area of branch segment | $cm^2$ | State variable |
| $A_T$ | Cross-sectional area of trunk | $cm^2$ | State variable |
| $B_L$ | Leaf biomass in leaf compartment | g | State variable |
| $D_B$ | Diameter of branch | cm | State variable |
| $D_S$ | Diameter of branch segment | cm | State variable |
| $D_T$ | Diameter of trunk | cm | State variable |
| $I_M$ | Light conditions at apical meristem | $\mu mol\ m^{-2}\ s^{-1}$ | State variable |
| $L_B$ | Length of branch | cm | State variable |
| $L_P$ | Pipe length of leaf compartment (corrected after apical control) | cm | State variable |
| $L_{PS}$ | Pipe length of leaf compartment | cm | State variable |
| $L_S$ | Length of branch segment | cm | State variable |
| $L_T$ | Length of trunk | cm | State variable |
| $O_B$ | Branch order | - | State variable |
| $P_{BEnd}^{XYZ}$ | End position of branch (in X, Y and Z direction) | cm | State variable |
| $P_{BStart}^{XYZ}$ | Start position of branch (in X, Y and Z direction) | cm | State variable |
| $P_{LC}^{XYZ}$ | Position of leaf compartment (in X, Y and Z direction) | cm | State variable |
| $P_M^{XYZ}$ | Position of apical meristem (in X, Y and Z direction) | cm | State variable |
| $P_{SEnd}^{XYZ}$ | End position of branch segment (in X, Y and Z direction) | cm | State variable |
| $P_{SStart}^{XYZ}$ | Start position of branch segment (in X, Y and Z direction) | cm | State variable |
| $P_T^{XY}$ | Position of trunk (in X and Y direction) | cm | State variable |
| $G_{max}$ | Maximum gross photosynthetic rate | $g\ g^{-1}\ d^{-1}$ | Functional trait |
| $k$ | Light intensity at which the gross photosynthetic rate is half maximal | $\mu mol\ m^{-2}\ s^{-1}$ | Functional trait |
| $LL$ | Leaf lifespan | d | Functional trait |
| $N_{mass}$ | Nitrogen concentration | % | Functional trait |
| $R_L$ | Leaf respiration rate | $g\ g^{-1}d^{-1}$ | Functional trait |
| $SLA$ | Specific leaf area | $cm^2\ g^{-1}$ | Functional trait |
| $\rho_W$ | Wood density | $g\ cm^{-3}$ | Functional trait |
| $A_{LProd}$ | Maximal leaf area production per leaf compartment | $cm^2$ | Structural trait |
| $I_T$ | Light intensity threshold regulating apical dominance of SAM | $\mu mol\ m^{-2}\ s^{-1}$ | Structural trait |
| $k_{int}$ | Factor controlling the increase in internode length | - | Structural trait |
| $LD_B$ | Length-diameter ratio of branches | $m\ cm^{-1}$ | Structural trait |
| $LD_T$ | Length-diameter ratio of trunks | $m\ cm^{-1}$ | Structural trait |
| $L_{IBMax}$ | Maximal internode length of branches | cm | Structural trait |
| $L_{IBMin}$ | Minimal internode length of branches | cm | Structural trait |
| $L_{ITMax}$ | Maximal internode length of trunk | cm | Structural trait |
| $L_{ITMin}$ | Minimal internode length of trunk | cm | Structural trait |
| $PH_{FO}$ | Number of first order branches arranged in a 360° circle | - | Structural trait |
| $P_{RU}$ | Pipe-reuse factor | - | Structural trait |
| $S_F$ | Shortening factor | - | Structural trait |
| $S_{Trop}$ | Strength of tropism | - | Structural trait |
| $ST_{Trop}$ | Stochasticity in tropism strength ($S_{Trop}$); only used if *Stochasticity*=1 | % | Structural trait |
| $ST_{Tw}$ | Stochasticity in branch growth; only used if *Stochasticity*=1 | % | Structural trait |
| $ST_{\alpha SFO}$ | Stochasticity in second order angle ($\alpha_{SFO}$); only used if *Stochasticity*=1 | % | Structural trait |
| $ST_{\alpha TFO}$ | Stochasticity in first order angle ($\alpha_{TFO}$); only used if *Stochasticity*=1 | % | Structural trait |
| $ST_{\alpha TSO}$ | Stochasticity in first order angle ($\alpha_{TSO}$); only used if *Stochasticity*=1 | % | Structural trait |
| $\alpha_{SFO}$ | Angle between first order branches and trunk from side view | ° | Structural trait |
| $\alpha_{TFO}$ | Angle between first order branches from top view | ° | Structural trait |
| $\alpha_{TSO}$ | Angle between second order branches and first order branches from top | ° | Structural trait |
| $\beta_D$ | Maximum relative increase in height growth when the $I_M < I_T$ | - | Structural trait |
| $\beta_S$ | Shape parameter regulating apical dominance of trunk apical meristem | - | Structural trait |

#### 1.3 Process overview and scheduling

At the beginning of each simulation, a pool of virtual species is generated with the number of species defined during experimental design. Each virtual species is characterized by a set of functional and structural traits (all species-specific traits are listed in Table S1). While the structural traits are uncorrelated and randomly selected from ranges defined during experimental design, functional leaf traits integrate between-trait correlations following the LES (Wright et al. 2004; Marino, Aqil and Shipley 2010). The LES quantifies relationships between leaf economic traits, such as the SLA, leaf lifespan or mass-based photosynthetic capacity. These leaf traits co-vary strongly and, in multidimensional trait space, the vast majority of variation is explained by a single principle axis (Wright et al. 2004). This axis can be considered as a spectrum, ranging from leaves with low SLA values, low photosynthetic capacities, and respiration rates, but long lifespans, to leaves with high SLA values, high photosynthetic capacities, and respiration rates, but short lifespans. To obtain a set of leaf traits, the values for SLA are randomly selected from ranges defined during experimentla design, and the values of the other traits are determined based on between-trait correlations, meaning that each set of leaf traits thus represents a position on the LES. Furthermore, functional traits characterizing specific photosynthetic light-response curves are also predicted from the leaf traits of the LES (following Marino, Aqil and Shipley 2010; Table S2).

After initialization, light distribution, tree establishment, tree growth, and tree mortality are simulated successively in annual time steps (Fig. S2). The 3D distribution of light intensity is calculated via the Lambert-Beer light extinction law considering the distribution of leaf area. Subsequently, establishment of tree seedlings is simulated as a neutral process. Depending on an average user-defined neutral germination rate (in number of seedlings per ha), new seedlings are initialized at random positions. Each seedling is randomly assigned to a species from the species pool. Seedlings with unsuitable traits for local voxel conditions may die due to carbon starvation.

Tree growth is simulated in three subsequent sub-processes: (I) apical control/dominance, (II) carbon balance, (III) structural growth (Fig. S2).

(I) Controlled by plant hormones, carbon allocation to apical meristems can either be inhibited (apical control) or intensified (apical dominance; Wilson 2000). These processes control how much of the carbon assimilated by photosynthesis is invested into primary growth of branches and the trunk. In this model, apical control is considered for branches. Branches inhibit carbon allocation to primary growth when branch apical meristems are 1) deeply shaded, i.e., if the carbon balance under local light conditions at the meristem is negative, or 2) when branches from neighboring trees grow into the same voxel. Hence, light and space competition takes place at the branch level. Apical dominance, in contrast, is considered at the trunk level, i.e., carbon allocation to trunk apical meristems is intensified under shade as a mechanism to quickly reach higher, potentially less shaded zones (Poorter 1999; Poorter et al. 2011). By influencing the within-tree carbon allocation, the processes of apical dominance/control slightly affect local carbon balance, the next tree growth subprocess.

(II) Local carbon balance is simulated at the level of leaf compartments. Apart from the usually only small percentage of carbon assimilated by leaf compartment and allocated to primary growth (influenced by apical control/dominance), leaf compartments are assumed to be independent from each other. Here, we follow the physiological principles of module autonomy stating that different tree parts may be regarded as autonomous modules (Sprugel et al. 1991). This means that no carbon flow occurs between leaf compartments and thus assimilated carbon is locally reinvested. Local re-investment means investments in leaf biomass within the leaf compartment, which, however, are coupled with investments in connected woody pipes following the pipe model theory (Shinozaki et al. 1964). For each new unit of leaf biomass an equivalent unit of pipes connecting leaves and roots has to be grown. Both construction and respiration costs for leaves and pipes are considered, which for the latter depend on the within-tree position of the leaf compartment. The carbon balance is central for the next and final tree growth sub-process – structural growth.

(III) In the structural growth sub-process, secondary and primary growth of branches and trunks is calculated. The sum of new pipes over all leaf compartments connected to a branch yields the branch diameter increment, and the sum over all leaf compartments of a tree yields the trunk diameter increment. Primary growth, in turn, is calculated based on the secondary growth using species-specific allometric relationships between diameter and length/height. Primary growth causes the establishment of new apical branch segments and sometimes new lateral branch segments, which may be associated with new leaf compartments and apical meristems. In addition, trees may also shed branches if all photosynthetically active leaf compartments are lost. After tree growth, tree mortality is simulated via different ecophysiological and environmental processes. Trees may die due to carbon starvation when they have lost all leaves. In addition, we integrated a biomass-dependent mortality rate according to metabolic theory of ecology (Brown et al. 2004). This rate accounts for processes not explicitly simulated (e.g., herbivory, pathogens) and assumes that the chance of survival increases non-linearly with tree biomass. Furthermore, gap dynamics are also important in tropical forests (Brokaw 1985). Therefore, falling dead trees may kill neighboring trees and create gaps. After each of the processes illustrated in Fig. S2, the state variables of all tree components are updated synchronously.

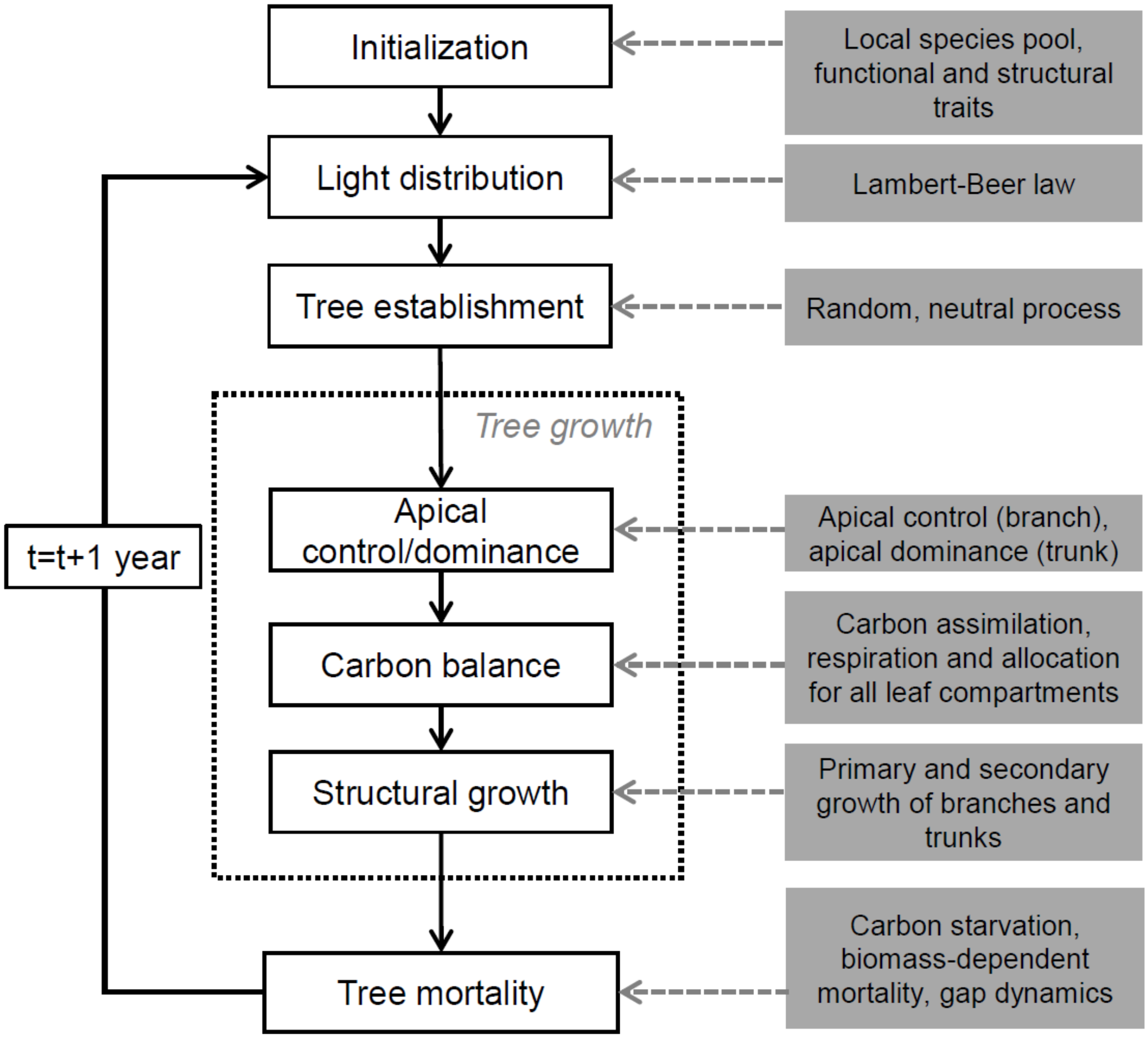


**Figure S2.** Flowchart of the model. After initialization, light distribution, tree establishment, growth and mortality are simulated consecutively in each annual time step. Tree growth is the most complex process and thus split into three submodels: apical control, carbon balance and structural growth. Details on all submodels are provided in in the section *Submodels* below.

#### 1.4 Design concepts

*Basic principles*

Carbon assimilation and allocation are the key processes in functional-structural tree and forest models. In our model, these processes are simulated based on the principles of the leaf economics spectrum (Wright et al. 2004), the pipe model theory (Shinozaki et al. 1964), and the principles of module autonomy (Sprugel et al. 1991). The LES quantifies relationships between crucial leaf economic traits, such as SLA, leaf lifespan or mass-based photosynthetic capacity. These leaf traits co-vary strongly and, in multidimensional trait space, the vast majority of variation is explained by a single principle axis (Wright et al. 2004). This axis can be considered as spectrum, ranging from leaves with low SLA values, low photosynthetic capacities, and respiration rates, but long leaf lifespans, to leaves with high SLA values, high photosynthetic capacities and respiration rates, but short leaf lifespans. The position on this spectrum thus has a direct influence on carbon assimilation and re-allocation at the leaf level. Furthermore, Marino, Aqil and Shipley (2010) observed that not only photosynthetic capacity, but also entire photosynthetic light-response curves can be predicted from the leaf traits of the LES. With this information, the carbon balance at the leaf level under varying light conditions can thus be simulated based on the specific leaf trait combination described by the position at the LES. However, the carbon assimilated by the leaves may be allocated among different tree parts, i.e., it may be invested into new leaves or branches at different within-tree positions. In this model, the within-tree carbon allocation is based mainly on the principles of module autonomy (Sprugel et al. 1991) and the pipe model theory (Shinozaki et al. 1964). The principles of module autonomy state that different parts of the tree may be regarded as autonomous modules whose carbon balance is independent of that of other modules. In our model, leaf compartments are these autonomous modules, which assimilate carbon based on their leaf traits and the local light intensity, and re-invest assimilates locally (Only a small exception from this rule is allowed in our model, as a small part of the assimilates in each leaf compartment is allocated for primary growth of the corresponding branch. There is, however, no carbon flow among leaf compartments). Local re-investment means investments in leaf biomass within leaf compartments which, however, are coupled with investments in connected woody pipes. In other words, for each new unit of leaf biomass an equivalent unit of pipes connecting leaves and roots has to be established, whereby the within-tree position of a leaf compartment determines the carbon costs for the pipes. New active pipes form the sapwood, which is equivalent to secondary or primary growth of branches and the trunk. By considering all leaf compartments of a tree, the whole-tree carbon balance and resulting structural growth can thus sufficiently be simulated based on the principles described above.

*Emergence*

Each tree is characterized by a set of traits, and structural tree growth, i.e., development, addition and removal of tree components, is a direct result of the interplay between these traits and light conditions. Hence, tree growth and tree mortality emerge from the traits of a tree. Some trait combinations might be unsuitable under low-light conditions and thus lead to carbon-based starvation. However, even under optimal conditions, each tree in the model will inevitably die at some point in time because it has lost all its photosynthetically-active parts. This is because the maximum height of each tree also emerges from its traits. When a tree grows close to its maximum height, it will enter senescence, which is characterized by the reduction of active meristems ultimately leading to the loss of all leaves (for more details see section *Structural growth*). Consequently, all crucial processes over the entire life cycle, as well as life expectancy itself, emerge directly from the functional and structural traits characterizing an individual tree.

While forest structure is the result of the growth of interacting and competing trees with different traits, community dynamics emerges from the trait-based mechanism at the tree level, as well as from tree establishment and additional source of tree mortality. The establishment rate defines how many new recruits enter the community, and different tree mortality rates are integrated to account for additional sources of mortality not captured by the FSTM.

*Adaptation/Sensing*

In this model, the interplay between the invariable functional and structural traits of trees and the dynamic environment determines their growth, but trees cannot adapt their traits to the environment. In reality, trait adaptations in response to environmental conditions may be observed within individuals. For instance, traits of sun and shade leaves might differ (Rozendaal et al. 2006, Markesteijn et al. 2007). However, in this model approach, we were more interested in interspecific trait differences than in trait differences within individuals

While adaptation and fitness-seeking of individual trees is not modelled explicitly at the trait level, we integrated two mechanisms controlling the primary growth of branches and trunks in dependence on the light conditions. The apical branch and trunk meristems sense their environment and, on this basis, either inhibit or intensify carbon allocation to the apical meristems. For branches, carbon allocation and thus primary growth is inhibited if the apical meristem senses insufficient light conditions or branches from neighboring trees in the immediate vicinity. This prevents carbon investments in tree parts with potentially low photosynthetic revenue. For trunks, primary growth is intensified under shade to reach higher, potentially less shaded zones faster. These apical control mechanisms can be understood as adaptation to the environment which may improve the fitness of the individuals.

*Interaction*

Both indirect and direct interactions among individuals are simulated. As the 3D light distribution is determined by the 3D leaf distribution in the community, competition for light is modelled as indirect interaction among the individuals. In contrast, crown development is directly influenced by competition for space between neighboring trees, because if trees sense tree components from neighboring trees in their immediate vicinity, they stop carbon allocation to this area. In addition, we integrated an option to simulate a direct feedback of falling trees on the mortality of neighboring trees, i.e., gap formation.

*Stochasticity*

The species pool containing the trait information of all local tree species is randomly drawn from user-defined ranges or estimated based on established between-trait correlations according to the LES (Wright et al. 2004). Tree establishment and mortality are also stochastic. The number of new seedlings at each time step can either be defined as a fixed value or as a range, from which the actual number is randomly chosen. Each new seedling is randomly distributed over the model area and a random species identity is assigned to it. Apart from trait-based cause of mortality (e.g., carbon starvation), we additionally integrated stochastic mortality: based on its current biomass, the mortality probability for each individual is estimated, and the decision whether to live or die is based on randomly drawn numbers. This additional mortality term covers sources of mortality which are not captured by the FSTM, such as infections by pathogens or excessive herbivory. Furthermore, if gap formation is simulated, trees die with a certain probability if large trees die nearby.

While the carbon balance of each individual tree is deterministic, the user can define if structural growth should be deterministic or stochastic. Deterministic structural growth means that trees strictly follow their structural model defined by their traits, i.e., branching angles are invariable and branches grow straight. Alternatively, stochastic structural growth can be switched on. In this case, individuals may randomly deviate from their regular structural growth within defined ranges and, as a consequence, branches grow irregularly. Choosing the stochastic structural growth model generates trees with a more natural and realistic appearance.

*Observation*

Emergent results can be monitored and saved at any hierarchical level (community, individuals, and tree components) at each time step from an omniscient perspective. Results at the community level include both stand variables and rates, such as the total above-ground biomass, the number of trees, total mortality rates or the net primary production. At the level of individual trees, aggregated variable such as the total tree height, crown width or height at first branching are recorded, while at the lowest hierarchical level, the state variables of all tree components are monitored (Table S1). As the amount of data at the low hierarchical levels can be enormous (a 1 ha forest stand may consist of several million tree components), we integrated the opportunity to select the time intervals at which the different model results are saved. Additionally, the graphical display of the simulated forest can be saved at each time step. More details on model outputs and options for customization are provided in the section *External model control, export and visualization*.

#### 1.5 Initialization

At the beginning of each simulation, a 3D grid space is initialized, whose spatial extent is defined by the parameters *MaxX*, *MaxY*, *MaxZ* and $L_{Cor}$. *MaxX* and *MaxY* define the core area in which trees can root, while $L_{Cor}$ defines the width of the corridor surrounding the core area in which trees may expand their crowns (Fig. S3). Cubic voxels of the grid space have a side length of $L_V$ and are clustered as 3D matrix (Fig. S1a). As the model space is initially empty, the total leaf area in all voxels is zero and thus the light intensity is at the global maximum $I_{max}$.

In addition, the species pool containing the trait information of $n_{Spec}$ species is initialized. For this purpose, the values of all structural traits for all species (Table S1) are randomly drawn from uniform distributions, whose minimum and maximum values are user-defined. In contrast, only two main functional traits characterizing the wood density ($\rho_W$) and the specific leaf area (*SLA*) are randomly drawn from uniform distributions with natural trait ranges. The additional functional traits are estimated based on correlations with these traits (Table S2) These correlations account for inevitable trade-offs, and the sets of species-specific leaf traits thus represent natural trait combinations.

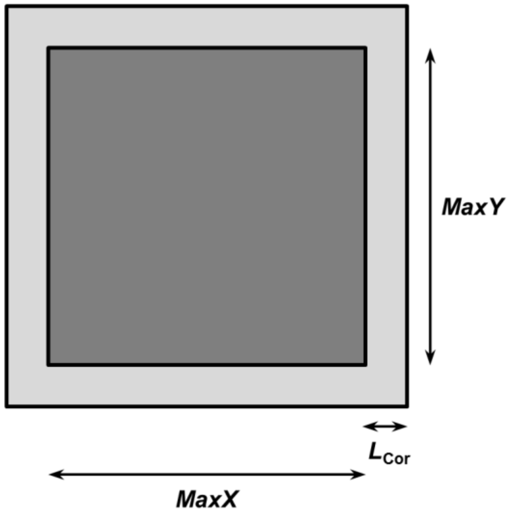


**Figure S3.** Top view on the model area. The core area in which trees can root is depicted in dark grey, the corridor in which trees may expand their crowns in light grey.

**Table S2.** Correlations between functional traits. The wood density $\rho_W$ and the specific leaf area SLA are the only traits which are freely chosen from defined ranges for each species. The leaf life span *LL* and foliar nitrogen concentration $N_{mass}$ are determined based on correlations with SLA following Wright et al. (2004). $R_L$, $G_{max}$ and $k$ are parameters of a hyperbolic Michaelis-Menten function determining the light response. Marino et al. (2010) found that these parameters are significantly correlated with the *SLA* and $N_{mass}$.

| Trait | Description | Trait value | Reference |
|---|---|---|---|
| *SLA* | Specific leaf area | Randomly selected from defined ranges | Wright et al. 2005, Patiño et al. 2012 |
| $\rho_W$ | Wood density | Randomly selected from defined ranges | Patiño et al. 2009, Quesada et al. 2012 |
| *LL* | Leaf lifespan | $LL = 30 \cdot 10^{-1.294+1.108 \cdot \log\left(\frac{10000}{SLA}\right)}$ | Wright et al. 2004 |
| $N_{mass}$ | Foliar nitrogen concentration | $N_{mass} = 10^{1.415-0.590 \cdot \log\left(\frac{10000}{SLA}\right)}$ | Wright et al. 2004 |
| $R_L$ | Leaf respiration rate | $R_L = 10^{3.06-1.01 \cdot \log\left(\frac{10000}{SLA}\right)}$ | Marino et al. 2010 |
| $G_{max}$ | Maximum gross photosynthetic rate | $G_{max} = 10^{3.71+0.47 \cdot \log(N_{mass})-0.85 \cdot \log\left(\frac{10000}{SLA}\right)}$ | Marino et al. 2010 |
| $k$ | Light intensity at which the gross photosynthetic rate is half maximal | $k_L = 10^{1.61-0.32 \cdot \log\left(\frac{10000}{SLA}\right)}$ | Marino et al. 2010 |

### 1.6 Input

The model does not use input data to represent time-varying processes.

#### 1.7 Submodels

In this section, all submodels (light distribution, tree establishment, tree growth, tree mortality; Fig. S2) are described in detail and chronologically. A list of all symbols, including explanations and units, is provided in Table S3.

##### 1.7.1 Light distribution

The 3D light environment is calculated based on the 3D leaf distribution. At first, the total leaf area in each voxel $A_{\mathrm{LTot}}{}^{\mathrm{XYZ}}$ is estimated based on the leaf area of leaf compartments within the particular voxel $A_{\mathrm{L}}{}^{\mathrm{XYZ}}$.

$$A_{LTot}{}^{XYZ} = \sum A_L{}^{XYZ} \tag{1}$$

Note that superscripts are used to indicate 3D positions. Second, based on the sum of $A_{\mathrm{LTot}}{}^{\mathrm{XYZ}}$ in all voxels above the specified voxel, the leaf area index $LAI^{\mathrm{XYZ}}$ for each voxel is calculated.

$$LAI^{XYZ} = \frac{\sum_{z}^{MaxZ} A_{Ltot}{}^{XYZ}}{L_V{}^2} \tag{2}$$

where $L_{\mathrm{V}}$ is the side length of a voxel. Assuming a Lambert-Beer extinction law, the single-column light intensity $I_{\mathrm{SC}}{}^{\mathrm{XYZ}}$ is calculated based on $LAI^{\mathrm{XYZ}}$.

$$I_{SC}{}^{XYZ} = I_{max} \cdot e^{-(k_L \cdot LAI^{XYZ})} \tag{3}$$

where $I_{\mathrm{max}}$ is the light intensity above the canopy and $k_{\mathrm{L}}$ the light extinction coefficient. This method assumes that solar radiation only penetrates directly from above and disregards additional processes like light reflection. This is an oversimplification, particularly in such heterogeneous forests as simulated here. To get a more realistic estimation of the average, effective light intensity within a voxel $I^{XYZ}$, the single column light intensity $I_{SC}{}^{XYZ}$ in the voxels surrounding the focal voxel in x and y direction are additionally taken into account. The number of surrounding voxels considered depends on the parameter *LR* which defines how many rectangular rings around the focal voxel are considered (Fig. S4). For each considered voxel, the relative contribution $C^{XYZ}$ is calculated, with $\Sigma C^{XYZ}=1$. The parameter $C^{XYZ}$ thus defines how strong $I_{SC}{}^{XYZ}$ in each voxel contributes to $I^{XYZ}$ and three different methods to calculate $C^{XYZ}$ defined by the parameter *LightC* can be applied: either (1) all voxels or (2) all rings contribute equally, or (3) the contribution of each voxel decays exponentially with distance from the focal voxel. On this basis, $I^{XYZ}$ is calculated as

$$I^{XYZ} = \sum_{X_{min}}^{X_{max}} \sum_{Y_{min}}^{Y_{max}} I_{SC}{}^{XYZ} \cdot C^{XYZ} \tag{4}$$

where $X_{min}=X\text{-}LR$ and $X_{max}=X+LR$ (likewise for $Y$; Fig. S4).

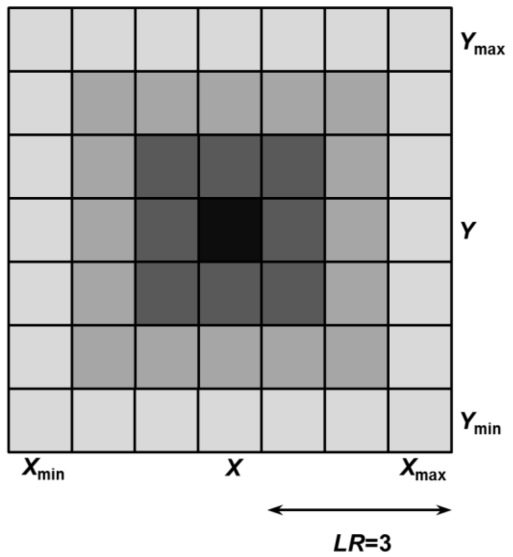


**Figure S4.** Illustration of voxels considered in calculations of effective light intensity. The light range *LR* defines how many rectangular rings of voxels (colored in grey shades) around the focal voxel (black) are considered. For each voxel (including the focal voxel), the relative contribution of the single-column light intensity to the effective light intensity of the focal voxel is calculated based on *LightC*. This parameter specifies whether each voxel or each ring contributes equally, or whether the contribution of each ring decreases exponentially with distance from the focal voxel. Adjacent voxels are only considered in X and Y direction, and not in Z direction.

As trees can only root in the core area but expand their crowns in the corridor, the total leaf area $A_{\mathrm{LTot}}^{\mathrm{XYZ}}$ decreases with distance from the forest edge, what increases the single-column light intensity $I_{\mathrm{SC}}^{\mathrm{XYZ}}$ at the corridor. Consequently, the effective light intensity $I^{\mathrm{XYZ}}$ also increases in the corridor or in the core area near the corridor. Such a pattern would resemble the light distribution in small forest fragments, whose edges permit light penetration. Because we were interested in also simulating pure forest core conditions, we integrated the possibility to choose between two options: small forest fragment ($EdgeC = 1$) or forest core ($EdgeC = 0$). In the latter case, $I^{\mathrm{XYZ}}$ would not be reasonably estimated in the vicinity of the edges of the model area and thus, periodic boundary conditions are applied. This means that the forest matrix surrounding the core area is similar to the forest in the core area and thus, before applying Eq. 4, the single-column light conditions $I_{\mathrm{SC}}^{\mathrm{XYZ}}$ calculated inside the core model are copied to the corridor in such a way that the conditions in the corridor resemble the conditions at the opposite side of the core area (Fig. S5). When the focal voxel for which $I^{\mathrm{XYZ}}$ is to be calculated is located near the edge of the entire model area (e.g., see voxel X2 in Fig. S5), not all adjacent voxels within the distance defined by $LR$ may exist. In this case, if periodic boundaries are specified, $I_{\mathrm{SC}}^{\mathrm{XYZ}}$ for these voxels can be obtained by strictly following the principles of periodic boundaries (Fig. S5). If real edge conditions are specified, $I_{\mathrm{SC}}^{\mathrm{XYZ}}$ for these voxels are obtained by mirroring $I_{\mathrm{SC}}^{\mathrm{XYZ}}$ at the outer border.

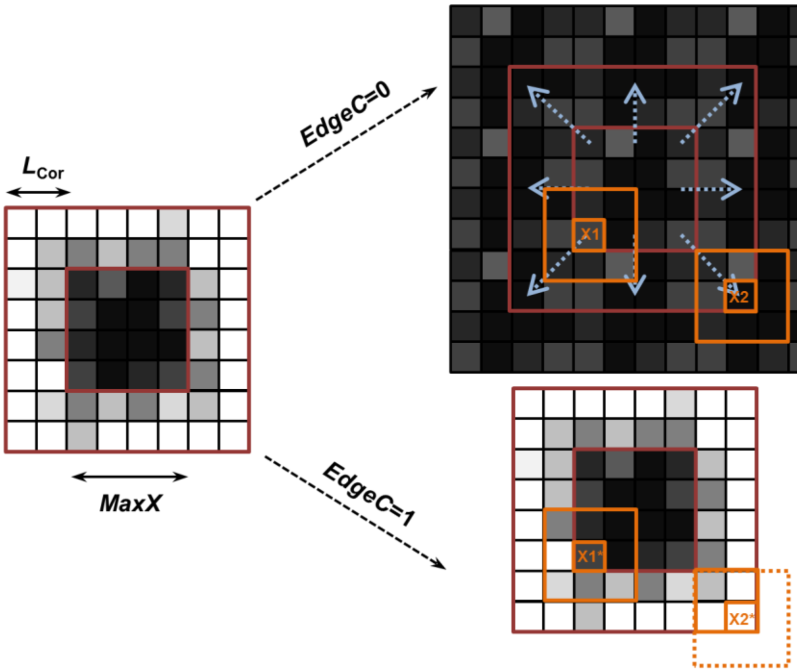


**Figure S5.** Illustration of principles applied when a small forest fragment (*EdgeC*=1) or a forest stand within a larger forest matrix (*EdgeC*=0) is simulated. The graph on the left side illustrates typical single-column light conditions for one horizontal voxel layer (darker colors represent lower light intensities). As trees only germinate within the core area, the single-column light conditions within the core area are typically higher compared to the corridor. If a small forest fragment (*EdgeC*=1) is simulated, the higher light intensity values in the corridor are used to calculate the effective light intensity (bottom right panel). When, as it is the case for the voxel X2*, not all surrounding voxel within *LR* (see Fig. S4) exists, voxel from inside are mirrored at the outer border and considered in light calculations. If a forest stand within a larger forest (*EdgeC*=0) is simulated, the single-column light conditions of the core area are first copied to the corridor before the effective light intensity is calculated (indicated by the blue arrows in top right panel; periodic boundary conditions).

##### 1.7.2 Tree establishment

Establishment is simulated as a neutral process, i.e., all species have the same probability of establishment. New seedlings are randomly distributed over the core model area (spatial resolution: 1 cm), whereby the total number of seedlings is controlled by the area-based establishment rate $n_{Seed}$. This rate can either be defined as a constant, or as a range from which the number of seedlings is randomly drawn at each time step. A randomly selected species ID from the species pool is assigned to each seedling, which is then initialized with the species-specific functional and structural traits (Table S1).

A seedling consists of a trunk with an apical meristem and an associated leaf compartment. Note that only at this seedling stage, leaf compartments are associated with the trunk. Thereafter leaf compartments are always associated with second order branches. The initial trunk diameter is given by $D_{ini}$. Because species differ in their intrinsic height-diameter relationships, the initial height is calculated based on $D_{ini}$ according to Eq. 6. Due to the relationship between leaf area and cross-sectional area of active pipes (Shinozaki et al. 1964), the initial leaf area is coupled to $D_{ini}$. Consequently, all seedlings start with a leaf compartment with equal initial leaf area $A_L$, but due to difference in *SLA*, the initial leaf biomass $B_L$ differs among species.

##### 1.7.3 Tree growth

Simulating tree growth using carbon-based FSTMs involves calculating carbon assimilation and allocation. Whereas the process of carbon assimilation is well-understood, the process of carbon allocation among different tree organs/components is debated (Lacointe 2000, Franklin et al. 2012). Several approaches to simulate carbon allocation have been proposed (Allen et al. 2005, Franklin et al. 2012, Mäkelä 2012). Here, we largely follow the principles of module autonomy, which state that plants are composed of repetitive modules which independently respond to their local environment (Sprugel et al. 1991, de Kroon et al. 2005). Hence, the assimilated carbon is reinvested locally into production of new leaves and branches (Sprugel et al. 1991). If light is unevenly distributed within canopies, module autonomy will create irregular tree crowns where the leaf biomass is mostly located in favorable, bright regions, which is a pattern often observed in nature.

In this model, the leaf compartments are the independent modules. Leaf compartments represent leaf-pipe elements attached to second order branches. While simulating the development of leaf compartments, all crucial processes (i.e., carbon assimilation, respiration, re-investment of surplus carbon) for both leaves and attached pipes are considered. Consequently, secondary growth of branches and the trunk emerge from the development of all connected leaf compartments. While secondary growth up to most distal branch junctions can sufficiently be simulated applying module autonomy in this model, the costs for primary growth are not explicitly accounted for. To account for theses carbon costs, leaf compartments would have to allocate a certain amount to the apical meristems for primary growth instead of re-investing it locally. This, however, means that leaf compartments cannot act as perfectly autonomous modules. Instead, a set of rules regulating carbon allocation among the different potential carbon sinks needs to be defined, which leads back to the initially stated problem concerning carbon allocation modeling methods.

Apart from the uncertainty which method to choose, carbon allocation models are commonly complex and thus computationally costly (Franklin et al. 2012). As model speed is a major constraint in this model, we chose not to use complex methods (e.g., maximization or optimization methods), but rather to simulate the carbon allocation to primary growth using a simple approximation method which largely keeps the autonomy of the leaf compartments. This means that we approximated the costs for primary growth based on the growth during the previous year and distribute these costs among all connected leaf compartments. We regard this approximation method as suitable trade-off between complexity and model speed, but in some situations this might not be appropriate. For instance, primary growth of branches predicted based on previous year growth might be overestimated if a branch collides with the crown of an adjacent tree or if its apical meristem is heavily shaded. In such situation, the apical meristem would commonly send the signal to cease or reduce carbon allocation for primary growth of this branch (King 1991, Stoll and Schmid 1998, Wilson 2000).

Thus, we integrated control mechanisms regulating primary growth based on the conditions of apical meristems (apical dominance/control). The carbon costs for primary growth according to these described approximation methods are estimated in the first submodel *Apical control*. For the sake of clarity, this step only approximates the costs that each leaf compartment contributes to primary growth, and not the actual primary growth, which is simulated thereafter. The relative costs for primary growth are usually small compared to the costs for new leaves and secondary growth and thus, approximating these costs seems sufficient. In the second submodel, the *Carbon balance* of all leaf compartments is calculated. This includes carbon assimilation, maintenance, and re-investment into new leaf and woody biomass. The change in leaf area/biomass and the leaf area/biomass production results from these processes. Secondary growth resulting from the carbon balance of all connected leaf compartments, as well as primary growth of all branches and the trunk is simulated in the submodel *Structural growth*.

##### *Apical control*

Each leaf compartment forms a leaf-pipe element whose pipe length $L_{PS}$ is calculated based on the relative position of the leaf compartment in the tree.

$$L_{PS} = P_{LC}{}^{Z} + \sqrt{(P_{LC}{}^{X} - P_{T}{}^{X})^2 + (P_{LC}{}^{Y} - P_{T}{}^{Y})^2} \qquad (5)$$

where $P_{LC}{}^{X}$, $P_{LC}{}^{Y}$ and $P_{LC}{}^{Z}$ are the spatial coordinates (centroids) of the leaf compartment, and $P_{T}{}^{X}$ and $P_{T}{}^{Y}$ are the coordinates of the trunk (we selected this simple method to approximate $L_{PS}$ for reasons of efficiency; calculating $L_{PS}$ based on the tree topology requires graph queries in

GroIMP, which are computationally demanding). By controlling the maintenance and construction cost of pipes, $L_{PS}$ influences the carbon balance of a leaf compartment. Now, we assume that not the entire carbon assimilated by a leaf compartment is locally re-invested, but that a certain proportion is allocated to the apical meristem of the connected first order branch and the trunk for primary growth. These additional costs for primary growth are taken into account by increasing the pipe length according to the predicted, potential length growth of the trunk and the first order branch. To predict the potential length growth in the current time step, we assume that the diameter increase equals the diameter increase in the previous year. On the basis of allometric relationships, the potential length increase can then be predicted:

$$L_T = 100 \cdot LD_T \cdot D_T^{2/3} \tag{6}$$

$$L_B = 100 \cdot LD_B \cdot {S_F}^{O_B} \cdot D_B^{2/3} \tag{7}$$

where $L_T$, $L_B$ is the length and $D_T$, $D_B$ is the diameter of trunks and branches, respectively. $LD_B$ and $LD_T$ are species-specific allometric shape parameter, with higher values representing more slender trunks or branches, and $S_F$ is a species-specific factor regulating the shortening of branches with their order $O_B$ ($S_F < 1$). The factor 100 converts m to cm as in such allometric relationships the diameter is generally given in cm and the length in m. These allometric relationships are based on McMahon (1973), who described that the critical length $L_{Cr}$ for buckling is proportional to the diameter $D$ raised to the 2/3 power.

$$L_{Cr} = 100 \cdot LD_{Cr} \cdot D^{2/3} = 100 \cdot 39 \cdot D^{2/3} \tag{8}$$

where $LD_{Cr}$ is the critical allometric shape parameter. This parameter is influenced by the ratio between wood density and elastic modulus, which is fairly constant in green wood, with the estimated values of $LD_{Cr}$=39 being regarded as upper limit across many tree species (McMahon

1973). Trees species commonly include stability safety factors, meaning that they grow below the critical length and hence $LD_B$ and $LD_T < LD_{Cr}$.

Based on Eqs. 6 and 7, the potential length increase can be predicted, but assuming that trunks and branches always grow according to allometric relationships might be too simplistic. Controlled by hormones, the allocation of carbon to apical meristems can either be inhibited (apical control; Wilson 2000) or intensified (apical dominance; Cline 1997), which modifies the shape of branches/trunks substantially. For instance, trees often intensify carbon allocation to the trunk apical meristem when they are shaded, most likely to quickly reach higher zones with more light (Poorter 1999). This process leads to more slender trunks. In contrast, branches commonly inhibit primary growth when their apical meristem is shaded or when branches collide. To account for these processes, we integrated additional mechanisms controlling the potential length increase.

For trunks, we assume that intensified carbon allocation to the trunk apical meristem is initiated when the light intensity at the apical meristem $I_M$ is below a species-specific threshold $I_T$. The relative intensification in height growth $L_{inc}$ (compared to the regular allometric growth) is implemented as function of the light intensity.

$$L_{inc} = \beta_D \cdot e^{-\left(\frac{I_M}{I_T}\right)^{\beta_S}} \tag{9}$$

where $\beta_S$ defines the shape of the function and $\beta_D$ is the maximum relative deviation. When considering $L_{inc}$, the potential length increase of the trunk $\Delta L_{TPotRg}$ is calculated as

$$\Delta L_{TPotRg} = LD_T \cdot \left(\left(2 \cdot D_{T(y0)} - D_{T(y-1)}\right)^{2/3} - \left(D_{T(y0)}\right)^{2/3}\right) \cdot (1 + L_{inc}) \tag{10}$$

where $D_{T(y0)}$ and $D_{T(y-1)}$ are the diameter at the beginning of the current time step and at the beginning of the previous time step, respectively. Continued apical dominance might lead to slender trunks which could potentially exceed the critical length (Eq. 8). Thus, the potential length increase up to the critical length $\Delta L_{TPotCr}$ is additionally estimated.

$$\Delta L_{TPotCr} = LD_{Cr} \cdot \left(2 \cdot D_{T(y0)} - D_{T(y-1)}\right)^{2/3} - L_{T(y0)} \qquad (11)$$

where $L_{T(y0)}$ is the length of the trunk at the beginning of the current time step. Each species has a maximum height $L_{TMax}$ resulting from its functional traits (Eqs. 41-42), which additionally limits the potential length increase. The potential length increase up to the maximum height $\Delta L_{TPotMax}$ is calculated as

$$\Delta L_{TPotMax} = L_{TMax} - L_{T(t)} \qquad (12)$$

Consequently, the effective potential length increase of the trunk $\Delta L_{TPot}$ is the minimum of these three variables.

$$\Delta L_{TPot} = \min(\Delta L_{TPotRg}, \Delta L_{TPotCrit}, \Delta L_{TPotMax}) \qquad (13)$$

For first order branches, we integrated two mechanisms regulating their potential length increase. First, branches stop to grow in length if the light intensity at the apical meristem is not sufficient to allow positive growth, i.e., if the photosynthetic rate $GR_{Pot}$<0 (see next section). Second, branches stop to grow in length if adjacent trees grow in the immediate surroundings, i.e., it is tested if there are any tree components from other trees in the same voxel as the apical meristem (this mechanism can be disabled by setting the global parameter *BrCollide*=0). In both cases, the potential length increase of branches is set to $\Delta L_{BPot}$=0. This means that branches may stop growing in length while continuing to grow in diameter, and thus they might deviate from their regular allometric relationship (Eq. 7). If, after a period of apical control, primary growth would be reactivated, for instance by more favorable light conditions, length increase would not be appropriately simulated based on Eq. 7, as branches could show an unrealistically huge increase in length in one time step. Thus, the potential length increase of branches $\Delta L_{\mathrm{BPot}}$, when not limited by low light or adjacent trees, is calculated as

$$\Delta L_{BPot} = LD_B \cdot S_F{}^{O_B} \cdot \left( \left(2 \cdot D_{B(y0)} - D_{B(y-1)}\right)^{2/3} - \left(D_{B(y0)}\right)^{2/3} \right) \quad (14)$$

This assumes that the increase in length at a given diameter can be approximated by the length increase if the branch would strictly have grown according to its regular allometric growth routine.

After the potential length increase of the first order branch and trunk associated with a leaf compartment has been calculated, the effective pipe length $L_{\mathrm{P}}$ of each leaf compartment is updated accordingly

$$L_P = L_{PS} + \Delta L_{TPot} + \Delta L_{BPot} \quad (15)$$

The effective pipe length $L_{\mathrm{P}}$ thus includes the pipe length of the leaf compartment according to its position within the tree plus the potential length increase of its associated first order branch $\Delta L_{\mathrm{BPot}}$ and trunk $\Delta L_{\mathrm{TPot}}$.

##### *Carbon balance*

This submodel simulates the carbon balance of all leaf compartments, which includes carbon assimilation and respiration, as well as loss of and investment into new biomass. When carbon assimilation exceeds the respiration/maintenance cost for leaves and connected pipes, the surplus carbon is invested into new leaf and pipe biomass. The sum of all leaf compartments of a tree comprises its total leaf and sapwood biomass.

To understand this submodel, we distinguish differences between voxels and leaf compartments. Voxels are not associated with any tree parts and contain aggregated information like the average light intensity (Eq. 4) or the total leaf area of all leaf compartments in the voxel (Eq. 1). If a new second order branch is generated within a voxel, a new leaf compartment is generated, which means that within the same voxel multiple leaf compartments may exit. Likewise, if a second order branch grows into a new voxel, a new leaf compartment is generated, which means that a leaf compartment is always associated with a *specific part* of a second order branch and consequently, each branch may have multiple leaf compartments.

While the model proceeds in annual time steps, many processes take place at shorter time intervals. For instance, new leaves may be produced during a year, which by increasing the photosynthetically active area positively influence the annual carbon balance. To better account for these effects, our model considers daily rates and simulates the development of the leaf compartments during one year. The annual rates are then estimated as the result of these simulations after $t_{\text{year}}$=360 days. An additional advantage of this approach is that seasonal forests can be simulated by reducing $t_{\text{year}}$.

Each leaf compartment contains leaves whose leaf dry mass $B_L$ and leaf area $A_L$ can be mutually converted via the species-specific SLA.

$$A_L = B_L \cdot SLA \qquad (16)$$

Leaves are the photosynthetically active organs and the gross carbon assimilation rate per unit of leaf dry mass $C_{\text{gross}}$ is calculated as hyperbolic Michaelis-Menten function.

$$C_{gross} = \frac{G_{max} \cdot I}{k + I} \cdot SI \qquad (17)$$

where $I$ is the light intensity at the leaf compartment (superscripts depicting spatial coordinates are not explicitly given in this section), and $G_{max}$ and $k$ are species-specific traits (Table S1). The site index $SI$ [0, 1] describes the relative environmental quality of the site and can be understood as aggregated information on all extrinsic factors which are not explicitly simulated in our model, e.g., nutrient, water availability or temperature. A $SI$ of 1 thus refers to optimum external factors and no resource limitation.

Maintenance costs have to be paid for both the leaves ($R_L$) and the sapwood, i.e., the pipes ($R_{WTot}$). While $R_L$ is a species-specific trait, the maintenance rate for connected pipes per unit of leaf dry mass $R_{WTot}$ depend on the position of the leaf compartment within the tree and are calculated as follows.

$$R_{wTot} = R_w \cdot \frac{L_P}{LP_{ratio}} \cdot \rho_W \cdot SLA \tag{18}$$

where $R_w$ are the general respiration costs per dry mass of pipes. Because we assume a fixed ratio between leaf area and cross-sectional area of connected pipes ($LP_{ratio}$), the total dry mass of pipes per unit of leaf dry mass can be calculated based on the length of the pipe system $L_P$, the wood density $\rho_W$ and the specific leaf area SLA.

Subtraction of the maintenance rates from the gross carbon assimilation rate yields the net carbon assimilation rate per unit of leaf mass $C_{net}$.

$$C_{net} = C_{gross} - R_L - R_{WTot} \tag{19}$$

If $C_{net}$ is positive, the surplus carbon can be reinvested into new leaf biomass and associated pipes. The amount of leaf dry mass that can be produced per unit of assimilated carbon $C_B$ depends on the ratio of leaf dry mass to pipe dry mass and can be calculated as follows.

$$C_B = \left( \left( CBL_{ratio} + \frac{L_P}{LP_{ratio}} \cdot CBW_{ratio} \cdot \rho_W \cdot SLA \cdot P_{RU} \right) \cdot C_O \right)^{-1} \quad (20)$$

While $C_{\mathrm{net}}$ is expressed in g carbon, the leaf and woody biomass is expressed in dry mass. Thus, the C-mass to biomass ratio of wood $CBW_{\mathrm{ratio}}$ and of leaves $CBL_{\mathrm{ratio}}$ is considered here. In addition, we assume that a certain proportion of C invested into new leaf or woody biomass is lost as growth respiration $C_{\mathrm{O}}$. $P_{\mathrm{RU}}$ [0, 1] is the pipe-reuse factor which specifies the ratio of new pipes to reused old pipes when new leaves are generated. When strictly following the pipe-model theory, new pipes are generated for each new leaf, while old pipes are converted from sapwood to heartwood when the leaves die (i.e., $P_{\mathrm{RU}}$=1). However, it is assumed that a certain proportion of old pipes can be reused (it is difficult to observe/measure this behavior, but see Mäkelä 1986, 2002). We thus added the possibility to include this mechanism ($P_{\mathrm{RU}}$<1). For the sake of clarity, $C_{\mathrm{B}}$ defines how much of the assimilated carbon is invested into leaf biomass considering the carbon costs for the pipes associated with the leaves. This means that, when calculating the total annual leaf biomass production based on $C_{\mathrm{net}}$ and $C_{\mathrm{B}}$, both the maintenance costs and the construction costs for leaves and pipes are fully included. From this it also follows that, due to the fixed leaf area to pipe area ratio ($LP_{\mathrm{ratio}}$), secondary growth is directly linked to the total annual leaf biomass production (next section).

Multiplication of $C_{\mathrm{net}}$ and $C_{\mathrm{B}}$ yields the relative growth rate of leaf biomass. Without considering leaf losses, the change in leaf biomass $B_{\mathrm{L}}$ over time in a leaf compartment could thus be described by the following ordinary differential equation.

$$\frac{dB_L}{dt} = C_B \cdot C_{net} \cdot B_L \quad (21)$$

It can be seen that, if $C_{net}$ is negative, leaf biomass is lost. In addition, as the average leaf lifespan $LL$ is an additional species-specific trait, leaves are constantly lost at a rate of $1/LL$. Thus, when considering both the (potential) production term ($C_{net}$ $C_B$) and the loss term (1/LL), the change in leaf biomass $B_L$ over time is

$$\frac{dB_L}{dt} = C_B \cdot C_{net} \cdot B_L - \frac{1}{LL} \cdot B_L = \left(C_B \cdot C_{net} - \frac{1}{LL}\right) \cdot B_L = GR_{pot} \cdot B_L \qquad (22)$$

Positive growth of leaf biomass is possible only if the carbon production rate is higher than the carbon loss rate (i.e., $GR_{pot}>0$). Solving this equation yields the leaf biomass as a general function of time.

$$B_L(t) = B_{L(t0)} \cdot e^{GR_{pot} \cdot t} \qquad (23)$$

where $B_{L(t0)}$ is the initial leaf biomass. Eq. 23 describes the temporal dynamics of leaf biomass by an exponential function to the base $e$, implying that surplus carbon is directly reinvested into new leaf biomass, which immediately participates in photosynthesis. However, in reality, surplus carbon is first allocated to leaf primordia, which develop into photosynthetically active organs with a time lag (Hallé et al. 1978). To account for this, we use the base 2 instead of $e$ in our simulations.

$$B_L(t) = B_{L(t0)} \cdot 2^{GR_{pot} \cdot t} \qquad (24)$$

As daily rates are used (Eqs. 17-19), the leaf biomass at the end of one year $B_{L(y+1)}$ can be calculated by inserting the number of suitable days $t_{year}$ and the initial biomass at the beginning of the year $B_{L(y0)}$.

$$B_{L(y+1)} = B_{L(y0)} \cdot 2^{GR_{pot} \cdot t_{year}} \qquad (25)$$

Equation 25 constitutes the basic rule to simulate the leaf biomass dynamics. Under sustained favorable light conditions, this equation would predict a potentially infinite accumulation of leaf biomass, which is not adequate as leaf compartments are limited by their discrete volumes (1 $m^3$). To get a more realistic behavior of the leaf biomass dynamics, two modifications are implemented. First, a global upper maximum of the total leaf area per voxel ($A_{LMax}$) is applied. Plants tend to avoid self-shading through efficient arrangements of leaf areas (King et al. 1997), and thus a maximal leaf area instead of a maximal leaf biomass is defined. Second, a species-specific maximum leaf production per leaf compartment ($A_{LProdMax}$) is implemented. The production of new leaves and branch segments is regulated by the activity of meristems, which generally follow specific intrinsic architectural rules (Hallé et al. 1978). Existing parts of branches do not have the potential to produce an unlimited number of new meristems capable of differentiating into leaves. $A_{LProdMax}$ can thus be understood as the maximum amount of leaves, expressed as leaf area, which can be produced within a leaf compartments associated with a specific section of a second order branch. As long as the total amount of leaves produced is lower than $A_{LProdMax}$, new leaf biomass can be produced if the light conditions are suitable. In the following, the modifications of the basic Eq. 25 under consideration of $A_{LMax}$ and $A_{LProdMax}$ are described. At first, to prevent the total leaf area in a voxel $A_{LTot}$ (Eq. 1) to exceed the maximum $A_{LMax}$, the theoretical maximal growth rate $GR_{max}$ is calculated so that $A_{LTot}$= $A_{LMax}$ when $GR_{max}$ is applied in Eq. 25 instead of $GR_{pot}$.

$$GR_{max} = \frac{log_2\left(\frac{A_{LMax}}{A_{LTot}}\right)}{t_{year}} \qquad (A_{LTot}, t_{year} \neq 0) \tag{26}$$

Note that the necessary conditions are always satisfied because naturally $t_{\text{year}}>0$ and, as leaf compartments without any leaf biomass are removed, for all existing leaf compartment $A_{\text{L}}>0$ and thus $A_{\text{LTot}}>0$. The effective growth rate $GR$ is then calculated as follows.

$$GR = \min(GR_{max}, GR_{pot}) \tag{27}$$

Integrating the effective growth rate $GR$ in Eq. 25 ensures that the total leaf area of all leaf compartments in a voxel never exceeds $A_{\text{LMax}}$. To ensure that the production maximum $A_{\text{LProdMax}}$ is never exceeded, it is essential to log the total leaf area production of a leaf compartment $A_{\text{LProdTot}}$.

$$A_{LProdTot(y+1)} = A_{LProdTot(y0)} + A_{LProd} \tag{28}$$

where $A_{\text{LProd}}$ is the annual leaf area production. Based on $A_{\text{LProdTot}}$ and $A_{\text{LProdMax}}$, the theoretical maximal leaf area production in the current time step $A_{\text{LProdTheo}}$ can be estimated.

$$A_{LProdTheo} = A_{LProdMax} - A_{LProdTot} \tag{29}$$

Dividing $A_{\text{LProdTheo}}$ by the SLA yields the theoretical maximal leaf biomass production in the current time step $B_{\text{LProdTheo}}$.

$$B_{LProdTheo} = \frac{A_{LProdTheo}}{SLA} \tag{30}$$

Consequently, it has to be verified whether the annual leaf biomass production, when applying the effective growth rate $GR$ (Eq. 27), would exceed this maximum. Please note that the annual leaf biomass production is not the same as the annual change in leaf biomass (Eq. 25), which is the result of leaf biomass production minus leaf loss. Thus, these two processes have to be separated. Using the effective growth rate $GR$, the potential leaf biomass production $B_{\text{LProdPot}}$ can be calculated as follows.

$$B_{LProdPot} = \begin{cases} \left(\frac{B_{L(y0)}}{GR} e^{GR \cdot t_{year}} - \frac{B_{L(y0)}}{GR}\right) \cdot \left(GR + \frac{1}{LL}\right) & if\ GR \neq 0 \\ B_{L(y0)} \cdot \frac{t_{year}}{LL} & if\ GR = 0 \end{cases} \quad (31)$$

where $B_{L(y0)}$ is the initial leaf biomass of the leaf compartment. The case discrimination is necessary because the regular equation to calculate $B_{LProdPot}$ is not defined if $GR$=0 (the additional necessary condition LL≠0 is always satisfied, as the leaf lifespan naturally is larger than zero). If $GR$=0, the leaf production rate must equal the leaf loss rate and consequently $B_{LProdPot}$ can be calculated based on the leaf loss rate 1/LL, which is a constant and species-specific rate. The effective leaf biomass production $B_{LProd}$ is simply calculated by applying the minimum function on $B_{LProdTheo}$ and $B_{LProdPot}$.

$$B_{LProd} = \min(B_{LProdPot}, B_{LProdTheo}) \quad (32)$$

Recapitulating, the effective leaf biomass production $B_{LProd}$ is the total amount of leaf biomass produced by a leaf compartment under consideration of $A_{LMax}$ and $A_{LProdmax}$. Now that the leaf biomass production for each leaf compartment is known, the change in leaf biomass equivalent to Eq. 25 has to be simulated. In Eq. 25, we assume that both the production rate and the loss rate are constant throughout the entire year. If the leaf production maximum $A_{LProdMax}$ would not be reached during a time step, which is the case if $B_{LProdTheo}$>=$B_{LProdPot}$, application of Eq. 25 would properly estimate the change in leaf biomass. However, if $A_{LProdMax}$ would be reached, i.e., if $B_{LProdTheo}$<$B_{LProdPot}$, Eq. 25 could not be applied. In this case, the production of new leaf biomass would stop during the year as soon as $A_{LProdMax}$ is reached. To account for this, we divide the year into two periods. In the first period, both leaf production and leaf loss are active and thus the leaf biomass dynamics can follow its regular mechanisms. In the second period, as soon as $A_{LProdMax}$ is reached, leaf production ceases and only leaf loss remains active. Based on the known leaf biomass production $B_{LProd}$ the length of the first period $t_p$, i.e., the 'productive time period', can be calculated as follows.

$$t_p = \begin{cases} \dfrac{ln\left(\dfrac{B_{LProd} \cdot GR}{B_{L(y0)} \cdot \left(GR + \dfrac{1}{LL}\right)} + 1\right)}{GR} & if\ GR \neq 0 \\ \dfrac{B_{LProd} \cdot LL}{B_{L(y0)}} & if\ GR = 0 \end{cases} \quad (33)$$

Under consideration of $t$p, the annual change in leaf biomass can be calculated as follows.

$$B_{L(y+1)} = B_{L(y0)} \cdot 2^{GR \cdot t_p} \cdot 2^{-\frac{(t_{year} - t_p)}{LL}} \quad (34)$$

This equation thus replaces the basic Eq. 25 and constitutes the final equation based on which the annual change in leaf biomass for each leaf compartment is calculated. This equation covers all possible scenarios. First, if there is no limitation in annual biomass production imposed by $A_{LProdMax}$, the productive time period becomes $t_P = t_{year}$ and thus Eq. 34 equals Eq. 25. Second, if $A_{LProdMax}$ is already reached at the beginning of the time step, i.e., if $B_{LProd}=0$, the productive time period becomes $t_P=0$. In this case, only leaf loss is considered in Eq. 34. Third, if $A_{LProdMax}$ is reached during the annual time step, the productive time period is estimated by Eq. 33 so that it exactly describes the time needed to reach $A_{LProdMax}$.

***Structural growth***

This submodel simulates the structural growth of trees and includes changes in the state variables of existing tree components, establishment of new tree components and removal of old ones. All of these processes result from the *Carbon balance*. The secondary growth results from the leaf biomass production in all topologically connected leaf compartments. The primary growth, in turn, is related to secondary growth via allometric relationships. Secondary and primary growth involve both changes in the state variables of existing tree components and the establishment of new ones. As we assume that photosynthetically inactive, leafless branches are shed, the removal of tree components is also a direct outcome of the *Carbon balance*. In the following, after an introduction to the modeling software GroIMP used here, we describe how the results of the preceding submodel are translated into structural growth. The structural traits are described in detail at the end of this section.

This model is implemented using the open-source software GroIMP (Growth Grammar Interactive Modeling Platform; available under the GNU General Public License at www.grogra.de). GroIMP is a 3D modeling platform suited to simulate the structural growth of plants. Here we illustrate the main concepts essential for understanding the functioning of this submodel (refer to Kniemeyer 2008 for detailed information on GroIMP). In GroIMP, relational growth grammars are implemented by the programming language XL, which is a graph-based extension of the Lindenmayer-Systems (L-Systems), a formal language for the description of plant structure (Lindenmayer 1968a, 1968b). XL is built on top of the programming language Java and thus both the XL-specific set of rules tailored to model plant structures, as well as the general Java classes can be used. Graphs are the underlying data structure in XL defining the tree topology. They describe how the different tree components of a tree, which can be defined as 3D geometric objects, are interconnected and spatially arranged to one another.

In our model, the trunk is defined as 3D cone, while the branch segments are defined as 3D cylinders. Taking into account the state variables of the tree components, the graph of each tree can thus be interpreted as 3D tree structure (Fig. S1c). XL contains a set of rules to modify the graph and thereby to induce structural growth. A rule consists of a *graph query*, an expression used to select specific parts of the graph, and a *statement* which specifies how to modify the selected parts. For example, a query may select all second order branches not connected to any leaf compartments, and a statement removes them. As another example, the rule to sum up the leaf biomass production over all leaf compartments topologically connected to a branch segment, and to change state variables of the branch segment accordingly, would, for each individual branch segment, traverse through the graph. Replacement rules are also common types of rules. Such rules select specific parts of a graph and replace them with other graph nodes, which, in this model, are the tree components. Meristems are the place of growth in trees and accordingly, apical meristems are replaced by other tree components to simulate primary growth in this model. Using the described rules, the results of the *Carbon balance* submodel are translated into structural growth in XL.

The first step is to calculate the updated total diameter of branches $D_{B(y+1)}$ and the trunk $D_{T(y+1)}$ based on the sum of the annual leaf biomass production $B_{LProd}$ of all topologically connected leaf compartments (Eq. 32). As the maintenance and construction costs of the pipes associated with leaf compartments have already been considered, the updated diameter is estimated using the ratio between leaf area and pipe cross-sectional area ($LP_{ratio}$).

$$D_{T(y+1)} = 2 \cdot \sqrt{\frac{\left(\left(\frac{D_{T(y0)}}{2}\right)^2 \cdot \pi + \frac{\sum B_{LProd}}{LP_{ratio}} \cdot SLA \cdot P_{RU}\right)}{\pi}} \tag{35}$$

$$D_{B(y+1)} = 2 \cdot \sqrt{\frac{\left(\left(\frac{D_{B(y0)}}{2}\right)^2 \cdot \pi + \frac{\sum B_{LProd}}{LP_{ratio}} \cdot SLA \cdot P_{RU}\right)}{\pi}} \tag{36}$$

Based on the updated diameter, the updated length of the trunks and branches can be calculated via allometric relationships (Eqs. 6-7), under consideration of the mechanisms of apical control/dominance. For branches, the potential length increase is set to $\Delta L_{BPot}$=0 either if their apical meristems are heavily shaded or if they collide with other trees (see submodel *Apical control*). On this basis, the updated branch length $L_{B(y+1)}$ is calculated as follows.

$$L_{B(y+1)} = \begin{cases} L_{B(y0)} + LD_B \cdot S_F{}^{O_B} \cdot \left( \left(D_{B(y+1)}\right)^{2/3} - \left(D_{B(y0)}\right)^{2/3} \right) & \Delta L_{BPot} \neq 0 \\ L_{B(y0)} & \Delta L_{BPot} = 0 \end{cases} \quad (37)$$

For trunks, no apical control mechanisms preventing length growth under unfavorable conditions are integrated. Rather, length growth can be intensified, and the relative intensification in height growth is expressed by $L_{inc}$ (Eq. 9; see submodel *Apical control* for details). The regular and otherwise unrestricted updated length of a trunk $L_{TRg}$ can thus be estimated according to Eq. 10 as follows.

$$L_{TRg} = L_{T(y0)} + LD_T \cdot \left( \left(D_{T(y+1)}\right)^{2/3} - \left(D_{T(y0)}\right)^{2/3} \right) \cdot (1 + L_{inc}) \quad (38)$$

While the environmental conditions at the trunk apical meristem do not limit height growth, it can be limited by the critical height $L_{Tcr}$ (Eq. 8) or the maximum trunk height $L_{TMax}$ (Eq. 42). Thus, the effective updated height of the tree $L_{T(y+1)}$ is calculated as follows.

$$L_{T(y+1)} = \min(L_{TRg}, L_{TCr}, L_{TMax}) \quad (39)$$

The critical length $L_{Tcr}$ is estimated based on $D_{T(y+1)}$ (Eq. 8). The maximum trunk height $L_{TMax}$ is a species-specific variable emerging from the functional traits, which is described in the following. A positive carbon balance in a leaf compartment can only be maintained if the carbon gain exceeds the carbon cost, i.e., if $GR_{Pot}$>0 (Eq. 22). The carbon gain generally increases with increasing light intensity $I$ (Eq. 17), while the carbon costs increase with the pipe length $L_P$ (Eqs. 18 and 20). At the theoretical maximal light intensity $I_{max}$, there is thus a maximum pipe length $L_{PMax}$ at which the carbon gain and the carbon costs are equal, i.e., at which $GR_{Pot}$=0.

$$GR_{Pot} = C_B \cdot C_{net} - \frac{1}{LL} = 0 \tag{40}$$

By substitution of Eqs. 19-20 into Eq. 40, and setting $I = I_{\max}$ and $L_{\mathrm{P}} = L_{\mathrm{PMax}}$, the maximum pipe length $L_{\mathrm{PMax}}$ can be calculated.

$$L_{PMax} = \frac{\frac{G_{max} \cdot I_{max} \cdot SI}{k + I_{max}} - R_L - \frac{CBL_{ratio} \cdot C_O}{LL}}{\frac{R_W \cdot \rho_W \cdot SLA}{LP_{ratio}} + \frac{C_O \cdot CBW_{ratio} \cdot \rho_W \cdot SLA \cdot P_{RU}}{LP_{ratio} \cdot LL}} \tag{41}$$

This equation contains only global constants and species-specific leaf and wood traits, making $L_{\mathrm{PMax}}$ an emergent species-specific variable. $L_{\mathrm{PMax}}$ thus represents the maximum pipe length under the given plot quality (i.e., site index *SI*), and the absolute maximum $L_{\mathrm{PMaxAbs}}$ can be estimated by setting $SI$=1.

As each tree is assumed to have only one trunk, the trunk length should never exceed $L_{\mathrm{PMax}}$. For branches, testing if the length of the pipe system exceeds $L_{\mathrm{PMax}}$ is not necessary, as this is implicitly done in the *apical control* submodel: if the carbon balance at the apical meristem would be negative ($GR_{\mathrm{Pot}}$<0), which is always the case if $L_{\mathrm{P}}$> $L_{\mathrm{PMax}}$, primary branch growth ceases (Eq. 37). However, if the maximum trunk height is defined as $L_{\mathrm{PMax}}$ and the apical control for branches is applied, the shape of trees can appear unrealistic. This is because $L_{\mathrm{PMax}}$ describes the theoretical, maximum pipe length at maximum light intensity, while the apical control of the branches considers the actual light intensity in the voxels. Hence, particularly when a trunk is close to $L_{\mathrm{PMax}}$, it might be that it continues to grow in length, while new lateral branches might not. In such a tree, it would appear as if the main trunk would grow through its own tree crown. To prevent this behavior, we introduce a safety factor for trunk growth *ST* (*ST*<1) that defines the ratio of the actual maximum trunk height to the theoretical maximum pipe length $L_{\mathrm{PMax}}$. The maximum trunk height is thus given as

$$L_{TMax} = ST \cdot L_{PMax} \quad (42)$$

While a trunk stops to grow in height at $L_{\mathrm{TMax}}$, lateral branches may grow above this point, by this creating realistic looking tree crowns (note that $ST$ is defined as a global constant and thus $L_{\mathrm{TMax}}$ remains a species-specific emergent trait).

While trunks are simply updated based on the updated state variables, updating the visual representation of branches is more complicated because branches are described at two scales. At the coarse scale, a branch is described by its total length and diameter as calculated above. At the fine scale, a branch is described by a set of topologically connected segments, which may further be connected to higher order segments. These branch segments at the fine scale are the tree components which are visually represented in GroIMP and thus, the state variables of the existing branch segments have to be updated and new branch segments have to be introduced according to the simulated total length and diameter growth. This means that in this model the total length growth of a branch is calculated first, resulting in establishment of a corresponding number of branch segments and not *vice versa* as in most FSTMs. We choose this two-scale approach as trade-off between computational costs and visual aspects. Treating the branch as an entity at the coarse scale reduced the computational cost by reducing the number of *graph queries*. If these coarse-scale branches would be visually displayed, the tree structure would appear unrealistic and consequently, we use smaller branch segments for visualization. This lends more realistic, irregular branch structures, including twisting of branches or effects of photo- or gravitropism (Fig. S6a). In the following, the essential information to calculate the fine-scale branch segments is provided.

Second order branches are the simplest case because they cannot ramify into higher order branches. The apical meristem of each second order branch is thus replaced by a segment with a length $L_S$ corresponding to the total length increase.

$$L_S = L_{B(y+1)} - L_{B(y0)} \tag{43}$$

The diameter of this new and all existing second order branch segments $D_S$ are updated based on their distance to their branch base $DI_S$ (Fig. S6a).

$$D_S = D_{B(y+1)} \frac{L_{B(y+1)} - DI_S}{L_{B(y+1)}} \tag{44}$$

The situation is more complex for first order branches because their primary growth might induce the establishment of new lateral branches. Thus, the number of lateral branches, as well as the length/diameter of both internodes and lateral branches, needs to be estimated (Fig. S6b). At first, the internode length $L_{IB}$, which defines the distance between two branching points, is calculated. In reality, the average internode length usually varies between species, but often also within individuals. A positive correlation between the total annual length growth and the internode length has been observed within individual trees (King 1997, Nicolini et al. 2003). On this basis, a flexible internode length $L_{IB}$ as function of total annual length growth, which can vary between the species-specific minimum $L_{IBMin}$ and maximum internode length $L_{IBMax}$, is used here.

$$L_{IB} = L_{IBMin} + \frac{(L_{IBMax} - L_{IBMin})}{1 + e^{(-k_{\mathrm{Int}} \cdot ((L_{B(y+1)} - L_{B(y0)} + L_{SLast}) - L_{IBMax} \cdot 2))}} \tag{45}$$

where $L_{SLast}$ is the lengths of the last apical branch segment (Fig. S6b) and $k_{Int}$ is a global constant controlling the change of $L_{IB}$. For clarity, $L_{IB}$ can differ between different branches of an individual tree, and obviously also from year to year, but for an individual branch we assume $L_{IB}$

to be invariable within one year. Based on $L_{\mathrm{IB}}$, the number of new lateral branches of a single branch $n_{\mathrm{BLat}}$ can thus be calculated.

$$n_{BLat} = floor(\frac{\left(L_{B(y+1)} - L_{B(y0)} + L_{SLast}\right)}{L_{IB}}) \quad (46)$$

Naturally, the number of new branch segments is $n_{\mathrm{BSeg}}$=$n_{\mathrm{BLat}}$+1. Since the total length growth $\Delta L_{\mathrm{B}}$ is usually not an integer multiple of the internode length $L_{\mathrm{IB}}$, the first and the last segment may be smaller than $L_{\mathrm{IB}}$ (Fig. S6b). The length of the first segment $L_{\mathrm{SFirst}}$ is estimated as follows.

$$L_{SFirst} = L_{IB} - L_{SLast} \quad (47)$$

where $L_{\mathrm{SLast}}$ refers to the last lateral segment of the previous year. The current $L_{\mathrm{SLast}}$ is estimated as

$$L_{SLast} = L_{B(y+1)} - L_{SFirst} - (N_{BLat} - 1) \cdot L_{IB} \quad (48)$$

This ensures that that the total length growth $\Delta L_{\mathrm{B}}$ equals the sum of the lengths of all new segments. To estimate the diameter of these branch segments $D_{\mathrm{S}}$, Eq. 44 is applied.

After the length and diameter of all new first order branch segments has been calculated, the length and diameter of all new lateral second order branches are estimated. For this, since the total length growth of the first order branch $\Delta L_{\mathrm{B}}$ is known, we first calculate the cross-sectional area $A_{\mathrm{Sec}}$ of the branch section representing this growth (Fig. S6c).

$$A_{Sec} = \left(\frac{D_{T(y+1)}}{2} \cdot \frac{L_{T(y+1)} - L_{T(y0)}}{L_{T(y+1)}}\right)^2 \cdot \pi \quad (49)$$

In compliance with the pipe model theory, the sum of the cross-sectional areas of all new lateral branches is assumed to equal $A_{\mathrm{Sec}}$. $A_{\mathrm{Sec}}$ is thus equally divided between all new lateral branches. This also means that we assume that all new lateral branches have the same size, i.e., we do not explicitly consider effects such as acrotony or mesotony. With $n_{BLat}$, the diameter of each new lateral branch $D_{BLat}$ is calculated as follows.

$$D_{BLat} = 2 \cdot \sqrt{\frac{A_{Sec}}{n_{BLat} \cdot \pi}} \tag{50}$$

Finally, the length of each new lateral second order branch $L_B$ is calculated based on the species-specific allometric diameter-length relationship (Eq. 7). We have to remember that branches are represented at two scales, and thus for each new branch, the total length and diameter as well as the segment lengths and diameters have to be calculated. As these new second order branches consist of a single segment, $L_S$=$L_B$ and $D_S$=$D_B$= $D_{BLat}$.

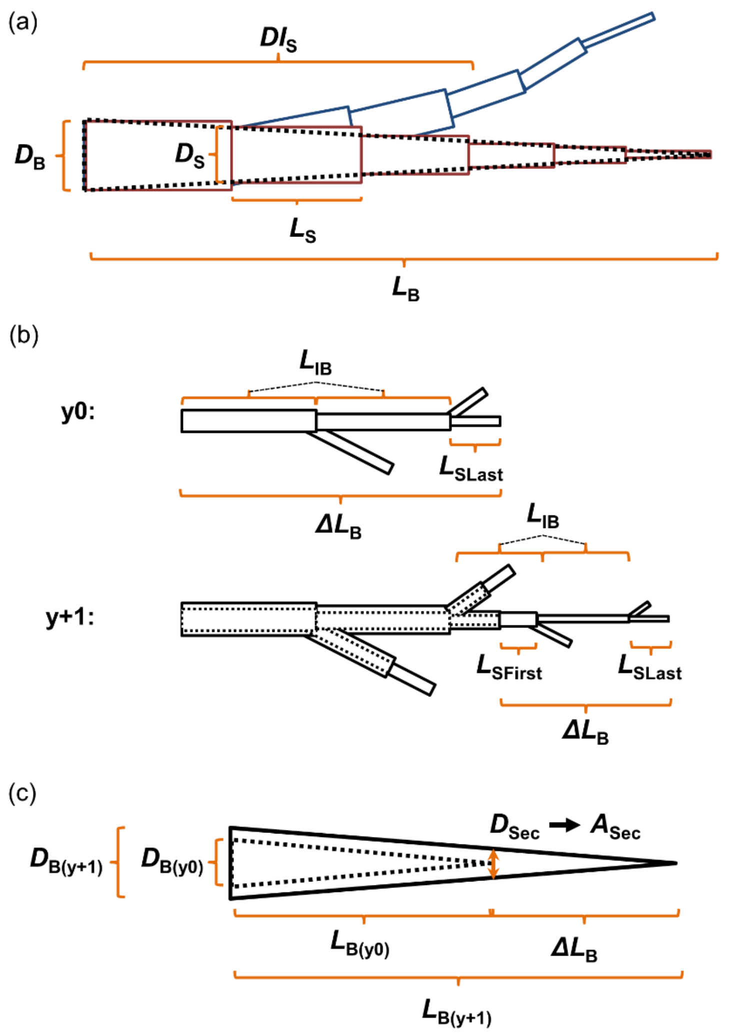


**Figure S6.** Illustration of structural variables. (a) Branches are represented at two different scales. At the coarse scale, branches are described by their total length $L_B$ and diameter $D_B$, while at the fine scale they are described as a collection of topologically connected smaller branch segments (length $L_S$, diameter $D_S$), which are visually represented by 3D cylinders (here: 2D representation). The distance of each branch segment to the branch base $DI_S$, which is exemplarily shown for the fourth branch segment, determines the diameter of the branch segment $D_S$. The fine-scale representation of branches allows a more realistic irregular visualization (blue colored branch as example). (b) Branch development in two successive years (y0 and y+1). The upper panel shows a newly created 1st order branch with lateral second order branch segments. The variable internode length $L_{IB}$ defines the length of the first two branch segments and the branching points between first and second order branches. Since the total length growth $\Delta L_B$ is not an integer multiple of the internode length $L_{IB}$, the length of the last lateral segment $L_{SLast}$ differs from $L_{IB}$. The lower panel shows the further development of this branch in the next time step. The internode length $L_{IB}$ in this time step may vary from that in the previous step, and because a shorter later lateral branch segment exists, and additional segment with a length of $L_{SFirst}$ is inserted so that distance between the branching points equals $L_{IB}$. (c) Branch growth at the coarse scale. Based on the diameter and length growth in one time step, the cross-sectional area $A_{Sec}$ of the branch section representing the current length growth $\Delta L_B$ can be calculated. $A_{Sec}$ is used to estimate the diameter of lateral branches.

So far, we have demonstrated how to calculate the state variables of branches at both scales when branches grow in length. However, when trunks grow in length, new lateral first order branches may establish, which, in turn, may ramify into second order branches. Trunks are not divided into separate segments, but nonetheless the internode length between two branching points at the trunk is an important information. The species-specific internode length of the trunk can be specified separately (minimum $L_{\mathrm{ITMin}}$, maximum $L_{\mathrm{ITMax}}$), and the method to calculate the variable internode length of trunks $L_{\mathrm{IT}}$ corresponds to that for branches (Eq. 45). $L_{\mathrm{IT}}$ thus defines the position at the trunk where to attach the new first order branch. When applying the methods describe above (Eqs. 43-50) in a recursive manner, all essential state variables of this first order branch and attached second order branches can be calculated.

The last step remaining is to update the diameter of all branch segments that already existed, which is done by applying Eq. 44. Obviously, the length of existing branch segments does not change.

After the structure of all woody tree components has been simulated, the remaining tree components, namely apical meristems and leaf compartments, need to be considered. Branches and trunks are always terminated by an apical meristem, and thus the structural growth simulations in GroIMP are carried out in a ways that this condition is true at all times. Each meristem is re-associated with the voxel in which it is located after the tree structure has been updated.

As a result of structural growth, new second order branches may be generated, and/or existing second order branches may grow into adjacent voxels. In these cases, new leaf compartments are associated with these branches and the initial leaf biomass is specified. We assume that the newly generated branches or branch sections consist entirely of sapwood and consequently, following the pipe model theory, their cross-sectional area and the leaf area of the associated leaf compartments are correlated via the parameter $LP_{\text{ratio}}$. For new second order branches, the cross-sectional area $A_B$ can thus be estimated from their known diameter, while for second order branches that increased in length, the cross-sectional area representing this growth $A_{Sec}$ can be calculated based on Eq. 49. Based on the cross-sectional area ($A_B$ or $A_{Sec}$) and the specific leaf area SLA, the total leaf area $A_{LSum}$ associated with the branch/branch section can be estimated.

$$A_{LSum} = \frac{A_B \cdot LP_{ratio}}{SLA} \tag{51}$$

For simplicity, we assume that $A_{LSum}$ is evenly divided among all new leaf compartments. For this, all new voxels a branch is intersecting with are estimated and in each voxel a new leaf compartment with the initial leaf biomass $B_{LInit}$ is generated.

$$B_{LInit} = \frac{A_{LSum}}{n_V} \tag{52}$$

where $n_V$ is the number of new voxels a branch is intersecting with.

Structural growth also includes the loss of existing tree components. Leaf compartments are lost if they no longer contain leaves (i.e., leaf biomass is zero). However, since the leaf biomass is simulated using an exponential function (Eq. 34), it would only converge to, but never reach zero, if the exponent is negative. Thus, we defined that leaf compartments are removed when the leaf biomass drops below a minimum threshold $B_{LMin}$. This threshold can be understood as the biomass of one leaf; the last leaf is thus dropped if $B_L < B_{LMin}$. If a leaf compartment is removed, it cannot be reestablished. This means that such leafless parts of a branch do not contain resting meristems.

Branches are shed if they lost all associated leaf compartments. This also implies that first order branches are shed when all connected second order branches are shed. Apart from this physiologically determined branch turnover, we also integrated the option to remove branches based on disturbances or mechanical stress. Branches may either be randomly removed (*BrMortMethod*=1) or based on their biomass (*BrMortMethod*=2). In the first case the branch mortality rate $m_{BR}$ defines the chance of a branch to be removed randomly at each time step, in the second case the branch mortality rate $m_B$ is calculated as follows.

$$m_B = m_{BB} \cdot \left(\frac{1}{3} \cdot \pi \cdot D_B \cdot L_B \cdot \rho_W\right)^{-M_{BS}} \tag{53}$$

where $m_{BB}$ is the biomass-based branch mortality rate, and the product within brackets is the mass of the branch, which is calculated by its diameter $D_B$, length $L_B$ and wood density $\rho_W$ assuming that it is cone-shaped. $M_{BS}$ is a scaling factor describing the decrease in mortality rate with increasing biomass (negative exponent). According to the metabolic theory, this scaling factor is assumed to be close to ¼ regarding the mortality of entire trees (Brown et al. 2004, Muller-Landau et al. 2006a). However, the scaling factor for branches may be site-specific and thus we integrated it as freely definable variable. Nevertheless, the user can choose to simulate only physiologically-determined branch fall (*BrMortMethod*=0).

In the previous description of structural growth, the structural traits were mentioned only briefly. The structural traits define how the different tree segments are spatially arranged, and thus they are required for a sufficient representation of the visible 3D tree structure. It was our intention to specify a minimal set of structural traits capable of reproducing the most obvious differences in tree structure observed in nature (Table S1; Fig. S7). The main structural traits and concepts are described hereafter.

The trunk is the orthotropic axis in each tree. First order branches are plagiotropic shoots that show a radial symmetry around the trunk. The angle between successive first order branches, seen from the bird's eye view, is calculated based on $Ph_{FO}$ which defines how many first order branches are arranged within a complete 360° circle (Fig. S7a). Thus, the angle between two successive first order branches $\alpha_{TFO}$ is calculated as

$$\alpha_{TFO} = \frac{360}{Ph_{FO}} \tag{54}$$

After a complete 360° circle, the successive first order branch is generated at an angle of ½ times $\alpha_{TFO}$ after its predecessor. This ensures that the branches do not directly shade the branches below. The angle of first order branches seen from the side is defined by $\alpha_{SFO}$ (Fig. S7b). In contrast to the first order branches, we assume that second order branches do not show a radial but rather a dorsiventral symmetry, i.e., they are arranged in the same plane as their mother branch. Their branching angle relative to the mother branch is defined by $\alpha_{TSO}$ (Fig. S7c). For simplicity, second order branches are always arranged in an alternating manner. As mentioned, the model differentiates between the internode lengths of branches $L_{IB}$ and trunks $L_{IT}$. The actual internode length at a given time step depends on the total growth of the specific branch or trunk and varies between the minimum ($L_{IBMin}$, $L_{ITMin}$) and maximum internode lengths ($L_{IBMax}$, $L_{ITMax}$), which are species-specific structural traits. Gravitropism or phototropism is often observed in trees: branches may bend downwards due to gravity and/or upwards to the sun. The strength of tropism $S_{Trop}$ is an additional functional trait, whereby negative values represent phototropism and positive values represent gravitropism.

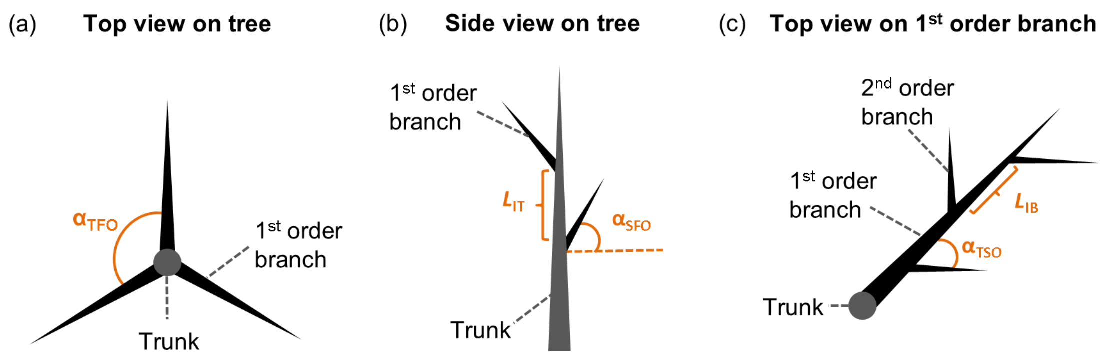


**Figure S7.** Illustration of the main structural traits. (a) Top view on tree showing the main trunk and first order branches. The angle between two consecutive first order branches is given by $\alpha_{TFO}$. (b) Side view on tree showing main trunk and first order branches. The trunk internode length $L_{IT}$ and the angle of the first order branch relative to the horizontal plane $\alpha_{SFO}$ define the coarse structure of the tree (c) Top view on tree showing one first and three second order branches. The branch internode length $L_{IB}$ and the angle between first order branch and second order branch $\alpha_{TSO}$ define the fine branching structure of the tree.

Differences in the mentioned structural traits create a variety of different tree structures. However, if a tree grows according to its structural traits in a deterministic manner, the resulting tree becomes too symmetrical. Thus, our model allows activating stochastic variation of structural traits if more realistic structures are desired (*Stochasticity*=0: deterministic growth, *Stochasticity*=1: stochastic growth). If stochastic growth is chosen, trees can deviate from their intrinsic structural growth pattern, whereby the strength of the random deviation is defined by a set of additional structural traits defining the maximum deviation of a specific species. For instance, the maximum deviation from $\alpha_{TSO}$ is given by $St_{\alpha TSO}$ and, in this case, the branching angle may thus vary within $\alpha_{TSO} \pm St_{\alpha TSO}$. The 'stochastic' structural traits can be understood as additional characteristic of species defining their structural irregularity. Such traits can be defined for all angles ($St_{\alpha TSO}$, $St_{\alpha TFO}$, $St_{\alpha SFO}$) and the tropism strength ($St_{Trop}$). In addition, $St_{TW}$ specifies the strength of branch twisting. This means that each time a new branch segment is generated, it may deviate from the direction of its predecessor by the three axis in space (head, left, up), whereby the maximal rotation along each axis is given by $St_{TW}$.

##### 1.7.4 Tree mortality

The metabolic theory of ecology generally predicts natural mortality rates to scale with biomass as the negative ¼ power (McCoy and Gillooly 2008). Muller-Landau et al. (2006b) tested this scaling relationship based on data from 10 old-growth tropical forest. They found large differences in the scaling factors between forest sites, which were mostly inconsistent with metabolic theory. However, at all sites the mortality rates consistently decreased with size when considering small trees < 20 cm diameter. For the larger trees, this trend differed and sometimes even reversed, i.e., the mortality rate of larger tree increased at some forest sites. Muller-Landau et al. (2006b) argued that there are additional site-specific mechanisms not explicitly considered in the metabolic theory.

In this model, there is only one explicit cause of mortality that directly emerges from the model approach, which is mortality due to carbon starvation. This happens when a tree has lost all its leaf compartments. The probability to lose a leaf compartment due to a negative carbon budget is higher in the dark understory compared to the upper forest zones. Therefore, the likelihood that a tree dies due to carbon starvation is higher for smaller trees and commonly decreases with size, which agrees with the pattern for small trees observed by Muller-Landau et al. (2006b). However, large trees growing in the canopy can also die due to carbon starvation in this model. When a tree grows close to its maximum height, it enters a phase of senescence where it loses more leaf biomass than it can produce, which ultimately leads to the loss of all leaf compartments and thus to carbon-based mortality. Consequently, the mortality rate due to carbon starvation might increase also for larger trees, explaining the trends observed by Muller-Landau et al. (2006b).

Nevertheless, it is unlikely that mortality due to carbon starvation is sufficient to capture all mortality mechanisms. We thus additionally integrated a mass-dependent mortality rate to account for additional causes of tree mortality, such as infections by pathogens or severe physical damages, which should scale with size. Due to the observed uncertainties in the scaling factor (Muller-Landau et al. 2006b), we integrated it as a free parameter $M_{TS}$. The biomass-dependent mortality rate $m_T$ is thus calculated as follows:

$$m_T = m_{TB} \cdot \left(\frac{1}{3} \cdot \pi \cdot D_T \cdot L_T \cdot \rho_W\right)^{-M_{TS}} \quad (55)$$

where $m_{TB}$ is the biomass-based tree mortality rate, and the product in the bracket is the mass of the tree trunk, which is calculated by its diameter $D_T$, length $L_T$ and wood density $\rho_W$ assuming that it is cone-shaped. This equation quantifies the probability of each tree to die, which decreases with biomass.

We further integrated the option to simulate mortality due to extrinsic factors, such as disturbances or gap formation. If the user intends to simulate disturbance events (*TrMortDist*=1), the average number of years between two events $F_{Dist}$ and the probability of the disturbance-mediated mortality $m_{Dist}$ are defined. If a direct effect of falling trees on neighboring trees mimicking gap formation shall be simulated (*TrMortNeigh*=1), the parameters $m_{Neigh}$ and $D_{NMin}$ have to be defined. We assume that only larger trees cause surrounding trees to break and die, and the minimum diameter of falling trees to be considered is given by $D_{NMin}$. The crown radius $CR_r$ of the falling trees defines the gap size, i.e., all smaller trees within distance $CR_r$ to the falling trees may die with a probability of $m_{Neigh}$. *Tree mortality* is the last submodel, thereafter the model proceeds with the next time step.

#### 2 External model control, export and visualization

This model is designed to be flexible and controlled by the user via simple text files. This allows manipulation and customization for simulation experiments without source code changes. There are two different types of text files, the *global* and the *pass* files.

The *global* file contains a set of parameters defining the basic set up of the model (Table S4). This includes the general decision whether a forest stand or an individual tree shall be simulated, the spatial extent and resolution of the model space, the number of time steps, and the number of replicates. Furthermore, the time intervals in which different types of model results are saved can be determined. The state variables of the tree components constitute the model results at the lowest hierarchical level, based on which higher level results are calculated. Users interested only in higher-level results can choose not to save the low-level results, or to save them in greater time intervals, by this reducing the required hard disk space (a 1 ha forest stand may consist of several millions branch segments). There are a total of six different types of result files: *Shoots*: state variable of tree components, *Trees*: tree level results, *Forest*: forest level results, *Species*: species pool, *Voxels*: leaf biomass and light in voxels, *Mortality*: number and causes of tree mortality. Which specific variable are saved in each of these files can be seen in Table S5. A short overview on important results is given in Fig. S8.

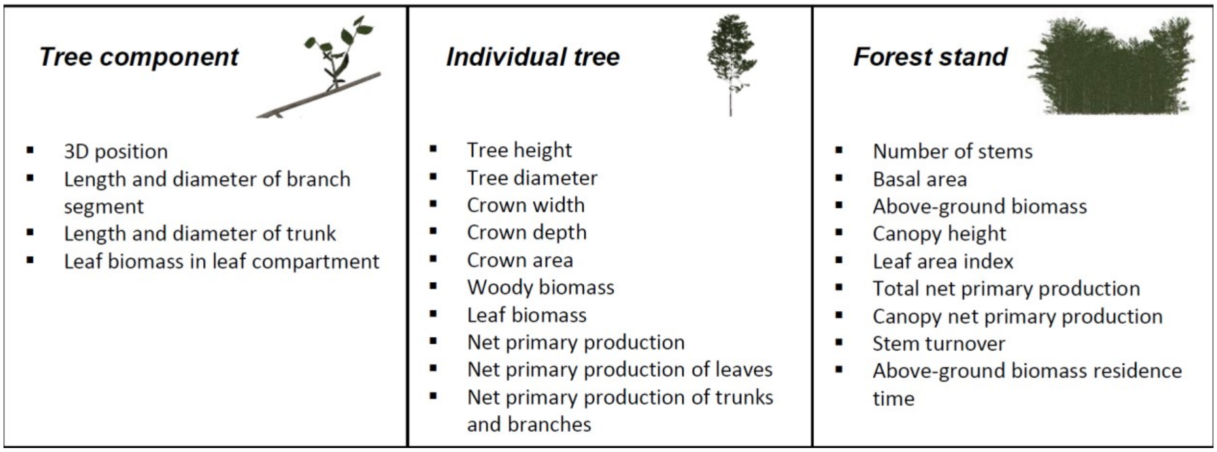


**Figure S8.** Exportable model results at the three hierarchical scales: tree component, individual tree and forest stand. The model allows saving model results as text files, and here, examples of important exportable variables are shown. A complete list of all exportable variables is provided in Table S5.

Visual control of simulated trees or forests is an important additional method to evaluate the quality of the model. Therefore, a picture showing the tree/forest structure is saved to disk at each time step. The perspective from which the picture is taken can be configured in GroIMP. Two general methods how trees are visualized are implemented, and they can be specified in the *global* file. First, trees can be represented by their woody components only (*VisualizationShader*=0), whereby second order branches connected to leaf compartments can be colored according to the state of the leaf compartment (Fig. S9). Second, trees can be represented by woody components and leaves (*VisualizationShader*=1). In our model we are not simulating individual leaves, however, for aesthetic purposed we integrated a technique which allows visually representing leaf compartments by leaf shaders (this technique is used in Fig. S1).

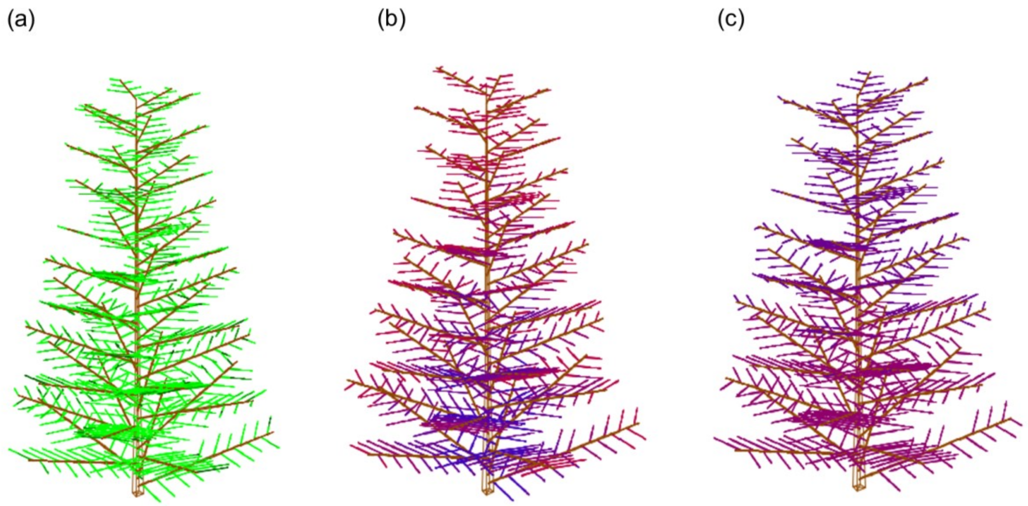


**Figure S9.** Visual representation of trees. If a wireframe model is chosen (*VisualizationShader*=0), three different methods to represent the leaf compartments attached to second order branches can be specified: (a) Second order branches are colored in different shades of green depending on the associated leaf biomass (*VisualizationMethod*=0). (b) Second order branches are colored according to the light conditions at the leaf compartments, with red colors representing high light intensities (*VisualizationMethod*=1). (c) Second order branches are colored according to the net carbon assimilation in the leaf compartments, with red colors representing higher values (*VisualizationMethod*=2). If a rendered representation is chosen (*VisualizationShader*=1), leaves representing the leaf compartments are visualized (see Fig. S1)

The *pass* files contain a set of parameters for each replicate, which means that the number of replicates specified in the global file and the number of pass files must be equal. Each pass file includes global parameters, ranges of functional and structural traits, but also parameters to switch on and off optional model mechanisms. An exhaustive list of all parameters in the pass file is available in Table S6.

### 3 Figures and Tables

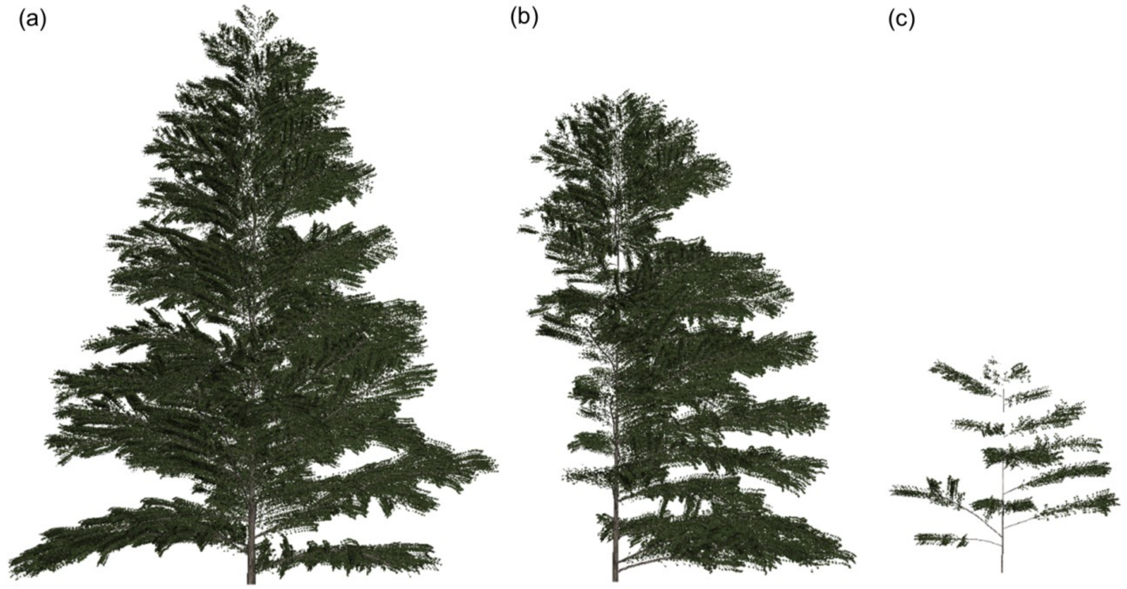


**Figure S10.** Effects of competition on tree growth. Three trees with identical functional and structural traits were simulated (a) without competition, (b) with one-sided competition and (c) with competition from 8 surrounding trees. The tree structures after 50 years of growth are displayed here.

1018 **Table S3.** List of abbreviations used in this study.

| Symbol | Explanation | Unit |
|---|---|---|
| $A_B$ | Cross-sectional area of branch | $cm^2$ |
| $A_{\mathrm{L}}$ | Leaf area in leaf compartment | $cm^2$ |
| $A_{\mathrm{LMax}}$ | Maximum leaf area in voxel | $cm^2$ |
| $A_{\mathrm{LProd}}$ | Leaf area produced in leaf compartment in one year | $cm^2$ |
| $A_{\mathrm{LProdMax}}$ | Absolute maximum leaf area production per leaf compartment | $cm^2$ |
| *ALProdMaxMethod* | Parameter specifying whether an invariable (*ALProdMaxMethod*=0) or a variable $A_{\mathrm{LProdMax}}$ is used (*ALProdMaxMethod*=1). | - |
| $A_{\mathrm{LProdTheo}}$ | Theoretical maximum annual leaf area production per leaf compartment | $cm^2$ |
| $A_{\mathrm{LProdTot}}$ | Total leaf area production in leaf compartment | $cm^2$ |
| $A_{LSum}$ | Total leaf area associated with growing branch | $cm^2$ |
| $A_{\mathrm{LTot}}$ | Total leaf area in voxel (per $m^2$) | $cm^2\ m^{-2}$ |
| $A_S$ | Cross-sectional area of branch segment | $cm^2$ |
| $A_{Sec}$ | Cross-sectional area of branch section | $cm^2$ |
| $A_T$ | Cross-sectional area of trunk | $cm^2$ |
| $B_{\mathrm{L}}$ | Leaf dry mass (in leaf compartment) | g |
| $B_{LInit}$ | Initial leaf biomass of new leaf compartment | g |
| $B_{\mathrm{LMin}}$ | Minimum leaf biomass below which leaf compartment is removed | g |
| $B_{\mathrm{LProd}}$ | Effective leaf biomass production in leaf compartment | g |
| $B_{\mathrm{LProdPot}}$ | Potential leaf biomass production when using the effective growth rate *GR* | g |
| $B_{\mathrm{LProdTheo}}$ | Theoretical maximum annual leaf dry mass production per leaf compartment | g |
| *BrCollide* | Parameter specifying whether branches stop to grow in length if the collide with surrounding trees (*BrCollide* =1) or not (*BrCollide* =0) | - |
| *BrMortMethod* | Parameter specifying whether branches are removed only if the lost all leaf compartments (*BrMortMethod*=0), or if they are additionally removed randomly (*BrMortMethod*=1) or based on their biomass (*BrMortMethod*=2) | - |
| *C* | Relative contribution of voxel in average light intensity calculations | - |
| $C_0$ | Carbon overhead costs | - |
| $C_B$ | Amount of leaf dry mass that can be produced per unit of assimilated carbon | $g\ gC^{-1}$ |
| $CBL_{\mathrm{ratio}}$ | C-mass to biomass ratio of leaves | $gC\ g^{-1}$ |
| $CBW_{\mathrm{ratio}}$ | C-mass to biomass ratio of wood | $gC\ g^{-1}$ |
| $C_{gross}$ | Gross carbon assimilation rate per unit of leaf dry mass | $gC\ g^{-1}\ d^{-1}$ |
| $C_{net}$ | Net carbon assimilation rate per unit of leaf dry mass | $gC\ g^{-1}\ d^{-1}$ |
| $CR_r$ | Crown radius of tree | cm |
| $D_{\mathrm{B}}$ | Diameter of branch | cm |
| $D_{\mathrm{BLat}}$ | Diameter of lateral branch | cm |
| $D_{\mathrm{ini}}$ | Initial diameter of seedling (fixed value) | cm |
| $DI_{\mathrm{S}}$ | Distance of branch segment to its branch base | cm |
| $D_{\mathrm{NMin}}$ | Trees with a diameter > $D_{\mathrm{NMin}}$ can create gaps (if *TrMortNeight=1)* | cm |
| $D_{\mathrm{S}}$ | Diameter of branch segment | cm |
| $D_{\mathrm{Sec}}$ | Diameter of branch section | cm |
| $D_{\mathrm{T}}$ | Diameter of trunk | cm |
| *EdgeC* | Parameter specifying whether a forest fragment with a real edge (*EdgeC*=1) or a forest patch within a forest matrix (*EdgeC*=0) is simulated | - |
| $F_{\mathrm{Dist}}$ | Frequency of disturbances (average number of years between two events) | a |
| $G_{\mathrm{max}}$ | Maximum gross photosynthetic rate (g C per g dry mass per day) | $gC\ g^{-1}\ d^{-1}$ |
| *GR* | Effective growth rate of leaf compartment | $d^{-1}$ |
| $GR_{\mathrm{max}}$ | Maximum growth rate of leaf compartment (considering $A_{\mathrm{LMax}}$) | $d^{-1}$ |
| $GR_{\mathrm{pot}}$ | Potential growth rate of leaf compartment | $d^{-1}$ |
| $h_{sun}$ | Assumed number of sun hours per day | h |
| *I* | Effective light intensity in voxel | $\mu mol\ m^{-2}\ s^{-1}$ |
| $I_{\mathrm{M}}$ | Light intensity at apical meristem | $\mu mol\ m^{-2}\ s^{-1}$ |

| | | |
|---|---|---|
| $I_{\mathrm{max}}$ | Light intensity above canopy | µmol $m^{-2}$ $s^{-1}$ |
| $I_{SC}$ | Single column light intensity | µmol $m^{-2}$ $s^{-1}$ |
| $I_{\mathrm{T}}$ | Light intensity threshold regulating apical dominance of trunk apical meristem | µmol $m^{-2}$ $s^{-1}$ |
| $k$ | Light intensity at which the gross photosynthetic rate is half of its maximum | µmol $m^{-2}$ $s^{-1}$ |
| $k_{int}$ | Factor controlling the relationship between internode length and total annual length growth | - |
| $k_{\mathrm{L}}$ | Light extinction coefficient (Lambert-Beer equation) | - |
| $LAI$ | Leaf area index | - |
| $L_{\mathrm{B}}$ | Length of branch | cm |
| $L_{Cor}$ | Width of corridor around core model area | m |
| $L_{Cr}$ | Critical length for buckling | cm |
| $LD_{\mathrm{B}}$ | Allometric parameter of length-diameter relationship of branches | - |
| $LD_{Cr}$ | Allometric parameter of length-diameter relationship (critical shape parameter) | - |
| $LD_{T}$ | Allometric parameter of length-diameter relationship of trunks | - |
| $L_{IB}$ | Branch internode length | cm |
| $L_{IBMax}$ | Species-specific maximum branch internode length | cm |
| $L_{IBMin}$ | Species-specific minimum branch internode length | cm |
| $LightC$ | Parameter specifying method to calculate average light intensity; $LightC$=[1,2,3] | - |
| $L_{\mathrm{inc}}$ | Relative intensification in height growth (apical dominance) | - |
| $L_{IT}$ | Trunk internode length | cm |
| $L_{ITMax}$ | Species-specific maximum trunk internode length | cm |
| $L_{ITMin}$ | Species-specific minimum trunk internode length | cm |
| $LL$ | Leaf lifespan | d |
| $L_{\mathrm{P}}$ | Length of pipe connected with leaf compartment (corrected after apical control) | cm |
| $L_{\mathrm{PMax}}$ | Maximum pipe length (under given $SI$) | cm |
| $L_{\mathrm{PMaxAbs}}$ | Maximum pipe length (theoretical maximum when $SI$=1) | cm |
| $LP_{\mathrm{ratio}}$ | Ratio between leaf area and pipe cross-sectional area | $cm^2$ $cm^{-2}$ |
| $L_{\mathrm{PS}}$ | Length of pipe connected with leaf compartment | cm |
| $LR$ | Maximal distance of surrounding voxels to be considered in light calculation | m |
| $L_{\mathrm{S}}$ | Length of branch segment | cm |
| $L_{SFirst}$ | Length of the first branch segment in each year | cm |
| $L_{SLast}$ | Length of the last branch segment in each year | cm |
| $L_{T}$ | Length of trunk | cm |
| $L_{TCr}$ | Critical trunk length | cm |
| $L_{TMax}$ | Maximum trunk length under consideration of $ST$ | cm |
| $L_{TRg}$ | Trunk length following unrestricted regular growth mechanism | cm |
| $L_{\mathrm{V}}$ | Side length of voxels | m |
| $MaxX$ | Spatial extent of core model area (in X direction) | m |
| $MaxY$ | Spatial extent of core model area (in Y direction) | m |
| $MaxZ$ | Spatial extent of core model area (in Z direction) | m |
| $m_{\mathrm{B}}$ | Biomass-based branch mortality rate (if $BrMortMethod$=2) | $a^{-1}$ |
| $m_{BB}$ | Parameter of biomass-based branch mortality rate (if $BrMortMethod$=2) | $g^{-1}$ $a^{-1}$ |
| $m_{\mathrm{BR}}$ | Random branch mortality rate (if $BrMortMethod$=1) | $a^{-1}$ |
| $M_{BS}$ | Scaling exponent in biomass-based branch mortality rate (if $BrMortMethod$=2) | - |
| $m_{\mathrm{Dist}}$ | Average relative mortality rate in a disturbance event (if $TrMortDist=1$) | $a^{-1}$ |
| $m_{\mathrm{Neigh}}$ | Trees affected by falling trees die with a probability of $m_{\mathrm{Neigh}}$ | $a^{-1}$ |
| $m_{\mathrm{T}}$ | Biomass-based tree mortality rate | $a^{-1}$ |
| $m_{TB}$ | Parameter of biomass-based tree mortality rate | $g^{-1}$ $a^{-1}$ |
| $M_{\mathrm{TS}}$ | Scaling exponent in biomass-based tree mortality rate | - |
| $n_{BLat}$ | Number of new lateral branches of a single branch in one time step | - |
| $n_{BSeg}$ | Number of new branch segments of a single branch in one time step | - |
| $N_{\mathrm{mass}}$ | Nitrogen concentration | % |

| | | |
|---|---|---|
| $n_{\mathrm{Seed}}$ | Number of seedlings dispersed at each time step | - |
| $n_{\mathrm{Spec}}$ | Number of species in species list | - |
| $n_{\mathrm{V}}$ | Number of new voxel a branch is intersecting with | - |
| $O_{\mathrm{B}}$ | Branch order | - |
| $P_{\mathrm{BEnd}}^{\mathrm{XYZ}}$ | End position of branch (in X, Y and Z direction) | cm |
| $P_{\mathrm{BStart}}^{\mathrm{XYZ}}$ | Start position of branch (in X, Y and Z direction) | cm |
| $Ph_{\mathrm{FO}}$ | Number of first order branches arranged in a 360° circle | - |
| $P_{\mathrm{LC}}^{\mathrm{XYZ}}$ | Position of leaf compartment (in X, Y and Z direction) | cm |
| $P_{\mathrm{M}}^{\mathrm{XYZ}}$ | Position of apical meristem (in X, Y and Z direction) | cm |
| $P_{RU}$ | Pipe-reuse factor | - |
| $P_{\mathrm{SEnd}}^{\mathrm{XYZ}}$ | End position of branch segment (in X, Y and Z direction) | cm |
| $P_{\mathrm{SStart}}^{\mathrm{XYZ}}$ | Start position of branch segment (in X, Y and Z direction) | cm |
| $P_{\mathrm{T}}^{\mathrm{XY}}$ | Position of trunk (in X and Y direction) | cm |
| $R_{\mathrm{L}}$ | Respiration rate per gram of leaf dry mass | gC $g^{-1}d^{-1}$ |
| $R_{\mathrm{w}}$ | Respiration rate per gram of sapwood | gC $g^{-1}$ $d^{-1}$ |
| $R_{\mathrm{WTot}}$ | Respiration rate of pipes per gram of leaf dry-mass | gC $g^{-1}$ $d^{-1}$ |
| $S_{\mathrm{F}}$ | Factor regulating the shortening of branches with their order | - |
| *SI* | Site index describing the relative quality of the forest patch | - |
| *SLA* | Specific leaf area | $cm^2$ $g^{-1}$ |
| *ST* | Safety factor for trunk growth. | - |
| *Stochasticity* | Parameter specifying whether stochastic variations of structural traits are simulated (*Stochasticity*=1) or not (*Stochasticity*=0) | - |
| $S_{\mathrm{Trop}}$ | Strength of tropism (negative values: phototropism; positive: gravitropism) | - |
| $\mathrm{St}_{\mathrm{Trop}}$ | Maximum deviation from $S_{\mathrm{Trop}}$ (if *Stochasticity*=1) | - |
| $\mathrm{St}_{\mathrm{Tw}}$ | Maximal rotation along the main growth axis (if *Stochasticity*=1) | ° |
| $\mathrm{St}_{\alpha\mathrm{SFO}}$ | Maximum deviation from $\alpha_{\mathrm{SFO}}$ (if *Stochasticity*=1) | ° |
| $\mathrm{St}_{\alpha\mathrm{TFO}}$ | Maximum deviation from $\alpha_{\mathrm{TFO}}$ (if *Stochasticity*=1) | ° |
| $\mathrm{St}_{\alpha\mathrm{TSO}}$ | Maximum deviation from $\alpha_{\mathrm{TSO}}$ (if *Stochasticity*=1) | ° |
| $t_{\mathrm{max}}$ | Number of simulated annual time steps | a |
| $t_p$ | Productive time period of leaf compartment during one year | d |
| *TrMortDist* | Parameter specifying if tree mortality due to disturbances is simulated (*TrMortDist=1)* | - |
| *TrMortNeigh* | Parameter specifying if tree mortality due to falling neighboring trees is simulated (*TrMortNeight=1)* | - |
| $t_{year}$ | Number of days per year suitable for photosynthesis | d |
| $\alpha_{\mathrm{SFO}}$ | Angle of first order branches from side view | ° |
| $\alpha_{\mathrm{TFO}}$ | Angle between first order branches from top view | ° |
| $\alpha_{\mathrm{TSO}}$ | Angle between second and first order branch from top view | ° |
| $\beta_{\mathrm{D}}$ | Maximum relative increase in height growth when $I_{\mathrm{M}} < I_{\mathrm{T}}$ | - |
| $\beta_{\mathrm{S}}$ | Shape parameter regulating apical dominance of trunk apical meristem | - |
| $\Delta L_{BPot}$ | Potential length increase of branch | cm $a^{-1}$ |
| $\Delta L_{TPot}$ | Effective potential length increase of trunk | cm $a^{-1}$ |
| $\Delta L_{TPotCr}$ | Potential length increase of trunk up to the critical buckling length | cm $a^{-1}$ |
| $\Delta L_{TPotMax}$ | Potential length increase of trunk up to the maximum tree height | cm $a^{-1}$ |
| $\Delta L_{TPotRg}$ | Potential length increase of trunk when considering apical dominance | cm $a^{-1}$ |
| $\rho_{\mathrm{W}}$ | Wood density | g $cm^{-3}$ |

**Table S4.** Parameters of the global file. The global file is a text file located in the main model folder and contains a set of parameters defining the basic set up of the model, such as the spatial extent of the number of time steps to be simulated.

| Parameter | Explanation | Unit | Symbol |
|---|---|---|---|
| *Timesteps* | Number of simulated annual time steps | a | $t_{max}$ |
| *Replicates* | Number of replicates to be simulated | - | - |
| *MaxX* | Spatial extent of core model area (in X direction) | m | *MaxX* |
| *MaxY* | Spatial extent of core model area (in Y direction) | m | *MaxY* |
| *MaxZ* | Spatial extent of core model area (in Z direction) | m | *MaxZ* |
| *WidthCorridor* | Width of corridor around core model area (only integer values allowed) | m | $L_{Cor}$ |
| *VoxelSize* | Side length of voxels (only integer values allowed) | m | $L_V$ |
| *ReportForest* | Time interval in which forest variables are saved | a | - |
| *ReportLight* | Time interval in which light variables are saved | a | - |
| *ReportMortality* | Time interval in which mortality variables are saved | a | - |
| *ReportShoots* | Time interval in which shoot variables are saved | a | - |
| *ReportTrees* | Time interval in which trees variables are saved | a | - |
| *ReportVoxel* | Time interval in which voxel variables are saved | a | - |
| *SimulateForest* | Parameter specifying whether a forest (*SimulateForest*=1) or an individual tree (*SimulateForest*=0) is simulated | a | - |
| *ThreadCount* | Number of threads that are used in parallel in light model | - | - |
| *VisualizationShader* | Parameter specifying whether rendered trees are shown (*VisualizationShader* =1) or not (*VisualizationShader* =0) | - | - |
| *VisualizationMethod* | Parameter specifying visualization method | - | - |

**Table S5.** Export parameters of the model. This table contains all parameters that are saved in each of the six different types of result files (Forest, Mortality, Shoots, Species, Trees and Voxels). The time interval at which each of this result files shall be saved to the hard disk can be defined by the user in the global file (Table S4).

| **Parameter** | **Explanation** | **Unit** | **File** |
|---|---|---|---|
| year | Year / time step | a | Forest |
| numberTrees | Number of trees | $ha^{-1}$ | Forest |
| basalArea | Basal area | $m^2\ ha^{-1}$ | Forest |
| maxHeight | Maximum tree height | m | Forest |
| meanHeight | Mean tree height | m | Forest |
| maxDiameter | Maximum tree diameter | m | Forest |
| meanDiameter | Mean tree diameter | m | Forest |
| woodyBiomass | Total biomass of all woody parts (trunks, branches) | $Mg\ ha^{-1}$ | Forest |
| trunkBiomass | Total trunk biomass | $Mg\ ha^{-1}$ | Forest |
| branchBiomass1stOrder | Total biomass of first order branches | $Mg\ ha^{-1}$ | Forest |
| branchBiomass2ndOrder | Total biomass of second order branches | $Mg\ ha^{-1}$ | Forest |
| leafBiomass | Total leaf biomass | $Mg\ ha^{-1}$ | Forest |
| trunkBiomassProduction | Total trunk biomass produced in one year | $Mg\ ha^{-1}\ a^{-1}$ | Forest |
| branchBiomass1stOrderProduction | Total biomass of first order branches produced in one year | $Mg\ ha^{-1}\ a^{-1}$ | Forest |
| branchBiomass2ndOrderProduction | Total biomass of second order branches produced in one year | $Mg\ ha^{-1}\ a^{-1}$ | Forest |
| leafBiomassProduction | Total leaf biomass produced in one year | $Mg\ ha^{-1}\ a^{-1}$ | Forest |
| branchBiomass1stOrderLoss | Total biomass of first order branches lost in one year | $Mg\ ha^{-1}\ a^{-1}$ | Forest |
| branchBiomass2ndOrderLoss | Total biomass of second order branches lost in one year | $Mg\ ha^{-1}\ a^{-1}$ | Forest |
| leafBiomassLoss | Total leaf biomass lost in one year | $Mg\ ha^{-1}\ a^{-1}$ | Forest |
| treeID | Tree ID | - | Mortality |
| speciesID | Species ID | - | Mortality |
| height | Tree height | m | Mortality |
| diameter | Tree diameter | m | Mortality |
| basalarea | Basal area of tree | m | Mortality |
| x | Position in core model area in X direction | m | Mortality |
| y | Position in core model area in Y direction | m | Mortality |
| age | Tree age | a | Mortality |
| causeDeath | Cause of death | - | Mortality |
| shootID | ID of branch segment | - | Shoots |
| branchID | ID of branch | - | Shoots |
| treeID | Tree ID | - | Shoots |
| speciesID | Species ID | - | Shoots |
| length | Length of branch segment | m | Shoots |
| diameter | Diameter of branch segment | m | Shoots |
| order | Branch order | - | Shoots |
| xbegin | Start position of branch segment (X direction) | m | Shoots |
| ybegin | Start position of branch segment (Y direction) | m | Shoots |
| zbegin | Start position of branch segment (Z direction) | m | Shoots |
| xend | End position of branch segment (X direction) | m | Shoots |
| yend | End position of branch segment (Y direction) | m | Shoots |
| zend | End position of branch segment (Z direction) | m | Shoots |
| SpeciesID | Species ID | - | Species |
| SLA | Specific leaf area | $cm^2\ g^{-1}$ | Species |
| rhoW | Wood density | $g\ cm^{-3}$ | Species |
| LL | Leaf lifespan | d | Species |
| Nmass | Nitrogen concentration | % | Species |
| RL | Respiration rate per gram of leaf dry mass | $gC\ g^{-1}d^{-1}$ | Species |
| Gmax | Maximum gross photosynthetic rate (g C per g dry mass per day) | $gC\ g^{-1}\ d^{-1}$ | Species |
| k | Light intensity at which the gross photosynthetic rate is half of its maximum | $\mu mol\ m^{-2}\ s^{-1}$ | Species |
| FirstOrderPhyllotaxis | Angle between first order branches from top view | ° | Species |

| | | | |
|---|---|---|---|
| FirstOrderPhyllotaxisNum | Number of first order branches arranged in a 360° circle | - | Species |
| FirstOrderAngleSide | Angle of first order branches from side view | ° | Species |
| HigherOrderAngle | Angle between second and first order branch from top view | ° | Species |
| InternodeLengthTrunkMin | Species-specific minimum trunk internode length | cm | Species |
| InternodeLengthTrunkMax | Species-specific maximum trunk internode length | cm | Species |
| InternodeLengthBranchMin | Species-specific minimum branch internode length | cm | Species |
| InternodeLengthBranchMax | Species-specific maximum branch internode length | cm | Species |
| kInt | Factor controlling the relationship between internode length and total annual length growth | - | Species |
| TropismStrength | Strength of tropism (negative values: phototropism; positive: gravitropism) | - | Species |
| LDRatioTrunk | Length-diamter ration of trunk | - | Species |
| ApicalDev | Maximum relative increase in height growth when IM < IT | | Species |
| IApical | Light intensity threshold regulating apical dominance of trunk apical meristem | $\mu$mol m$^{-2}$ s$^{-1}$ | Species |
| ShorteningFactor | Factor regulating the shortening of branches with their order | - | Species |
| maxPipeLength | Maximum pipe length of tree (emergent property) | cm | Species |
| StochasticityTwisting | Maximal rotation along the main growth axis (if *Stochasticity*=1) | ° | Species |
| StochasticityBranchingAngle | Maximum deviation from $\alpha_{TSO}$ (if *Stochasticity*=1) | ° | Species |
| StochasticityTropism | Maximum deviation from $S_{Trop}$ (if *Stochasticity*=1) | - | Species |
| StochasticityAnglePlane | Maximum deviation from $\alpha_{SFO}$ (if *Stochasticity*=1) | ° | Species |
| StochasticityPhyllo | Maximum deviation from $\alpha_{TFO}$ (if *Stochasticity*=1) | ° | Species |
| ALProdMax | Absolute maximum leaf area production per leaf compartment | cm$^2$ | Species |
| PipeReuseFactor | Pipe-reuse factor | - | Species |
| treeID | Tree ID | - | Trees |
| speciesID | Species ID | - | Trees |
| height | Tree height | m | Trees |
| diameter | Tree diameter | m | Trees |
| basalArea | Basal area of tree | m$^2$ | Trees |
| x | Position in core model area in X direction | m | Trees |
| y | Position in core model area in Y direction | m | Trees |
| age | Tree age | a | Trees |
| heightDelta | Height increase in one time step | m | Trees |
| heightRGR | Relative height increase in one time step | % | Trees |
| diameterDelta | Diameter increase in one time step | m | Trees |
| diameterRGR | Relative diameter increase in one time step | % | Trees |
| basalareaDelta | Basal area increase in one time step | m | Trees |
| basalareaRGR | Relative Basal area increase in one time step | % | Trees |
| woodyBiomass | Biomass of all woody tree parts (trunk and branches) | Mg | Trees |
| trunkBiomass | Biomass of trunk | Mg | Trees |
| branchBiomass1stOrder | Biomass of first order branches | Mg | Trees |
| branchBiomass2ndOrder | Biomass of second order branches | Mg | Trees |
| leafBiomass | Total leaf biomass of tree | g | Trees |
| leafArea | Total leaf area of tree | g | Trees |
| trunkBiomassProduction | Total trunk biomass produced in one year | Mg ha$^{-1}$ a$^{-1}$ | Trees |
| branchBiomass1stOrderProduction | Total biomass of first order branches produced in one year | Mg ha$^{-1}$ a$^{-1}$ | Trees |
| branchBiomass2ndOrderProduction | Total biomass of second order branches produced in one year | Mg ha$^{-1}$ a$^{-1}$ | Trees |
| leafBiomassProduction | Total leaf biomass produced in one year | Mg ha$^{-1}$ a$^{-1}$ | Trees |
| branchBiomass1stOrderLoss | Total biomass of first order branches lost in one year | Mg ha$^{-1}$ a$^{-1}$ | Trees |
| branchBiomass2ndOrderLoss | Total biomass of second order branches lost in one year | Mg ha$^{-1}$ a$^{-1}$ | Trees |
| leafBiomassLoss | Total leaf biomass lost in one year | Mg ha$^{-1}$ a$^{-1}$ | Trees |
| apicalLight | Light conditions at apical stem meristem | $\mu$mol m$^{-2}$ s$^{-1}$ | Trees |
| crownArea | Crown area of tree | m$^{-2}$ | Trees |

| | | | |
|---|---|---|---|
| crownWidth | Crown width of tree | m | Trees |
| crownDepth | Crown depth of tree | m | Trees |
| crownWidthRelative | Crown width relative to tree height | % | Trees |
| crownDepthRelative | Crown depth relative to tree height | % | Trees |
| heightFirstBranching | Height of first branching | m | Trees |
| x | Position of Voxel (in X direction) | m | Voxels |
| y | Position of Voxel (in Y direction) | m | Voxels |
| z | Position of Voxel (in Z direction) | m | Voxels |
| leafarea | Leaf area in Voxel | $cm^2$ | Voxels |

**Table S6.** Parameters of the pass file. The pass file is a text file located in the main model folder and contains a set of parameters for each replicate. Each pass file includes global parameters, ranges of functional and structural traits, but also parameters to select a specific optional model mechanism. The parameter values shown in this table are the values of the model shown in the main manuscript.

| Parameter | Explanation | Unit | Symbol | Value |
|---|---|---|---|---|
| LightC | Parameter specifying method to calculate average light intensity; *LightC*=[1,2,3] | - | *LightC* | 1 |
| ALMax | Maximum leaf area in voxel | $cm^2$ | $A_{LMax}$ | 15000 |
| ALProdMax (Min,Max) | Absolute maximum leaf area production per leaf compartment | $cm^2$ | $A_{LProdMax}$ | 65000 |
| LDTreeDev | Allometric parameter of length-diameter relationship of branches, LDT=LDB+LDRatioDev | - | $LD_T$ | -0.8/0.8 |
| BetaD (Min/Max) | Maximum relative increase in height growth when $I_M < I_T$ | - | $\beta_D$ | 0.1/0.3 |
| LightThreshApical | Light intensity threshold regulating apical dominance of trunk apical meristem | $\mu mol\ m^{-2}\ s^{-1}$ | $I_T$ | 30/100 |
| BetaS | Shape parameter regulating apical dominance of trunk apical meristem | - | $\beta_S$ | 3 |
| CarbonOverheadCosts | Carbon overhead costs | - | $C_0$ | 1.45 |
| CBLratio | C-mass to biomass ratio of leaves | $gC\ g^{-1}$ | $CBL_{ratio}$ | 0.5 |
| CBWratio | C-mass to biomass ratio of wood | $gC\ g^{-1}$ | $CBW_{ratio}$ | 0.5 |
| MortalityDisturbanceRate | Average relative mortality rate in a disturbance event (if *TrMortDist=1*) | $a^{-1}$ | $m_{Dist}$ | 0 |
| MortalityDisturbanceFrequency | Frequency of disturbances (average number of years between two events) | a | $F_{Dist}$ | 0 |
| AngleFirstOrderSideView (Min/Max) | Angle of first order branches from side view | ° | $\alpha_{SFO}$ | 0/40 |
| PhyllotaxisFirstOrder (Min/Max) | Number of first order branches arranged in a 360° circle | - | $Ph_{FO}$ | 3/5 |
| AngleSecondOrderTopView (Min/Max) | Angle between second and first order branch from top view | ° | $\alpha_{TSO}$ | 20/60 |
| Imax | Light intensity above canopy | $\mu mol\ m^{-2}\ s^{-1}$ | $I_{max}$ | 900 |
| InitialDiamter | Initial diameter of seedling (fixed value) | m | $D_{ini}$ | 0.0005 |
| InternodeLengthBranchMin (Min/Max) | Species-specific minimum branch internode length | m | $L_{IBMin}$ | 0.3/0.4 |
| InternodeLengthBranchMax (Min/Max) | Species-specific maximum branch internode length | m | $L_{IBMax}$ | 0.4/0.6 |
| InternodeLengthTrunkMin (Min/Max) | Species-specific minimum trunk internode length | m | $L_{ITMin}$ | 0.3/0.5 |
| InternodeLengthTrunkMax (Min/Max) | Species-specific maximum trunk internode length | m | $L_{ITMax}$ | 0.5/0.7 |
| KInt (Min/Max) | Factor controlling the relationship between internode length and total annual length growth | - | $k_{int}$ | 0.01/0.02 |
| LightExtinctionCoeff | Light extinction coefficient (Lambert-Beer equation) | - | $k_L$ | 0.6 |
| LDBranch | Allometric parameter of length-diameter relationship of branches | - | $LD_B$ | 3 |
| LPratio | Ratio between leaf area an pipe cross-sectional area | $cm^2\ cm^{-2}$ | $LP_{ratio}$ | 40000 |
| DistanceVoxelLightCal | Maximal distance of surrounding voxels to be considered in light calculation | m | *LR* | 4 |
| MinLeafBiomass | Minimum leaf biomass below which leaf compartment is removed | g | $B_{LMin}$ | 30 |
| MortalityBiomassRate | Parameter of biomass-based tree mortality rate | $g^{-1}\ a^{-1}$ | $m_{TB}$ | 0.032 |
| MortalityBiomassScalingExponent | Scaling exponent in biomass-based tree mortality rate | - | $M_{TS}$ | 0.13 |
| MortalityNeighMinDiameter | Trees with a diameter > $D_{NMin}$ can create gaps (if *TrMortNeight=1)* | cm | $D_{NMin}$ | 0.15 |
| MortalityNeighRate | Trees affected by falling trees die with a probability of $m_{Neigh}$ | $a^{-1}$ | $m_{Neigh}$ | 0.05 |
| RespirationRateWood | Respiration rate per gram of sapwood | $gC\ g^{-1}\ d^{-1}$ | $R_w$ | 0.0005 |
| NumberSeedlingPerHa (Min/Max) | Number of seedlings dispersed at each time step (per hectare) | $ha^{-1}\ a^{-1}$ | $n_{Seed}$ | 500/500 |
| SiteIndex | Site index describing the relative quality of the forest patch | - | *SI* | 0.9 |

| | | | | |
|---|---|---|---|---|
| SLA (Min/Max) | Specific leaf area | $cm^2$ $g^{-1}$ | *SLA* | 50/200 |
| NumberSpecies | Number of species in species list | - | $n_{Spec}$ | 1000 |
| Stochasticity | Parameter specifying whether stochastic variations of structural traits are simulated (*Stochasticity*=1) or not (*Stochasticity*=0) | - | *Stochasticity* | 1 |
| StochasticityTwisting (Min/Max) | Maximal rotation along the main growth axis (if *Stochasticity*=1) | ° | $St_{Tw}$ | 2/7 |
| StochasticityAngleSecondOrderTopView (Min/Max) | Maximum deviation from $\alpha_{TSO}$ (if *Stochasticity*=1) | ° | $St_{\alpha TSO}$ | 0/10 |
| StochasticityTropismStrength (Min/Max) | Maximum deviation from $S_{Trop}$ (if *Stochasticity*=1) | - | $St_{Trop}$ | 0/0.02 |
| StochasticityAngleFirstOrderSideView (Min/Max) | Maximum deviation from $\alpha_{SFO}$ (if *Stochasticity*=1) | ° | $St_{\alpha SFO}$ | 5/10 |
| StochasticityAngleFirstOrderTopView (Min/Max) | Maximum deviation from $\alpha_{TFO}$ (if *Stochasticity*=1) | ° | $St_{\alpha TFO}$ | 0/20 |
| StopCriterionBasalArea | Model stops and continues with next replicate if the total basal area exceeds $BA_{Stop}$ | $m^2$ $ha^{-1}$ | $BA_{Stop}$ | 80 |
| TropismStrength (Min/Max) | Strength of tropism (negative values: phototropism; positive: gravitropism) | - | $S_{Trop}$ | -0.02/0.02 |
| WoodDensity (Min/Max) | Wood density | g $cm^{-3}$ | $\rho_W$ | 0.5/0.7 |
| PipeReuseFactor (Min/Max) | Pipe-reuse factor | - | $P_{RU}$ | 0.6/0.6 |
| Tyear | Number of days per year suitable for photosynthesis | d | $t_{year}$ | 270 |
| Hsun | Assumed number of sun hours per day | h | $h_{sun}$ | 8 |
| TreeCompetionNum | Number of additional trees competing with tree (only if SimualteForest=0) | - | - | 0 |
| TreeCompetionDist | Distance of additional competing trees from tree (only if SimualteForest=0) | | - | 0 |
| BranchMortMethod | Parameter specifying whether branches are removed only if the lost all leaf compartments (*BrMortMethod*=0), or if they are additionally removed randomly (*BrMortMethod*=1) or based on their biomass (*BrMortMethod*=2) | - | *BrMortMethod* | 0 |
| BranchMortRandomRate | Random branch mortality rate (if *BrMortMethod*=1) | $a^{-1}$ | $m_{BR}$ | 0 |
| BranchMortMassRate | Parameter of biomass-based branch mortality rate (if *BrMortMethod*=2) | $g^{-1}$ $a^{-1}$ | $m_{BB}$ | 0.02 |
| BranchMortMassScalingExponent | Scaling exponent in biomass-based branch mortality rate (if *BrMortMethod*=2) | - | $M_{BS}$ | 0.2 |
| TreeMortNeigh | Parameter specifying if tree mortality due to falling neighboring trees is simulated (*TrMortNeight*=1) or not (*TrMortNeight*=0) | - | *TrMortNeigh* | 1 |
| TreeMortCarbon | Parameter specifying if tree mortality due to carbon starvation is simulated (*TrMortDist*=1) or not (*TrMortDist*=0) | - | *TrMortCarbon* | 1 |
| TreeMortDist | Parameter specifying if tree mortality due to disturbances is simulated (*TrMortDist*=1) or not (*TrMortDist*=0) | - | *TrMortDist* | 0 |
| PipeLengthMethod | Parameter specifying whether pipe length is calculated based on within-tree position (*PipeLengthMethod*=1) or based on height only (*PipeLengthMethod*=0) | - | - | 1 |
| FormFactorWood | Form factor used to calculate trunk biomass | - | - | 0.55 |
| SafetyFactorTrunk | Safety factor for trunk growth. | - | *ST* | 0.3 |
| EdgeC | Parameter specifying whether a forest fragment with a real edge (*EdgeC*=1) or a forest patch within a forest matrix (*EdgeC*=0) is simulated | - | *EdgeC* | 0 |
| BrCollide | Parameter specifying whether branches stop to grow in length if the collide with surrounding trees (*BrCollide* =1) or not (*BrCollide* =0) | - | *BrCollide* | 1 |

**Table S7.** Forest attributes in Neotropical forests based on a literature review. We concentrated on studies covering multiple forest plots or larger forest areas. When available, the number of 1 ha plots or the total study area is given in column ‘Extent’. If forest attributes were not estimated based on inventory data, this is mentioned in ‘Annotation’. Where available, means±sd are shown in column ‘Values’. Otherwise, means were estimated, for instance based on published maps. Values in brackets represent the range (min/max) of the forest attributes in the specific study, whereby extreme outliers were removed.

| Forest attribute | Unit | Value | Extent | Reference | Annotations |
|---|---|---|---|---|---|
| Above-ground biomass | Mg ha$^{-1}$ | ~280 | - | Mitchard et al. 2014 | Amazonia, Remote sensing |
| Above-ground biomass | Mg ha$^{-1}$ | ~330 | - | Mitchard et al. 2014 | Guiana Shield, Remote sensing |
| Above-ground biomass | Mg ha$^{-1}$ | ~270 | - | Mitchard et al. 2014 | SW Amazonia, Remote sensing |
| Above-ground biomass | Mg ha$^{-1}$ | ~280 (200/400) | n=82 | Malhi et al. 2015 | Amazonia |
| Above-ground biomass | Mg ha$^{-1}$ | 253 | n=28 | Banin et al. 2014 | Amazonia |
| Above-ground biomass | Mg ha$^{-1}$ | 195 (108/308) | n=35 | Feldpausch et al. 2012 | Brazilian Shield |
| Above-ground biomass | Mg ha$^{-1}$ | 344 (237/510) | n=44 | Feldpausch et al. 2012 | Eastern-Central Amazonia |
| Above-ground biomass | Mg ha$^{-1}$ | 434 (291/728) | n=45 | Feldpausch et al. 2012 | Guyana Shield |
| Above-ground biomass | Mg ha$^{-1}$ | 252 (142/392) | n=101 | Feldpausch et al. 2012 | Western Amazonia |
| Above-ground biomass | Mg ha$^{-1}$ | ~276 | n=20 | Baker et al. 2004 | NW Amazonia |
| Above-ground biomass | Mg ha$^{-1}$ | ~340 | n=17 | Baker et al. 2004 | Eastern-Central Amazonia |
| Above-ground biomass | Mg ha$^{-1}$ | ~246 | n=19 | Baker et al. 2004 | SW Amazonia |
| Above-ground biomass | Mg ha$^{-1}$ | ~300 (250/350) | n=227 | Malhi et al. 2006 | Amazon-wide interpolation |
| Above-ground biomass | Mg ha$^{-1}$ | 287.8±105.0 | n=33 | Slik et al. 2013 | Amazonia |
| AGB residence time | a | ~40 (20/100) | n=82 | Malhi et al. 2015 | Amazonia |
| AGB residence time | a | ~50 | n=127 | Galbraith et al. 2013 | Neotropics |
| AGB residence time | a | ~50 | - | Malhi et al. 2011 | Amazonia |
| AGB residence time | a | ~52 | - | Malhi et al. 2013 | W Amazonia |
| AGB residence time | a | ~80 | - | Malhi et al. 2013 | E Amazonia |
| Basal area | m$^2$ ha$^{-1}$ | 22.2±5.3 (7.1/32.4) | n=35 | Feldpausch et al. 2011 | Brazilian Shield |
| Basal area | m$^2$ ha$^{-1}$ | 23.5±10.2 (1.7/47.7) | n=44 | Feldpausch et al. 2011 | Eastern-Central Amazonia |
| Basal area | m$^2$ ha$^{-1}$ | 27.6±5.4 (16/37) | n=45 | Feldpausch et al. 2011 | Guyana Shield |
| Basal area | m$^2$ ha$^{-1}$ | 27.8±2.9 (15.6/39) | n=101 | Feldpausch et al. 2011 | Western Amazonia |
| Basal area | m$^2$ ha$^{-1}$ | 28.2 (21.7/36.8) | n=50 | Lewis et al. 2004a | Amazonia |
| Basal area | m$^2$ ha$^{-1}$ | ~26 (20/32) | - | Mitchard et al. 2014 | Amazonia, Remote sensing |
| Basal area | m$^2$ ha$^{-1}$ | 28.1 | n=28 | Banin et al. 2014 | Amazonia |
| Basal area | m$^2$ ha$^{-1}$ | 28.1±1.6 | n=15 | Chao et al. 2008 | North-Western Amazonia |
| Basal area | m$^2$ ha$^{-1}$ | 31.3±8 | n=9 | Chao et al. 2008 | North-Eastern Amazonia |
| Basal area | m$^2$ ha$^{-1}$ | ~28 (27/30) | n=20 | Laurance et al. 2009 | Central Amazonia |
| Basal area | m$^2$ ha$^{-1}$ | 29.9 | n=12 | Phillips et al. 1994 | Amazonia |
| Basal area | m$^2$ ha$^{-1}$ | ~27 (24/36) | n=20 | Baker et al. 2004 | NW Amazonia |
| Basal area | m$^2$ ha$^{-1}$ | ~29 (23/35) | n=17 | Baker et al. 2004 | Eastern-Central Amazonia |
| Basal area | m$^2$ ha$^{-1}$ | ~26 (20/30) | n=19 | Baker et al. 2004 | SW Amazonia |
| Basal area | m$^2$ ha$^{-1}$ | (25/31) | n=227 | Malhi et al. 2006 | Amazon-wide interpolation |
| Basal area growth | m a$^{-1}$ | 0.51 | n=50 | Lewis et al. 2004a | Amazonia |
| Basal area growth | m ha$^{-1}$ a$^{-1}$ | 0.68 (0.2/1.2) | n=28 | Banin et al. 2014 | Amazonia |
| Basal area growth | m ha$^{-1}$ a$^{-1}$ | 0.4/1.0 | n=104 | Malhi et al. 2004 | Amazonia |
| Canopy height | m | ~24 (15/37) | n=35 | Feldpausch et al. 2011 | South American dry forests (annual precipitation <1.5 m), height trees >40cm |
| Canopy height | m | ~31 (18/48) | n=44 | Feldpausch et al. 2011 | South American moist forests (annual precipitation 1.5-3.5 m), height trees >40cm |
| Canopy height | m | ~28 (18/38) | n=45 | Feldpausch et al. 2011 | South American wet forests (annual precipitation >3.5 m), height trees >40cm |
| Canopy height | m | 30.5±8.2 | n=101 | Helmer and Lefsky 2006 | Amazon river basin, lidar measurement |

| | | | | | |
|---|---|---|---|---|---|
| Canopy height | m | ~30 (25/40) | - | Simard et al. 2011 | Lidar measurement, Values for Amazon basin taken from published map |
| Canopy NPP | $Mg\ ha^{-1}\ a^{-1}$ | ~9 (6/12) | n=10 | Malhi et al. 2015 | Amazonia |
| Canopy NPP | $Mg\ ha^{-1}\ a^{-1}$ | ~10 | n=1 | Malhi et al. 2013 | Peru |
| Canopy NPP | $Mg\ ha^{-1}\ a^{-1}$ | ~5 | n=1 | Doughty et al. 2014 | Eastern Amazonia |
| Canopy NPP | $Mg\ ha^{-1}\ a^{-1}$ | 5.5/11.2 | n=10 | Aragão et al. 2009 | Amazonia |
| Canopy NPP | $Mg\ ha^{-1}\ a^{-1}$ | 6.6 (3/12) | n=33 | Malhi et al. 2011 | Neotropics |
| Canopy NPP | $Mg\ ha^{-1}\ a^{-1}$ | (6/13) | n=9 | Girardin et al. 2010 | Amazonia Lowland |
| Canopy NPP | $Mg\ ha^{-1}$ | ~7.5 (6.6-9.6) | n=3 | Malhi et al. 2009 | Amazonia |
| Mean dbh (>10cm) | cm | 27 | n=28 | Banin et al. 2014 | Amazonia |
| Mean dbh (>10cm) | cm | 20-22 | n=800 | Sawada et al. 2015 | Amazonia |
| Mean dbh (>10cm) | cm | 20-22 | n=14 | Lieberman et al. 1996 | Lowland Costa Rica |
| Number of Stems (>10cm) | $ha^{-1}$ | 551±110 (236/828) | n=35 | Feldpausch et al. 2011 | Brazilian Shield |
| Number of Stems (>10cm) | $ha^{-1}$ | 595±170 (153/927) | n=44 | Feldpausch et al. 2011 | Eastern-Central Amazonia |
| Number of Stems (>10cm) | $ha^{-1}$ | 515±99 (297/992) | n=45 | Feldpausch et al. 2011 | Guyana Shield |
| Number of Stems (>10cm) | $ha^{-1}$ | 559±74 (278/814) | n=101 | Feldpausch et al. 2011 | Western Amazonia |
| Number of Stems (>10cm) | $ha^{-1}$ | 581 (470/724) | n=50 | Lewis et al. 2004a | Amazonia |
| Number of Stems (>10cm) | $ha^{-1}$ | 589 | n=28 | Banin et al. 2014 | Amazonia |
| Number of Stems (>10cm) | $ha^{-1}$ | 595±23 | n=15 | Chao et al. 2008 | North-Western Amazonia |
| Number of Stems (>10cm) | $ha^{-1}$ | 560±35 | n=9 | Chao et al. 2008 | North-Eastern Amazonia |
| Number of Stems (>10cm) | $ha^{-1}$ | 645 (500/750) | n=12 | Phillips et al. 1994 | Amazonia |
| Number of Stems (>1cm) | $ha^{-1}$ | 2600/4700 | 50 ha | Chave et al. 2003 | Barro Colorado Island |
| Number of Stems (>1cm) | $ha^{-1}$ | 2000/2300 | n=8 | DeWalt and Chave 2004 | Neotropical lowland |
| Number of Stems (>1cm) | $ha^{-1}$ | ~4000 | n=3 | Muller-Landau et al. 2006b | Neotropical lowland |
| Stem turnover (>10cm) | $a^{-1}$ | ~2 (0.5/4) | n=97 | Phillips et al. 2004b | Pan-Amazon study |
| Stem turnover (>10cm) | $a^{-1}$ | 1.5± (0.5/3.5) | n=67 | Phillips 1996 | Amazonia, Mean tree turnover increased from ~1% to 2% from 1956-1991 |
| Stem turnover (>10cm) | $a^{-1}$ | ~1.8 (1.2/3.1) | n=50 | Lewis et al. 2004a | Amazonia |
| Stem turnover (>10cm) | $a^{-1}$ | 2.34±0.31 | n=15 | Chao et al. 2008 | North-Western Amazonia, mortality rate |
| Stem turnover (>10cm) | $a^{-1}$ | 1.21±0.53 | n=9 | Chao et al. 2008 | North-Eastern Amazonia, mortality rate |
| Stem turnover (>10cm) | $a^{-1}$ | (1/1.7) | n=20 | Laurance et al. 2009 | Central Amazonia |
| Stem turnover (>10cm) | $a^{-1}$ | ~1.85 (1.5/2.5) | n=12 | Phillips et al. 1994 | Amazonia |
| Stem turnover (>10cm) | $a^{-1}$ | (1/2) | n=14 | Lewis et al. 2011 | Amazonia |
| Stem turnover (>10cm) | $a^{-1}$ | (1.5/2.5) 95%, (1-5) | n=50 | Phillips et al. 2009 | Amazonia |
| Total ANPP | $Mg\ ha^{-1}\ a^{-1}$ | ~16 | 2 ha | Doughty et al. 2014 | Terra preta |
| Total ANPP | $Mg\ ha^{-1}\ a^{-1}$ | ~15 | n=3 | Malhi et al. 2009 | Amazonia |
| Total ANPP | $Mg\ ha^{-1}\ a^{-1}$ | 10/22 | n=10 | Aragão et al. 2009 | Amazonia |
| Total ANPP | $Mg\ ha^{-1}\ a^{-1}$ | 10/20 | n=9 | Girardin et al. 2010 | Amazonia Lowland |
| Total ANPP | $Mg\ ha^{-1}\ a^{-1}$ | ~12.9 | 21 ha | Chambers et al. 2001 | Amazonia Lowland |
| Leaf area index | $m\ m^{-2}$ | ~7 | - | Asner et al. 2004 | Amazonia (mean) |
| Leaf area index | $m\ m^{-2}$ | ~8 | - | Caldararu et al. 2012 | Central and Southern Amazonia; Satellite Observations |
| Leaf area index | $m\ m^{-2}$ | ~0 | - | Caldararu et al. 2012 | Eastern Amazonia; Satellite Observations |
| Leaf area index | $m\ m^{-2}$ | ~7 | - | Myneni et al. 2007 | Amazonia (mean) |

**Table S8.** Value ranges of model parameters. Many parameters used in the present model have natural ranges, which were estimated based on literature values.

| Symbol | Unit | Explanation | Value (range) | References |
|---|---|---|---|---|
| $A_{\mathrm{LMax}}$ | $cm^2$ | Maximum leaf area in voxel | 10000-20000 | Model specific parameter, own estimates |
| $A_{\mathrm{LProdMax}}$ | $cm^2$ | Absolute maximum leaf area production per leaf compartment | 40000-80000 | Model specific parameter, own estimates |
| $LD_{\mathrm{T}}$ | - | Allometric parameter of length-diameter relationship of branches, LDT=LDB+LDRatioDev | -1-1 | McMahon 1973, Bertram 1989, Niklas 1995, West, Brown and Enquist 1999, van Gelder, Poorter and Sterck 2006, Banin et al. 2012 |
| $C_0$ | - | Carbon overhead costs (construction costs) | 1.4-1.5 | Poorter and De Jong 1999, Cannell and Thornley 2000, Sterck et al. 2005, Pons and Poorter 2014 |
| $CBL_{\mathrm{ratio}}$ | $gC\ g^{-1}$ | C-mass to biomass ratio of leaves (carbon content) | 0.45-0-5 | Houghton et al. 2001, Elias and Potvin 2003, Martin and Thomas 2011 |
| $CBW_{\mathrm{ratio}}$ | $gC\ g^{-1}$ | C-mass to biomass ratio of wood (carbon content) | 0.45-0.5 | Houghton et al. 2001, Elias and Potvin 2003, Martin and Thomas 2011 |
| $I_{\mathrm{max}}$ | $\mu mol\ m^{-2}\ s^{-1}$ | Light intensity above canopy | 700-1200 | Chazdon and Fetcher 1984, Berry, Varney and Flanagan 1997, Valladares, Allen and Pearcy 1997, Sterck et al. 2011, Seyoum et al. 2014 |
| $k_{\mathrm{L}}$ | - | Light extinction coefficient (Lambert-Beer equation) | 0.5-0.8 | Huth and Ditzer 2000, Kitajima, Mulkey and Wright 2005b, Malhi et al. 2013 |
| $LD_{\mathrm{B}}$ | - | Allometric parameter of length-diameter relationship of branches. Important: 1.2-3 for trunks | 1.2-3 | McMahon 1973, Bertram 1989, Niklas 1995, West, Brown and Enquist 1999, van Gelder, Poorter and Sterck 2006, Banin et al. 2012 |
| $LP_{\mathrm{ratio}}$ | $cm^2\ cm^{-2}$ | Ratio between leaf area and pipe cross-sectional area. Important: PipeReuseFactor also plays a role here | 5000-50000 | Wright et al. 2006, Calvo-Alvarado, McDowell and Waring 2008, Patiño et al. 2012 |
| $m_{TB}$ | $g^{-1}\ a^{-1}$ | Parameter of biomass-based tree mortality rate | 0.01-0.05 | Model specific parameter, own estimates |
| $M_{TS}$ | - | Scaling exponent in biomass-based tree mortality rate, The metabolic theory predicts a scaling exponent of ¼, but significant deviations have been observed. | 0.1-0.4 | Brown et al. 2004, Muller-Landau et al. 2006a,b |
| $m_{\mathrm{Neigh}}$ | $a^{-1}$ | Trees affected by falling trees die with a probability of $m_{\mathrm{Neigh}}$ | 0.05-0.3 | Model specific parameter, own estimates |
| $R_{\mathrm{w}}$ | $gC\ g^{-1}\ d^{-1}$ | Respiration rate per gram of sapwood | $10^{-4}$-$10^{-6}$ | Penning De Vries 1975, Ryan et al. 1994, 1995, Vose and Ryan 2002, Sterck et al. 2005 |
| $n_{\mathrm{Seed}}$ | $ha^{-1}\ a^{-1}$ | Number of seedlings dispersed at each time step (per hectare) | 100-1000 | Model specific parameter, own estimates |
| $S_{\mathrm{F}}$ | - | Factor regulating the shortening of branches with their order | 0.7-0.9 | Bertram 1989, Perttunen et al. 1996, Perttunen, Sieva and Nikinmaa 1998 |
| $SI$ | - | Site index describing the relative quality of the forest patch | 0.5-1 | Model specific parameter, own estimates |
| $SLA$ | $cm^2\ g^{-1}$ | Specific leaf area, mean value for Amazonia ~90-110 | 50-200 | Poorter 1999, Rijkers et al. 2000, Kitajima, Mulkey and Wright 2005a, Wright et al. 2005, 2010, Rozendaal et al. 2006, Markesteijn et al. 2007, Domingues, Martinelli and Ehleringer 2007, Poorter et al. 2009, Malhi et al. 2009, Patiño et al. 2012, Niinemets, Keenan and Halllik 2015 |
| $\rho_{\mathrm{W}}$ | $g\ cm^{-3}$ | Wood density, mean value~0.63 for Amazonia | 0.4-0.8 | Baker et al. 2004, Patiño et al. 2009, Quesada et al. 2012, Iida et al. 2014 |
| $P_{RU}$ | - | Pipe-reuse factor | 0.5-0.9 | Mäkelä 1986, 2002, Perttunen et al. 1996, Sterck and Schieving 2007 |
| $t_{year}$ | d | Number of days per year suitable for photosynthesis | 180-360 | Model specific parameter, own estimates |
| $h_{sun}$ | h | Assumed number of sun hours per day | 6-10 | Model specific parameter, own estimates |
| $ST$ | - | Safety factor for trunk growth. | 0.3-0.5 | Sterck and Bongers 1998, van Gelder et al. 2006, Osunkoya et al. 2007 |

Feldpausch, T. R., L. Banin, O. L. Phillips, T. R. Baker, S. L. Lewis, C. A. Quesada, K. Affum-Baffoe, E. J. M. M. Arets, N. J. Berry, M. Bird, E. S. Brondizio, P. de Camargo, J. Chave, G. Djagbletey, T. F. Domingues, M. Drescher, P. M. Fearnside, M. B. França, N. M. Fyllas, G. Lopez-Gonzalez, A. Hladik, N. Higuchi, M. O. Hunter, Y. Iida, K. A. Salim, A. R. Kassim, M. Keller, J. Kemp, D. A. King, J. C. Lovett, B. S. Marimon, B. H. Marimon-

Junior, E. Lenza, A. R. Marshall, D. J. Metcalfe, E. T. A. Mitchard, E. F. Moran, B. W. Nelson, R. Nilus, E. M. Nogueira, M. Palace, S. Patiño, K. S.-H. Peh, M. T. Raventos, J. M. Reitsma, G. Saiz, F. Schrodt, B. Sonké, H. E. Taedoumg, S. Tan, L. White, H. Wöll, and J. Lloyd. 2011. Height-diameter allometry of tropical forest trees. Biogeosciences 8:1081–1106.

Feldpausch, T. R., J. Lloyd, S. L. Lewis, R. J. W. Brienen, E. Gloor, A. Montagudo Mendoza, G. Lopez-Gonzalez, and L. Banin. 2012. Tree height integrated into pan-tropical forest biomass estimates. Biogeosciences 9:1–23.

Franklin, O., J. Johansson, R. C. Dewar, U. Dieckmann, R. E. McMurtrie, A. Brännström, and R. Dybzinski. 2012. Modeling carbon allocation in trees: a search for principles. Tree Physiology 32:648–66.

Galbraith, D., Y. Malhi, K. Affum-Baffoe, A. D. A. Castanho, C. E. Doughty, R. A. Fisher, S. L. Lewis, K. S.-H. Peh, O. L. Phillips, C. A. Quesada, B. Sonké, and J. Lloyd. 2013. Residence times of woody biomass in tropical forests. Plant Ecology & Diversity 6:139–157.

van Gelder, H. A., L. Poorter, and F. J. Sterck. 2006. Wood mechanics, allometry, and life-history variation in a tropical rain forest tree community. New Phytologist 171:367–378.

Girardin, C. A. J., Y. Malhi, L. E. O. C. Aragão, M. Mamani, W. Huaraca Huasco, L. Durand, K. J. Feeley, J. Rapp, J. E. Silva-Espejo, M. Silman, N. Salinas, and R. J. Whittaker. 2010. Net primary productivity allocation and cycling of carbon along a tropical forest elevational transect in the Peruvian Andes. Global Change Biology 16:3176–3192.

Grimm, V., U. Berger, F. Bastiansen, S. Eliassen, V. Ginot, J. Giske, J. Goss-Custard, T. Grand, S. K. Heinz, G. Huse, A. Huth, J. U. Jepsen, C. Jørgensen, W. M. Mooij, B. Müller, G. Pe’er, C. Piou, S. F. Railsback, A. M. Robbins, M. M. Robbins, E. Rossmanith, N. Rüger, E. Strand, S. Souissi, R. A. Stillman, R. Vabo, U. Visser, and D. L. DeAngelis. 2006. A standard protocol for describing individual-based and agent-based models. Ecological Modelling 198:115–126.

Grimm, V., U. Berger, D. L. DeAngelis, J. G. Polhill, J. Giske, and S. F. Railsback. 2010. The ODD protocol: A review and first update. Ecological Modelling 221:2760–2768.

Hallé, F., R. A. A. Oldeman, and P. B. Tomlinson. 1978. Tropical trees and forests: an architectural analysis. Springer, Berlin, Heidelberg, New York.

Helmer, E. H., and M. A. Lefsky. 2006. Forest canopy heights in Amazon river basin forests as estimated with the Geoscience Laser Altimeter System (GLAS). USDA Forest Service Proceedings RMRS-P-42CD.

Houghton, R. A., K. T. Lawrence, J. L. Hackler, and S. Brown. 2001. The spatial distribution of forest biomass in the Brazilian Amazon: a comparison of estimates. Global Change Biology 7:731–746.

Huth, A., and T. Ditzer. 2000. Simulation of the growth of a lowland Dipterocarp rain forest with FORMIX3. Ecological Modelling 134:1–25.

Iida, Y., L. Poorter, F. Sterck, A. R. Kassim, M. D. Potts, T. Kubo, and T. S. Kohyama. 2014. Linking size-dependent growth and mortality with architectural traits across 145 co-occurring tropical tree species. Ecology 95:353–63.

King, D. A. 1997. Branch growth and biomass allocation in *Abies amabilis* saplings in contrasting light environments. Tree Physiology 17:251–258.

King, D. A. 1991. Correlations between biomass allocation, relative growth rate and light environment in tropical forest saplings. Functional Ecology 5:485–492.

King, D. A., E. G. Leigh, R. Condit, R. B. Foster, and S. P. Hubbell. 1997. Relationships between branch spacing, growth rate and light in tropical forest saplings. Functional Ecology 11:627–635.

Kitajima, K., S. S. Mulkey, and S. J. Wright. 2005. Variation in crown light utilization characteristics among tropical canopy trees. Annals of Botany 95:535–547.

Kniemeyer, O. G. M. 2008. Design and implementation of a graph grammar based language for functional-structural plant modelling. PhD Thesis, University of Cottbus.

de Kroon, H., H. Huber, J. F. Stuefer, and J. M. van Groenendael. 2005. A modular concept of phenotypic plasticity in plants. New Phytologist 166:73–82.

Lacointe, A. 2000. Carbon allocation among tree organs: A review of basic processes and representation in functional-structural tree models. Annals of Forest Science 57:521–533.

Laurance, S. G. W., W. F. Laurance, H. E. M. Nascimento, A. Andrade, P. M. Fearnside, E. R. G. Rebello, and R. Condit. 2009. Long-term variation in Amazon forest dynamics. Journal of Vegetation Science 20:323–333.

Lewis, S. L., O. L. Phillips, T. R. Baker, J. Lloyd, Y. Malhi, S. Almeida, N. Higuchi, W. F. Laurance, D. A. Neill, J. N. M. Silva, J. Terborgh, A. Torres Lezama, R. Vasquez Martinez, S. Brown, J. Chave, C. Kuebler, P. Nunez Vargas, and B. Vinceti. 2004. Concerted changes in tropical forest structure and dynamics: evidence from 50 South American long-term plots. Philosophical Transactions of the Royal Society B: Biological Sciences 359:421–436.

Lewis, S. L., O. L. Phillips, D. Sheil, B. Vinceti, T. R. Baker, S. Brown, A. W. Graham, N. Higuchi, D. W. Hilbert, W. F. Laurance, J. Lejoly, Y. Malhi, A. Monteagudo, P. Nunez Vargas, B. Sonke, J. W. Terborgh, and R. Vasquez Martinez. 2004. Tropical forest tree mortality, recruitment and turnover rates: calculation, interpretation and comparison when census intervals vary. Journal of Ecology 92:929–944.

Lieberman, D., M. Lieberman, R. Peralta, and G. S. Hartshorn. 1996. Tropical forest structure and composition on a large-scale altitudinal gradient in Costa Rica. Journal of Ecology 84:137–152.

Lindenmayer, A. 1968a. Mathematical models for cellular interactions in development. II. Simple and branching filaments with two-sided inputs. Journal of Theoretical Biology 18:300–315.

Lindenmayer, A. 1968b. Mathematical models for cellular interactions in development. I. Filaments with one-sided inputs. Journal of Theoretical Biology 18:280–299.

Mäkelä, A. 1986. Implications of the pipe model theory on dry matter partitioning and height growth in trees. Journal of Theoretical Biology 123:103–120.

Mäkelä, A. 2002. Derivation of stem taper from the pipe theory in a carbon balance framework. Tree Physiology 22:891–905.

Mäkelä, A. 2012. On guiding principles for carbon allocation in eco-physiological growth models. Tree Physiology 32:644–647.

Malhi, Y., L. E. O. C. Aragão, D. B. Metcalfe, R. Paiva, C. A. Quesada, S. Almeida, L. Anderson, P. Brando, J. Q. Chambers, A. C. L. Da Costa, L. R. Hutyra, P. Oliveira, S. Patiño, E. H. Pyle, A. L. Robertson, and L. M. Teixeira. 2009. Comprehensive assessment of carbon productivity, allocation and storage in three Amazonian forests. Global Change Biology 15:1255–1274.

Malhi, Y., T. R. Baker, O. L. Phillips, S. Almeida, E. Alvarez, L. Arroyo, J. Chave, C. I. Czimczik, A. Di Fiore, N. Higuchi, T. J. Killeen, S. G. Laurance, W. F. Laurance, S. L. Lewis, L. M. M. Montoya, A. Monteagudo, D. A. Neill, P. N. Vargas, S. Patiño, N. C. A. Pitman, C. A. Quesada, R. Salomao, J. N. M. Silva, A. T. Lezama, R. V. Martinez, J. Terborgh, B. Vinceti, and J. Lloyd. 2004. The above-ground coarse wood productivity of 104 Neotropical forest plots. Global Change Biology 10:563–591.

Malhi, Y., C. E. Doughty, G. R. Goldsmith, D. B. Metcalfe, C. A. J. Girardin, T. R. Marthews, J. del Aguila-Pasquel, L. E. O. C. Aragão, A. Araujo-Murakami, P. Brando, A. C. L. da Costa, J. E. Silva-Espejo, F. Farfán Amézquita, D. R. Galbraith, C. A. Quesada, W. Rocha, N. Salinas-Revilla, D. Silvério, P. Meir, and O. L. Phillips. 2015. The linkages between photosynthesis, productivity, growth and biomass in lowland Amazonian forests. Global Change Biology 21:2283–2295.

Malhi, Y., C. Doughty, and D. Galbraith. 2011. The allocation of ecosystem net primary productivity in tropical forests. Philosophical Transactions of the Royal Society B: Biological Sciences 366:3225–45.

Malhi, Y., F. Farfán Amézquita, C. E. Doughty, J. E. Silva-Espejo, C. A. J. Girardin, D. B. Metcalfe, L. E. O. C. Aragão, L. P. Huaraca-Quispe, I. Alzamora-Taype, L. Eguiluz-Mora, T. R. Marthews, K. Halladay, C. A. Quesada, A. L. Robertson, J. B. Fisher, J. Zaragoza-Castells, C. M. Rojas-Villagra, Y. Pelaez-Tapia, N. Salinas, P. Meir, and O. L. Phillips. 2013. The productivity, metabolism and carbon cycle of two lowland tropical forest plots in south-western Amazonia, Peru. Plant Ecology & Diversity 7:1–21.

Malhi, Y., D. Wood, T. R. Baker, J. Wright, O. L. Phillips, T. Cochrane, P. Meir, J. Chave, S. Almeida, L. Arroyo, N. Higuchi, T. J. Killeen, S. G. Laurance, W. F. Laurance, S. L. Lewis, A. Monteagudo, D. A. Neill, P. N. Vargas, N. C. A. Pitman, C. A. Quesada, R. Salomao, J. N. M. Silva, A. T. Lezama, J. Terborgh, R. V. Martinez, and B. Vinceti. 2006. The regional variation of aboveground live biomass in old-growth Amazonian forests. Global Change Biology 12:1107–1138.

Marino, G., M. Aqil, and B. Shipley. 2010. The leaf economics spectrum and the prediction of photosynthetic light-response curves. Functional Ecology 24:263–272.

Markesteijn, L., L. Poorter, and F. Bongers. 2007. Light-dependent leaf trait variation in 43 tropical dry forest tree species. American Journal of Botany 94:515–525.

Martin, A. R., and S. C. Thomas. 2011. A reassessment of carbon content in tropical trees. PloS one 6:e23533.

McCoy, M. W., and J. F. Gillooly. 2008. Predicting natural mortality rates of plants and animals. Ecology Letters 11:710–716.

McMahon, T. 1973. Size and shape in biology. Science 179:1201–1204.

Mitchard, E. T. A., T. R. Feldpausch, R. J. W. Brienen, G. Lopez-Gonzalez, A. Monteagudo, T. R. Baker, S. L. Lewis, J. Lloyd, C. A. Quesada, M. Gloor, H. ter Steege, P. Meir, E. Alvarez, A. Araujo-Murakami, L. E. O. C. Aragão, L. Arroyo, G. Aymard, O. Banki, D. Bonal, S. Brown, F. I. Brown, C. E. Cerón, V. Chama Moscoso, J. Chave, J. A. Comiskey, F. Cornejo, M. Corrales Medina, L. Da Costa, F. R. C. Costa, A. Di Fiore, T. F. Domingues, T. L. Erwin, T. Frederickson, N. Higuchi, E. N. Honorio Coronado, T. J. Killeen, W. F. Laurance, C. Levis, W. E. Magnusson, B. S. Marimon, B. H. Marimon Junior, I. Mendoza Polo, P. Mishra, M. T. Nascimento, D. Neill, M. P. Núñez Vargas, W. A. Palacios, A. Parada, G. Pardo Molina, M. Peña-Claros, N. Pitman, C. A. Peres, L. Poorter, A. Prieto, H. Ramirez-Angulo, Z. Restrepo Correa, A. Roopsind, K. H. Roucoux, A. Rudas, R. P. Salomão, J. Schietti, M. Silveira, P. F. de Souza, M. K. Steininger, J. Stropp, J. Terborgh, R. Thomas, M. Toledo, A. Torres-Lezama, T. R. van Andel, G. M. F. van der Heijden, I. C. G. Vieira, S. Vieira, E. Vilanova-Torre, V. A. Vos, O. Wang, C. E. Zartman, Y. Malhi, and O. L. Phillips. 2014. Markedly divergent estimates of Amazon forest carbon density from ground plots and satellites. Global Ecology and Biogeography 23:935–946.

Muller-Landau, H. C., R. S. Condit, J. Chave, S. C. Thomas, S. A. Bohlman, S. Bunyavejchewin, S. Davies, R. Foster, S. Gunatilleke, N. Gunatilleke, K. E. Harms, T. Hart, S. P. Hubbell, A. Itoh, A. R. Kassim, J. V. LaFrankie, H. S. Lee, E. Losos, J. R. Makana, T. Ohkubo, R. Sukumar, I. F. Sun, M. N. Nur Supardi, S. Tan, J. Thompson, R. Valencia, G. V. Muñoz, C. Wills, T. Yamakura, G. Chuyong, H. S. Dattaraja, S. Esufali, P. Hall, C. Hernandez, D. Kenfack, S. Kiratiprayoon, H. S. Suresh, D. Thomas, M. I. Vallejo, and P. Ashton. 2006a. Testing metabolic ecology theory for allometric scaling of tree size, growth and mortality in tropical forests. Ecology Letters 9:575–588.

Muller-Landau, H. C., R. S. Condit, K. E. Harms, C. O. Marks, S. C. Thomas, S. Bunyavejchewin, G. Chuyong, L. Co, S. Davies, R. Foster, S. Gunatilleke, N. Gunatilleke, T. Hart, S. P. Hubbell, A. Itoh, A. R. Kassim, D. Kenfack, J. V. LaFrankie, D. Lagunzad, H. S. Lee, E. Losos, J.-R. Makana, T. Ohkubo, C. Samper, R. Sukumar, I.-F. Sun, M. N. Nur Supardi, S. Tan, D. Thomas, J. Thompson, R. Valencia, M. I. Vallejo, G. V. Muñoz, T. Yamakura, J. K. Zimmerman, H. S. Dattaraja, S. Esufali, P. Hall, F. He, C. Hernandez, S. Kiratiprayoon, H. S. Suresh, C. Wills, and P. Ashton. 2006b. Comparing tropical forest tree size distributions with the predictions of metabolic ecology and equilibrium models. Ecology letters 9:589–602.

Myneni, R. B., W. Yang, R. R. Nemani, A. R. Huete, R. E. Dickinson, Y. Knyazikhin, K. Didan, R. Fu, R. I. Negrón Juárez, S. S. Saatchi, H. Hashimoto, K. Ichii, N. V. Shabanov, B. Tan, P. Ratana, J. L. Privette, J. T. Morisette, E. F. Vermote, D. P. Roy, R. E. Wolfe, M. A. Friedl, S. W. Running, P. Votava, N. El-Saleous, S. Devadiga, Y. Su, and V. V. Salomonson. 2007. Large seasonal swings in leaf area of Amazon rainforests. Proceedings of the National Academy of Sciences of the United States of America 104:4820–4823.

Nicolini, E., Y. Caraglio, R. Pelissier, C. Leroy, and J.-C. Roggy. 2003. Epicormic branches: a growth indicator for the tropical forest tree, *Dicorynia guianensis* Amshoff (Caesalpiniaceae). Annals of Botany 92:97–105.

Niinemets, Ü., T. F. Keenan, and L. Halllik. 2015. A worldwide analysis of within-canopy variations in leaf structural, chemical and physiological traits across plant functional types. New Phytologist 205:973–993.

Niklas, K. J. 1995. Size-dependent allometry of tree height, diameter and trunk-taper. Annals of Botany 75:217–227

Osunkoya, O. O., K. Omar-Ali, N. Amit, J. Dayan, D. S. Daud, and T. K. Sheng. 2007. Comparative height crown allometry and mechanical design in 22 tree species of Kuala Belalong rainforest, Brunei, Borneo. American Journal of Botany 94:1951–62.

Patiño, S., N. M. Fyllas, T. R. Baker, R. Paiva, C. A. Quesada, A. J. B. Santos, M. Schwarz, H. Ter Steege, O. L. Phillips, and J. Lloyd. 2012. Coordination of physiological and structural traits in Amazon forest trees. Biogeosciences 9:775–801.

Patiño, S., J. Lloyd, R. Paiva, T. R. Baker, C. A. Quesada, L. M. Mercado, J. Schmerler, M. Schwarz, A. J. B. Santos, A. Aguilar, C. I. Czimczik, J. Gallo, V. Horna, E. J. Hoyos, E. M. Jimenez, W. Palomino, J. Peacock, A. Peña-Cruz, C. Sarmiento, A. Sota, J. D. Turriago, B. Villanueva, P. Vitzthum, E. Alvarez, L. Arroyo, C. Baraloto, D. Bonal, J. Chave, A. C. L. Costa, R. Herrera, N. Higuchi, T. Killeen, E. Leal, F. Luizão, P. Meir, A. Monteagudo, D. Neil, P. Núñez-Vargas, M. C. Peñuela, N. Pitman, N. Priante Filho, A. Prieto, S. N. Panfil, A. Rudas, R. Salomão, N. Silva, M. Silveira, S. Soares de Almeida, A. Torres-Lezama, R. Vásquez-Martínez, I. Vieira, Y. Malhi, and O. L. Phillips. 2009. Branch xylem density variations across the Amazon Basin. Biogeosciences 6:545–568.

Penning De Vries, F. W. T. 1975. The cost of maintenance processes in plant cells. Annals of Botany 39:77–92.

Perttunen, J., R. Sieva, and E. Nikinmaa. 1998. LIGNUM : a model combining the structure and the functioning of trees. Ecological Modelling 108:189–198.

Perttunen, J., R. Sievänen, E. Nikinmaa, H. Salminen, H. Saarenmaa, and J. Väkevä. 1996. LIGNUM: a tree model based on simple structural units. Annals of Botany 77:87–98.

Phillips, O. L., P. Hall, A. H. Gentry, S. A. Sawyer, and R. Vásquez. 1994. Dynamics and species richness of tropical rain forests. Proceedings of the National Academy of Sciences of America 91:2805-2809

Phillips, O. L. 1996. Long-term environmental change in tropical forests: increasing tree turnover. Environmental Conservation 23:235.

Phillips, O. L., T. R. Baker, L. Arroyo, N. Higuchi, T. J. Killeen, W. F. Laurance, S. L. Lewis, J. Lloyd, Y. Malhi, A. Monteagudo, D. A. Neill, P. Nunez Vargas, J. N. M. Silva, J. Terborgh, R. Vasquez Martinez, M. Alexiades, S. Almeida, S. Brown, J. Chave, J. A. Comiskey, C. I. Czimczik, A. Di Fiore, T. Erwin, C. Kuebler, S. G. Laurance, H. E. M. Nascimento, J. Olivier, W. Palacios, S. Patino, N. C. A. Pitman, C. A. Quesada, M. Saldias, A. Torres Lezama, and B. Vinceti. 2004. Pattern and process in Amazon tree turnover, 1976-2001. Philosophical Transactions of the Royal Society B: Biological Sciences 359:381–407.

Phillips, O. L., N. Higuchi, S. Vieira, T. R. Baker, K. J. Chao, and S. L. Lewis. 2009. Changes in Amazonian forest biomass, dynamics, and composition, 1980-2002. Geophysical Monograph Series 186:373–387.

Pons, T. L., and H. Poorter. 2014. The effect of irradiance on the carbon balance and tissue characteristics of five herbaceous species differing in shade-tolerance. Frontiers in Plant Science 5:12.

Poorter, H., and R. De Jong. 1999. A comparison of specific leaf area, chemical composition and leaf construction costs of field plants from 15 habitats differing in productivity. New Phytologist 143:163–176.

Poorter, H., Ü. Niinemets, L. Poorter, I. J. Wright, and R. Villar. 2009. Causes and consequences of variation in leaf mass per area (LMA): a meta-analysis. New Phytologist 182:565–588.

Poorter, L. 1999. Growth responses of 15 rain-forest tree species to a light gradient: the relative importance of morphological and physiological traits. Functional Ecology 13:396–410.

Poorter, L., F. Bongers, F. J. Sterck, and H. Wöll. 2011. Architecture of 53 rain forest tree species differing in adult stature and shade tolerance. Ecology 84:602–608.

Quesada, C. A., O. L. Phillips, M. Schwarz, C. I. Czimczik, T. R. Baker, S. Patiño, N. M. Fyllas, M. G. Hodnett, R. Herrera, S. Almeida, E. Alvarez Dávila, A. Arneth, L. Arroyo, K. J. Chao, N. Dezzeo, T. Erwin, A. di Fiore, N. Higuchi, E. Honorio Coronado, E. M. Jimenez, T. Killeen, A. T. Lezama, G. Lloyd, G. López-González, F. J. Luizão, Y. Malhi,

A. Monteagudo, D. A. Neill, P. Núñez Vargas, R. Paiva, J. Peacock, M. C. Peñuela, A. Peña Cruz, N. Pitman, N. Priante Filho, A. Prieto, H. Ramírez, A. Rudas, R. Salomão, A. J. B. Santos, J. Schmerler, N. Silva, M. Silveira, R. Vásquez, I. Vieira, J. Terborgh, and J. Lloyd. 2012. Basin-wide variations in Amazon forest structure and function are mediated by both soils and climate. Biogeosciences 9:2203–2246.

Rijkers, T., T. L. Pons, and F. Bongers. 2000. The effect of tree height and light availability on photosynthetic leaf traits of four neotropical species differing in shade tolerance. Functional Ecology 14:77–86.

Rozendaal, D. M. A., V. H. Hurtado, and L. Poorter. 2006. Plasticity in leaf traits of 38 tropical tree species in response to light; relationships with light demand and adult stature. Functional Ecology 20:207–216.

Ryan, M. G., S. T. Gower, R. M. Hubbard, R. H. Waring, H. L. Gholz, W. P. Cropper, and S. W. Running. 1995. Woody tissue maintenance respiration of four conifers in contrasting climates. Oecologia 101:133–140.

Ryan, M. G., R. M. Hubbard, D. A. Clark, and R. L. Sanford. 1994. Woody-tissue respiration for *Simarouba amara* and *Minquartia guianensis*, two tropical wet forest trees with different growth habits. Oecologia 100:213–220.

Sawada, Y., R. Suwa, K. Jindo, T. Endo, K. Oki, H. Sawada, E. Arai, Y. E. Shimabukuro, C. H. S. Celes, M. A. A. Campos, F. G. Higuchi, A. J. N. Lima, N. Higuchi, T. Kajimoto, and M. Ishizuka. 2015. A new 500-m resolution map of canopy height for Amazon forest using spaceborne LiDAR and cloud-free MODIS imagery. International Journal of Applied Earth Observation and Geoinformation 43:92–101.

Seyoum, F., M. Fetene, S. Strobl, and E. Beck. 2014. Daily and seasonal courses of gas exchange and niche partitioning among coexisting tree species in a tropical montane forest. Flora 209:191–200.

Shinozaki, K., K. Yoda, K. Hozumi, and T. Kira. 1964. A quantitative analysis of plant form-the pipe model theory I. Basic analyses. Japanese Journal of Ecology 14:97–105.

Simard, M., N. Pinto, J. B. Fisher, and A. Baccini. 2011. Mapping forest canopy height globally with spaceborne lidar. Journal of Geophysical Research 116: G04021.

Slik, J. W. F., G. Paoli, K. McGuire, I. Amaral, J. Barroso, M. Bastian, L. Blanc, F. Bongers, P. Boundja, C. Clark, M. Collins, G. Dauby, Y. Ding, J.-L. Doucet, E. Eler, L. Ferreira, O. Forshed, G. Fredriksson, J.-F. Gillet, D. Harris, M. Leal, Y. Laumonier, Y. Malhi, A. Mansor, E. Martin, K. Miyamoto, A. Araujo-Murakami, H. Nagamasu, R. Nilus, E. Nurtjahya, Á. Oliveira, O. Onrizal, A. Parada-Gutierrez, A. Permana, L. Poorter, J. Poulsen, H. Ramirez-Angulo, J. Reitsma, F. Rovero, A. Rozak, D. Sheil, J. Silva-Espejo, M. Silveira, W. Spironelo, H. ter Steege, T. Stevart, G. E. Navarro-Aguilar, T. Sunderland, E. Suzuki, J. Tang, I. Theilade, G. van der Heijden, J. van Valkenburg, T. Van Do, E. Vilanova, V. Vos, S. Wich, H. Wöll, T. Yoneda, R. Zang, M.-G. Zhang, and N. Zweifel. 2013. Large trees drive forest aboveground biomass variation in moist lowland forests across the tropics. Global Ecology and Biogeography 22:1261–1271.

Sprugel, D. G., T. M. Hinckley, and W. Schaap. 1991. The theory and practice of branch autonomy. Annual Review of Ecology and Systematics 22:309–334.

Sterck, F. J., and F. Bongers. 1998. Ontogenetic changes in size, allometry, and mechanical design of tropical rain forest trees. American Journal of Botany 85:266–272.

Sterck, F. J., L. Markesteijn, F. Schieving, and L. Poorter. 2011. Functional traits determine trade-offs and niches in a tropical forest community. Proceedings of the National Academy of Sciences 108:20627–20632.

Sterck, F. J., and F. Schieving. 2007. 3-D growth patterns of trees: effects of carbon economy, meristem activity, and selection. Ecological Monographs 77:405–420.

Sterck, F. J., F. Schieving, A. Lemmens, and T. L. Pons. 2005. Performance of trees in forest canopies: explorations with a bottom-up functional-structural plant growth model. New Phytologist 166:827–843.

Stoll, P., and B. Schmid. 1998. Plant foraging and dynamic competition between branches of *Pinus sylvestris* in contrasting light environments. Journal of Ecology 86:934–945.

Valladares, F., M. T. Allen, and R. W. Pearcy. 1997. Photosynthetic responses to dynamic light under field conditions in six tropical rainforest shrubs occuring along a light gradient. Oecologia 111:505–514.

Vose, J. M., and M. G. Ryan. 2002. Seasonal respiration of foliage, fine roots, and woody tissues in relation to growth, tissue N, and photosynthesis. Global Change Biology 8:182–193.

West, G. B., J. H. Brown, and B. J. Enquist. 1999. A general model for the structure and allometry of plant vascular systems. Nature 400:664–667.

Wilson, B. F. 2000. Apical control of branch growth and angle in woody plants. American Journal of Botany 87:601–607.

Wright, I. J., D. S. Falster, M. Pickup, and M. Westoby. 2006. Cross-species patterns in the coordination between leaf and stem traits, and their implications for plant hydraulics. Physiologia Plantarum 127:445–456.

Wright, I. J., P. B. Reich, J. H. C. Cornelissen, D. S. Falster, E. Garnier, K. Hikosaka, B. B. Lamont, W. Lee, J. Oleksyn, N. Osada, H. Poorter, R. Villar, D. I. Warton, and M. Westoby. 2005. Assessing the generality of global leaf trait relationships. New Phytologist 166:485–496.

Wright, I. J., P. B. Reich, M. Westoby, D. D. Ackerly, Z. Baruch, F. Bongers, J. Cavender-Bares, T. Chapin, J. H. C. Cornelissen, M. Diemer, J. Flexas, E. Garnier, P. K. Groom, J. Gulias, K. Hikosaka, B. B. Lamont, T. Lee, W. Lee, C. Lusk, J. J. Midgley, M.-L. Navas, U. Niinemets, J. Oleksyn, N. Osada, H. Poorter, P. Poot, L. Prior, V. I. Pyankov, C. Roumet, S. C. Thomas, M. G. Tjoelker, E. J. Veneklaas, and R. Villar. 2004. The worldwide leaf economics spectrum. Nature 428:821–827.

Wright, S. J., K. Kitajima, N. J. B. Kraft, P. B. Reich, I. J. Wright, D. E. Bunker, R. Condit, J. W. Dalling, S. J. Davies, S. Díaz, B. M. J. Engelbrecht, K. E. Harms, S. P. Hubbell, C. O. Marks, M. C. Ruiz-Jaen, C. M. Salvador, and A. E. Zanne. 2010. Functional traits and the growth-mortality trade-off in tropical trees. Ecology 91:3664–3674.
